# Rewiring Fibroblast–Muscle Axis Drives Progressive Pathology in Bethlem Myopathy

**DOI:** 10.64898/2026.05.14.725126

**Authors:** Shivashakthi Shivaraman, Laurent Gilquin, Frederic Sohm, Rawan Fareh, Laurence Legeais-Mallet, Antonella Forlino, Emilie Dambroise, Sandrine Bretaud, Florence Ruggiero

## Abstract

Collagen VI–related myopathies, including Bethlem myopathy (BM), are progressive muscle disorders, but the mechanisms driving age-dependent disease progression remain poorly understood. Here, we used a zebrafish BM model carrying an exon-skipping mutation that generates a shorter collagen VI α1 chain and disrupts supramolecular assembly, recapitulating key features of the human disease. We further demonstrated that this model reproduces disease progression, with worsening muscle wasting, increased myofiber size variability, and age-associated skeletal deformities consistent with secondary consequences of muscle dysfunction rather than intrinsic bone defects. Single-nucleus RNA sequencing of trunk skeletal muscle revealed an early shift in cellular composition, with reduced myonuclei and increased fibroblast abundance, indicative of disease-associated aging. Myonuclei activated stress and quality control pathways, including autophagy and mitophagy, along with metabolic rewiring. In contrast, fibroblasts displayed early translational activation followed by progressive proteostatic and endoplasmic reticulum stress. At later stages, fibroblasts adopted a pro-fibrotic state, driving extracellular matrix remodeling and enhanced muscle–fibroblast communication. Consistently, analyses at the protein level confirmed early intracellular retention of the mutant protein, along with increased extracellular matrix deposition and fibrotic tissue formation in BM muscle. Among the three tested drugs targeting ER-stress and protein degradation, only TUDCA significantly ameliorated collagen VI deposition in the extracellular space in larvae. These findings identify fibroblasts as key drivers of disease progression and potential therapeutic targets.

## INTRODUCTION

Skeletal muscle function depends not only on the contractile apparatus of myofibers, but also on the surrounding extracellular matrix (ECM). This specialized muscle ECM, or myomatrix, forms a dynamic scaffold that supports force transmission, protects the sarcolemma during contraction, maintains the satellite cell niche, and coordinates vascular and neural integration. Beyond its structural role, the myomatrix acts as a biochemically active microenvironment that undergoes continuous remodeling during ageing, regeneration, and disease (Kragstrup et al., 2011; Lamandé & Bateman, 2018). Central to myomatrix are collagens, a superfamily of ECM proteins whose dysfunction is sufficient to drive connective tissue diseases called collagenopathies (Salamito et al., 2021).

Among myomatrix major components, collagen VI (ColVI) occupies a central position at the interface between the basement membrane and interstitial matrix. Pathogenic variants in the three genes encoding the canonical ColVI α-chains, *COL6A1*, *COL6A2* and *COL6A3*, cause collagen VI-related myopathies (COL6-RM), a spectrum of inherited neuromuscular disorders ranging from severe Ullrich Congenital Muscular Dystrophy (UCMD) to the milder Bethlem myopathy (BM) (Di Martino et al., 2023a; Lampe & Bushby, 2005). Although COL6-RM has long been viewed as a structural ECM disorder, accumulating evidence indicates that ColVI deficiency also disrupts autophagy, mitochondrial homeostasis, calcium handling and cell survival, revealing a broader dysfunction of the muscle microenvironment (Idoux et al., 2025; Irwin et al., 2003; Mohassel et al., 2023).

Being primarily synthesized by interstitial fibroblasts in skeletal muscle, ColVI, among other collagens (Guillon et al., 2025), is also produced by muscle stem cells (a.k.a. satellite cells) where it contributes to stem cell maintenance and regeneration (Urciuolo et al., 2013). Through interactions with ECM components, basement membrane proteins and transmembrane receptors, ColVI acts as both a mechanical linker and a signaling hub that transduces extracellular cues into intracellular responses regulating muscle homeostasis (Tonelotto et al., 2022). These observations support a non–cell-autonomous disease mechanism in which defective fibroblast–myofiber communication contributes to disease progression.

Clinically, BM is characterized by slowly progressive muscle weakness, contractures and skeletal abnormalities that worsen with age and resemble features of premature ageing (Bönnemann, 2011). While skeletal muscle pathology has been extensively investigated, the contribution of muscle-bone crosstalk to disease progression remains poorly understood. Consistent with this possibility, *Col6a1^-/-^* and *Col6a2^-/-^* mouse models display reduced trabecular bone mass and altered bone remodeling (Christensen et al., 2012; Pham et al., 2020), although whether these defects arise intrinsically within bone or secondarily to muscle dysfunction remains unresolved.

Our zebrafish BM model reproducing an exon skipping mutation (referred to as *col6a1^Δexon14^* KI line), frequently found in patients, faithfully reproduces key features of human disease, including muscle intracellular defects such as altered mitochondria and dilated sarcoplasmic reticulum, alongside progressive muscle weakness (Radev et al., 2015). Recent work further demonstrated that ColVI deficiency disrupts RyR/DHPR organization and calcium homeostasis, establishing a mechanistic link between extracellular matrix defects and intracellular muscle dysfunction (Idoux et al., 2025). However, the mechanisms driving disease progression remain unclear.

Here, we show that progressive muscle weakness in BM zebrafish leads to secondary skeletal deformities without intrinsic defects in mineralization or bone remodeling, supporting a reduced mechanical loading mechanism. Using single-nucleus RNA sequencing and immunofluorescence analyses of skeletal muscle across disease progression, we identify a profound disruption of fibroblast–myofiber communication in aged BM muscle, characterized by exacerbated ECM production and a striking increase in myofibers-fibroblast interactions. Together, our findings redefine COL6-RM as a disorder of the muscle microenvironment in which fibroblast dysfunction, ECM disorganization and impaired muscle-fibroblast coupling are central drivers of disease progression.

## RESULTS

### BM fish recapitulate progressive muscle wasting and age-associated skeletal alterations

To address muscle phenotype progression in BM fish, we first assessed overall muscle condition using the weight-to-length ratio. A significant reduction in muscle mass was evident in BM fish at 1 year compared to age-matched WT controls (Figure 1A), independent of sex (Figure S1A). Based on collagen VI localization and expression dynamics (Figure 1B and C; and Idoux et al, 2025), we selected 3 months (juveniles, with intracellular collagen VI aggregates still present) and 1 year (adults, with extracellular collagen VI staining absent) as representative early and late disease stages. Consistent with human pathology, WGA staining to outline muscle fibers revealed pronounced fiber size variability in BM fish at both 3 months and 1 year compared to WT (Figure 1D. This heterogeneity in myofiber size indicates ongoing dystrophic remodeling and impaired muscle maintenance over time.

**Figure 1:**
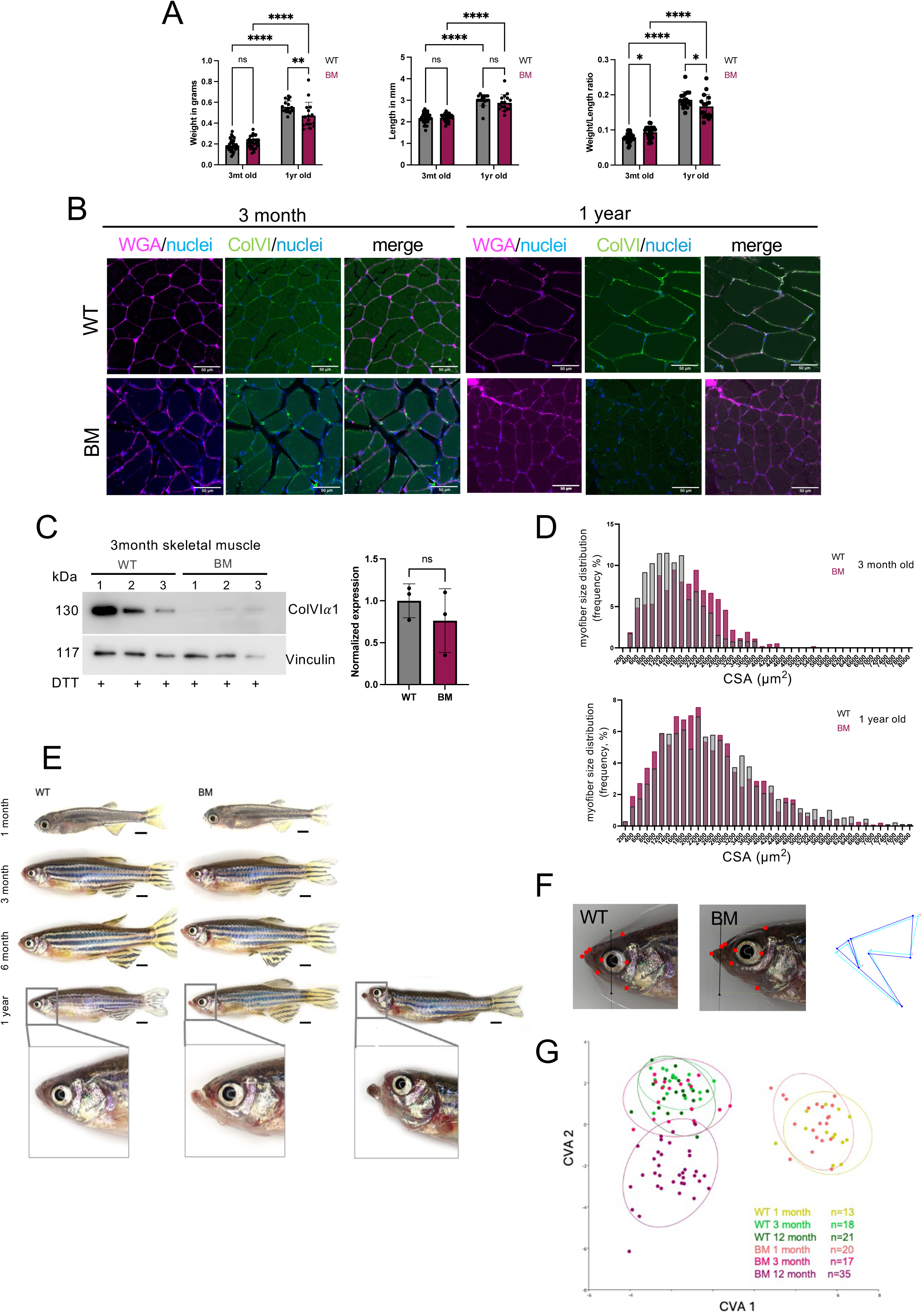
Progressive muscle (A-D) and bone (E-G) phenotype of BM fish –. **A** Measurements of skeletal muscle mass loss (weight, standard length and ratio) at 3 months and 1 year-old WT and BM fish. Sample sizes: 3mt WT n=24, BM n=24; 1yr WT n=19, BM n=18. **B**. Immunofluorescent staining of collagen VI (guinea pig colVI antibodies, in green) and WVA (magenta) in frozen cross sections of the BM and WT trunk at 3 months and 1 year-old. Nuceli are in blue. Scale bars = 50 µm. **C**. Representative membrane of Western-blot analysis of the presence of collagen VIα1 band in total protein extracts of WT or BM trunk at 3 months (DTT, reduced condition), right panel: densitometric analysis of band intensities in BM and WT at 3 month (n= 3). **D**. Myofiber size distribution (Frequency in percentage of fibers) in WT and BM cross sections trunk skeletal muscle at 3 month (upper panel) and 1 year (lower panel). CSA, cross section areas. 3 months old BM and WT: n_fish=3, n_field=12, n_myofibers = 865 (WT and 704 BM); 1-year old BM and WT: n_fish=5, n_field=9, n_myofibers = 1291 (WT) and 1255 (BM). **E**. Representative images of WT and BM zebrafish at 1, 3, 6, and 12 months of age (lateral views). Boxed areas in 1-year fish images correspond to the zoomed images of the head (below). Scale bars = 1 mm. **F.** Reference landmarks on the WT and BM fish head and the corresponding shape variations between WT (light blue) and BM (indigo). **G**. Canonical Variate Analysis (CVA) of first two morphometric variates of WT and BM fish at 1 month, 3 month and 1-year old. WT 1 month n=13, WT 3 months n=18, WT 12 months n=21; BM 1 month n=20, BM 3 months n=17, BM 12 months n=35. Statistical comparisons are performed using morphometric landmark analysis in TpsDig and MorphoJ. Each point represents one individual; ellipses indicate 95% confidence regions. Pillais Trace test, ****p<0.0001.

BM in humans is also associated with secondary skeletal deformities, generally attributed to muscle weakness and contractures rather than primary bone defects. Zebrafish similarly exhibit age-related skeletal changes, including spinal curvature and vertebral abnormalities (Hayes et al., 2013; Kishi et al., 2008). In BM fish, we observed progressive abnormalities in jaw and spine morphology with age, whereas WT fish did not display such alterations (Figure 2E). Notably, jaw deformities have not been previously reported as an age-associated feature in zebrafish, suggesting a disease-specific effect. Multivariate morphometric and canonical variate analysis (CVA) (Figure 1F) of head shape revealed a clear separation of 1-year-old BM fish from all other groups, indicating a distinct age-associated bone phenotype linked to disease progression (Figure 1F and G).

**Figure 2:**
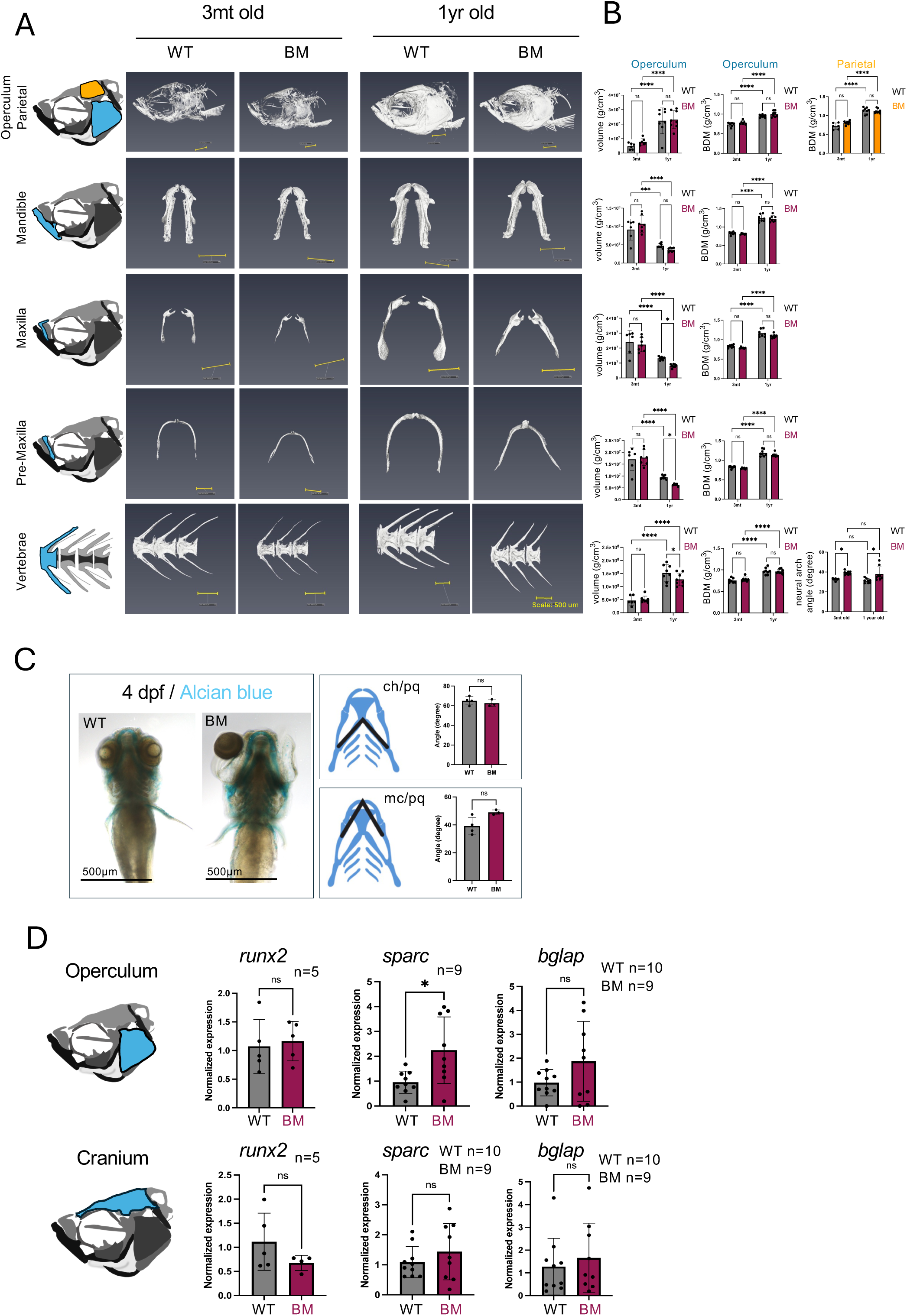
Progressive bone phenotype is not due to a late consequence of defective skeletal development. **A.** Representative micro-CT reconstructions of WT and BM zebrafish craniofacial (right operculum and parietal, mandible, maxilla, pre-maxilla) and axial skeletal (caudal vertebrae) elements as indicated at 3 months and 1 year (Sample sizes: WT 3 months n=7, WT 1 year n=8; BM 3 months n=8, BM 1 year n=8). Scale bars = 500 µm. **B.** Bone volume and bone mineral density (BMD) of the skeletal elements shown in A. For the vertebrae, measurements of the neural arch angle are also shown. WT 3 months n=7, WT 1 year n=8; BM 3 months n=8, BM 1 year n=8. All statistical analysis were performed using two-way ANOVA. Data represent ± SEM; two-way ANOVA; ns = not significant, *p < 0.05, ***p < 0.001, ****p < 0.0001. **C.** Alcian blue staining of 4dpf WT and BM fish (left panel) and quantification of the angle of the ceratohyal (ch) and Meckel’s (mc) cartilage to palatoquadrate (pq) (right panel). Sample size: WT = 4, BM = 3. Data are mean ± SEM; unpaired t-test; ns = not significant. Statistical analysis was performed using the Mann-Whitney unpaired t-test. ns: no significance. Error bars are SEM. **D.** Quantitative RT-PCR of osteogenic marker genes, *runx2b*, *sparc*, and *bglap* in operculum and cranium dissected from WT and BM fish at 1 year. Expression was normalised to the housekeeping gene expression (*actb*). Statistical significance was assessed using an unpaired t-test. Data are mean ± SEM. ns: no significance; *p<0.05.

To determine whether these skeletal alterations (spine and jaw) reflected primary bone defects or secondary consequences of muscle dysfunction, we performed microCT-based analyses of bone mineral density (BMD) and bone volume (Figure 2A and B). No significant differences in BMD were observed between BM and WT fish in cranial or vertebral structures (Figure 2B). However, alterations in neural arch angle were detected in aged BM fish, consistent with reduced mechanical loading due to muscle weakness. Further volumetric analyses showed significant changes in craniofacial bone morphology in BM fish at 3 months compared to 1 year (Figure 2A and B). While similar age-dependent changes were also observed in WT fish, indicating that these effects reflect normal growth rather than disease-specific defects, we observed a significant reduction between BM and WT fish at 1 year for maxilla and pre-maxilla bones (Figure 2B). Together, these findings likely support the interpretation that skeletal alterations in BM fish are secondary to impaired muscle function rather than intrinsic bone abnormalities. To further exclude developmental bone defects, we performed skeletal staining staining at 4 dpf (Alcian blue) (Figure 2C, quantification) and 1 month (Alizarin red and Alcian blue) (Figure S2), which revealed no overt skeletal abnormalities in BM larvae and juveniles. In addition, expression levels of osteoblast and bone remodeling markers (*runx2*, *sparc*, and *bglap*) were assessed by RT-qPCR in WT and BM opercular (muscle-associated) and cranium (non-muscle-associated control) bones at 3 months and 1 year. No significant differences were observed between BM and WT fish, except for a modest increase in *sparc* expression in the operculum of BM fish (Figure 2D).

Collectively, these results demonstrate that BM fish recapitulate the progressive muscle wasting observed in human BM and develop secondary skeletal and craniofacial deformities over time. Importantly, these alterations occur in the absence of primary defects in bone development or mineralization, strongly supporting the conclusion that they arise as a consequence of progressive muscle weakness and reduced mechanical loading.

### snRNA-seq reveals early muscle stress responses and fibrotic remodeling in collagen VI–deficient BM fish

To investigate the molecular basis underlying the skeletal and muscular phenotypes observed during disease progression, we performed single-nucleus RNA sequencing (snRNA-seq) of trunk muscle tissue across multiple conditions, including normal aging (WT 3 months vs 1 year), disease-associated aging (BM 3 months vs 1 year), early disease (WT vs BM at 3 months), and late disease (WT vs BM at 1 year) (Figure 3A).

**Figure 3:**
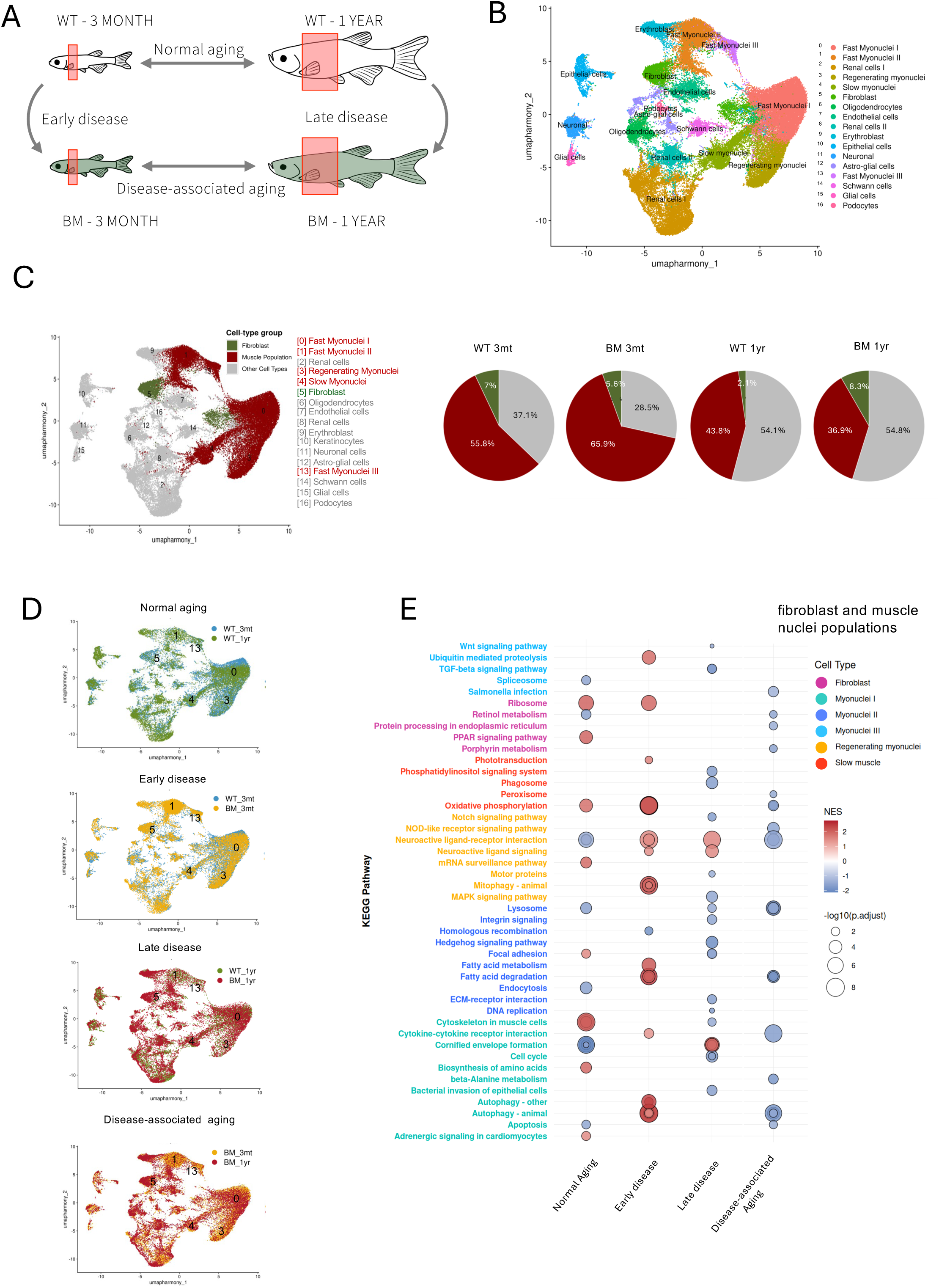
Single-nucleus RNA sequencing reveals condition-specific transcriptional states in skeletal muscle. **A.** Schematic representation of trunk skeletal muscle sampling across experimental conditions. The skin was removed mechanically prior to trunk dissection. Samples were collected to enable comparative analysis of transcriptional profiles associated with normal aging, early disease, late disease, and disease-associated aging. **B.** UMAP plot of integrated snRNA-seq data from WT and BM zebrafish trunk skeletal muscle at 3 months and 1 year. Distinct clusters (0 to 16) are color-coded and annotated according to identified cell types or nuclei populations. **C.** Left panel: Condition specific UMAP plot highlighting muscle nuclei populations and fibroblast clusters in dark red and green, respectively. Right panel: Pie charts showing shifts in the muscle (dark red) and fibroblast (green) nuclei populations compared to total nuclei analyzed (in %) in each sample, WT and BM at 3 months and 1 year. **D.** Split UMAP plots comparing the four pairwise conditions: normal aging (WT 3mt vs. WT 1yr), early disease (WT 3mt vs. BM 3mt), late disease (WT 1yr vs. BM 1yr) and disease-associated aging (BM 3mt vs. BM 1yr). **E.** KEGG pathway enrichment analysis (GSEA) across the four conditions comparison in muscle nuclei populations and fibroblasts across conditions including normal aging, early disease, late disease and disease-associated aging. Bubble size represents −log₁₀ (adjusted p-value); bubble colour represents normalised enrichment score (NES; red = positive enrichment, blue = negative enrichment).

Unsupervised clustering of nuclei based on known markers identified 17 distinct clusters (Figure 3B and Figure S3, Table S1), including five muscle-related clusters corresponding to fast (clusters 0, 1, 3, and 13) and slow (cluster 4) myofibers, as well as one fibroblast cluster (cluster 5). Additional clusters corresponded to neural and glial populations, including oligodendrocytes, astro-glial cells, glial cells, Schwann cells, and one unidentified neuronal-like population (cluster 11).

Quantification of cell-type or myonuclear populations composition revealed a progressive reduction in the proportion of muscle nuclei with age in both genotypes, with the most pronounced decrease observed in BM fish (from 65.9% at 3 months to 36.9% at 1 year) (Figure 3C). In contrast, fibroblast abundance increased markedly in BM fish at the later stage compared to WT, consistent with progressive fibrotic remodeling. Together, these results indicate an imbalance between muscle and fibroblast populations during disease progression. Split UMAP of condition-specific analysis further revealed that regenerating myonuclei decrease with normal aging, whereas they transiently increase in early disease, potentially reflecting a compensatory response to muscle damage. This regenerative capacity declines at later stages and during disease aging, suggesting exhaustion of the regenerative pool over time (Figure 3D). In WT fish, fibroblast numbers decreased with age, consistent with physiological aging. This pattern supports an early and persistent fibrotic response in diseased muscle. Importantly, the increase in fibroblasts, together with reduced muscle nuclei, indicates a shift from regenerative myogenesis toward a fibrogenic program and impaired muscle repair capacity.

KEGG pathway analysis further supported this interpretation. In muscle nuclei, early disease was associated with enrichment of stress-response and quality control pathways, including mitophagy and autophagy in regenerating and fast myonuclei, as well as ubiquitin-mediated proteolysis in additional myonuclear populations (Figure 3E). These pathways reflect activation of proteostatic and organelle quality control mechanisms, likely aimed at mitigating intrinsic cellular damage. In addition, metabolic remodeling was evident, with enrichment of fatty acid metabolism and oxidative phosphorylation pathways, suggesting an adaptive response to sustain energy demands during stress. Notably, enrichment of neuroactive ligand–receptor interaction pathways in regenerating myonuclei suggests altered neuromuscular signaling, indicating that regenerative processes may also be functionally altered. Fibroblasts, in contrast, exhibited enrichment of ribosomal pathways, indicating increased translational activity and biosynthetic capacity. This suggests that fibroblasts are not merely more abundant in BM muscle but are also transcriptionally and functionally activated during early disease stages.

Overall, snRNA-seq analysis reveals that collagen VI deficiency induces an early and coordinated muscle stress response characterized by activation of proteostatic and metabolic pathways, alongside altered regenerative signaling. In parallel, fibroblasts become transcriptionally activated and biosynthetically primed, supporting an early shift from regenerative myogenesis toward fibrotic remodeling. These findings identify fibroblasts as key early responders in the pathological progression of collagen VI-related myopathy.

### Fibroblasts exhibit increased stress responses and undergo functional decline during disease progression

As the disease progresses, fibroblasts exhibit marked changes in pathways related to proteostasis. In particular, KEGG analysis revealed a negative enrichment of endoplasmic reticulum (ER) protein processing pathways in aged BM fibroblasts (Figure 3E), suggesting a defective ER-associated protein folding and quality control capacity. Given that aging is generally associated with impaired proteostasis and reduced efficiency of ER quality control systems, leading to accumulation of damaged or misfolded proteins (Lopez-Otin et al, 2023), we investigated whether a similar process occurs in fibroblasts within the disease context. Consistent with this hypothesis, fibroblasts from older myopathic fish displayed increased expression of ER stress–associated genes, including *capn1*, *calr3a*, and *edem1* (Figure 4A). These genes are functionally linked to proteostatic stress responses: *capn1* encodes a stress-associated protease, *edem1* plays a key role in ER-associated degradation (ERAD) by targeting misfolded glycoproteins, and *calr3a* encodes a chaperone-related protein involved in protein folding. Together, these findings indicate activation of compensatory stress-response pathways in the context of impaired ER function. Collagen VI genes (*col6a1–3*) are predominantly expressed by fibroblasts, as expected (Figure 4B). We hypothesize that the accumulation of misfolded mutant collagen VI contributes to ER stress in fibroblasts. Supporting this idea, we previously showed that mutant collagen VI accumulates within fibroblasts in isolated skeletal muscle bundles (Idoux et al., 2025). To further investigate collagen VI biosynthesis, we generated transient CRISPR/Cas9-mediated knock-in embryos in which endogenous *col6a1* was fused to a fluorescent reporter mNeonGreen (mNG). In WT crispants, the ColVI–mNG fusion protein was primarily detected as thin extracellular filaments, with minimal intracellular accumulation (Figure 4C). In contrast, BM crispants exhibited prominent intracellular and extracellular aggregates of ColVI–mNG, indicating altered biosynthesis and/or secretion from early developmental stages (Figure 4C). In support to this, our snRNAseq data indicate a progressive increase in *col6a1–3* expression in BM fibroblasts from 3 months to 1 year, pointing to sustained or elevated collagen VI production over time (Figure 4D). We therefore tested the effect of a 4-day treatment with selected compounds, 4-PBA, carbamazepine (CBZ) and taurursodiol (TUDCA), known for their effect on ER-stress and unfolded protein response, on ColVI secretion in 5 dpf WT and BM larvae. The effect of compounds on ColVI secretion was then assessed at the myoseptal level with whole-mount immunofluorescence with anti-ColVI antibodies (Figure 4E). The presence of collagen VI short thin filaments versus patches in one vertical myoseptum (per hemisegment) was scored at 1, the persistence of patches along the myoseptum was scored as 0 (6 myosepta per larvae were analyzed, the maximum score per larva being 6). Unlike BM larvae treated with 4-PBA, improvement on ColVI secretion and assembly, as judged by ColVI immunostaining of extracellular deposition, was significantly observed following TUDCA treatment (Figure 4E). Taken together, these results support a model in which fibroblasts initially increase collagen production but progressively develop an imbalance between protein synthesis and processing capacity. This leads to chronic ER stress and impaired proteostasis, ultimately resulting in fibroblast dysfunction. Such a transition from activation to stress and dysfunction is consistent with mechanisms described in fibrosis and aging, and may contribute to the altered extracellular matrix and tissue remodeling observed in BM muscle.

**Figure 4:**
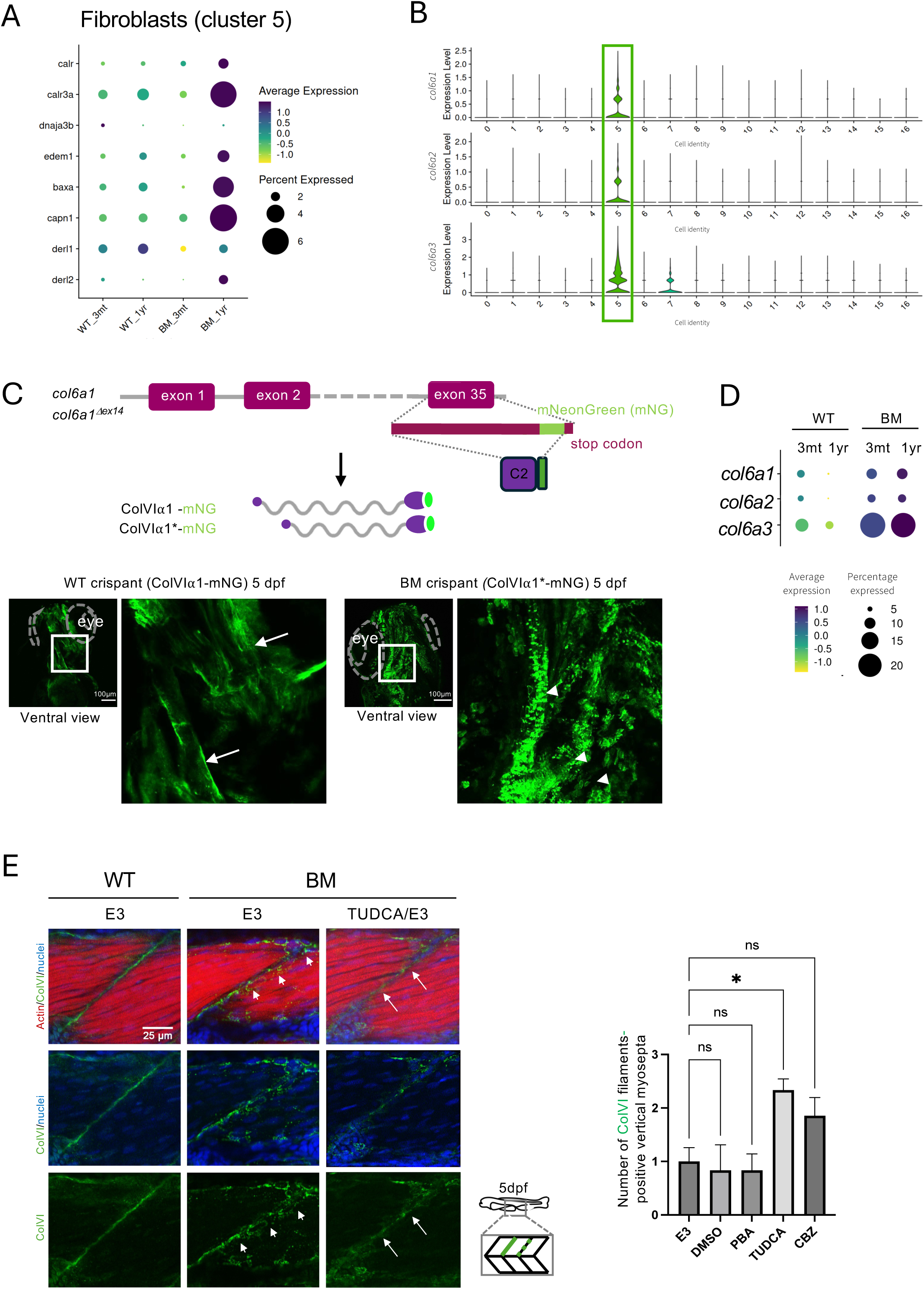
BM fibroblasts develop an ER stress gene signature at late disease stage. **A.** Expression of canonical ER stress markers in fibroblasts (cluster 5) across conditions (left panel) compared to all clusters (0-16, right panel). **B.** Violin plots showing expression levels of *col6a1*, *col6a2*, and *col6a3* in all clusters, revealing that fibroblasts (cluster 5) are the primary source of collagen VI transcripts (highlighted in green). **C.** Top panel: Schematic of the Crispr/Cas9 mediated knock in strategy used to express mosaically a ColVI⍺1–mNeonGreen (mNG) fusion protein both in WT (ColVI⍺1-mNG) and BM (ColVI⍺1*-mNG) larvae. Lower panel: Confocal imaging of mNeonGreen (mNG) signal in WT and BM crispants at 5 dpf, focusing on skeletal muscle in the head region. In WT larvae, arrows indicate extracellular assembly of the fluorescent collagen VI fusion protein. In BM crispants, arrowheads point to bright patches with no distinction between intracellular retention or extracellular deposition. Scale bars = 100 µm. **D.** Dot plot of the expression levels of collagen VI genes (*col6a1*, *col6a2*, and *col6a3*) in BM at 3 months and 1-year old fish compared to WT (snRNAseq data) in cluster 5 (fibroblasts). **E.** Left, Immunostaining for ColVI of 5dpf WT, BM and BM treated with TUDCA. Arrows point to ColVI filaments compared to patches (arrowheads). Scale bars = 25µm. Right, Quantification of ColVI deposition as filaments. Statistics were performed using ordinary ONE-way ANOVA test. Data are mean ± SEM (N=6).

### Fibroblasts acquire pro-fibrotic and microenvironment-regulating signaling signatures during disease progression

Fibrotic remodeling has previously been reported in this model (Radev et al., 2015). In the present study, differential gene expression analysis revealed upregulation of *calpain-1* in BM fish, a protease known to play a key role in extracellular matrix (ECM) remodeling through activation of MMP2 in fibrotic contexts (Jiang et al., 2012). This observation prompted us to further investigate ECM dynamics and remodeling in BM muscle during disease progression. To this end, we characterized the matrisome of BM and WT fibroblasts (cluster 5) at 3 months and 1 year using the zebrafish matrisome database ((Nauroy et al., 2018); http://matrisomeproject.mit.edu) (Tables S2–S5). Analysis across all matrisome categories, including core matrisome components (collagens, proteoglycans, glycoproteins), ECM regulators, ECM-affiliated proteins, and secreted factors, revealed a marked increase in gene expression in BM fish at 1 year compared to WT (Figure 5A). This global upregulation was further supported by the number of differentially expressed genes (DEGs) in the disease-associated aging condition, particularly within ECM glycoproteins and collagens, indicating a strong disease-associated remodeling signature (Figure 5B). These transcriptional changes were predominantly observed in fibroblasts (cluster 5) across all matrisome categories (Figure S4). To further explore the organization of these changes, we performed protein–protein interaction analysis (STRING data base) (Tables S6-S7), which revealed a highly interconnected and complex ECM network specifically in diseased aging conditions, in contrast to normal aging. Within this network, three major functional hubs emerged: ECM remodeling, collagen biosynthesis and modifying enzymes, and collagen crosslinking. The ECM remodeling hub comprised genes upregulated in 1-year BM fibroblasts, including *mmp2*, *mmp14a/b*, *adam10*, *sparc*, and *fstl1a*, all of which are associated with fibrotic processes (Figure 5C; Tables S6–S7) (Li et al., 2021; Trombetta-eSilva & Bradshaw, 2012). Notably, members of the ADAMTS metalloproteinase family (Yuan et al., 2026) were also enriched, further supporting extensive ECM remodeling during disease progression. In line with previous reports, the interaction between calpain-1 and MMP2 may enhance ECM remodeling through increased activation of MMP2 and subsequent collagen deposition (Jiang et al., 2012). The second hub, related to collagen biosynthesis and modification, included multiple collagen genes (*col1a2, col5a1–3, col6a1–3, col8a1*) along with key modifying enzymes such as *plod2*, *p4ha1a*, and *tll1*. The third hub comprised genes involved in collagen fibril crosslinking, including members of the *lox* family, which are critical for ECM stabilization and stiffness (Figure 5C). Consistent with these transcriptomic findings, immunofluorescence staining of 1-year BM muscle sections using collagen XII (ColXII) confirmed increased ECM deposition and an elevated number of nuclei within the myomatrix (Figure 5D), indicative of fibrotic tissue formation. Finally, fibroblasts from BM fish exhibited increased expression of genes encoding axon guidance molecules, including semaphorins, plexins, netrins, and slit family members Figure C, Tables S7). This suggests that, beyond ECM production, fibroblasts acquire a signaling profile capable of influencing nerve guidance and potentially vascular remodeling, reflecting a broader microenvironmental reprogramming during late-stage disease.

**Figure 5:**
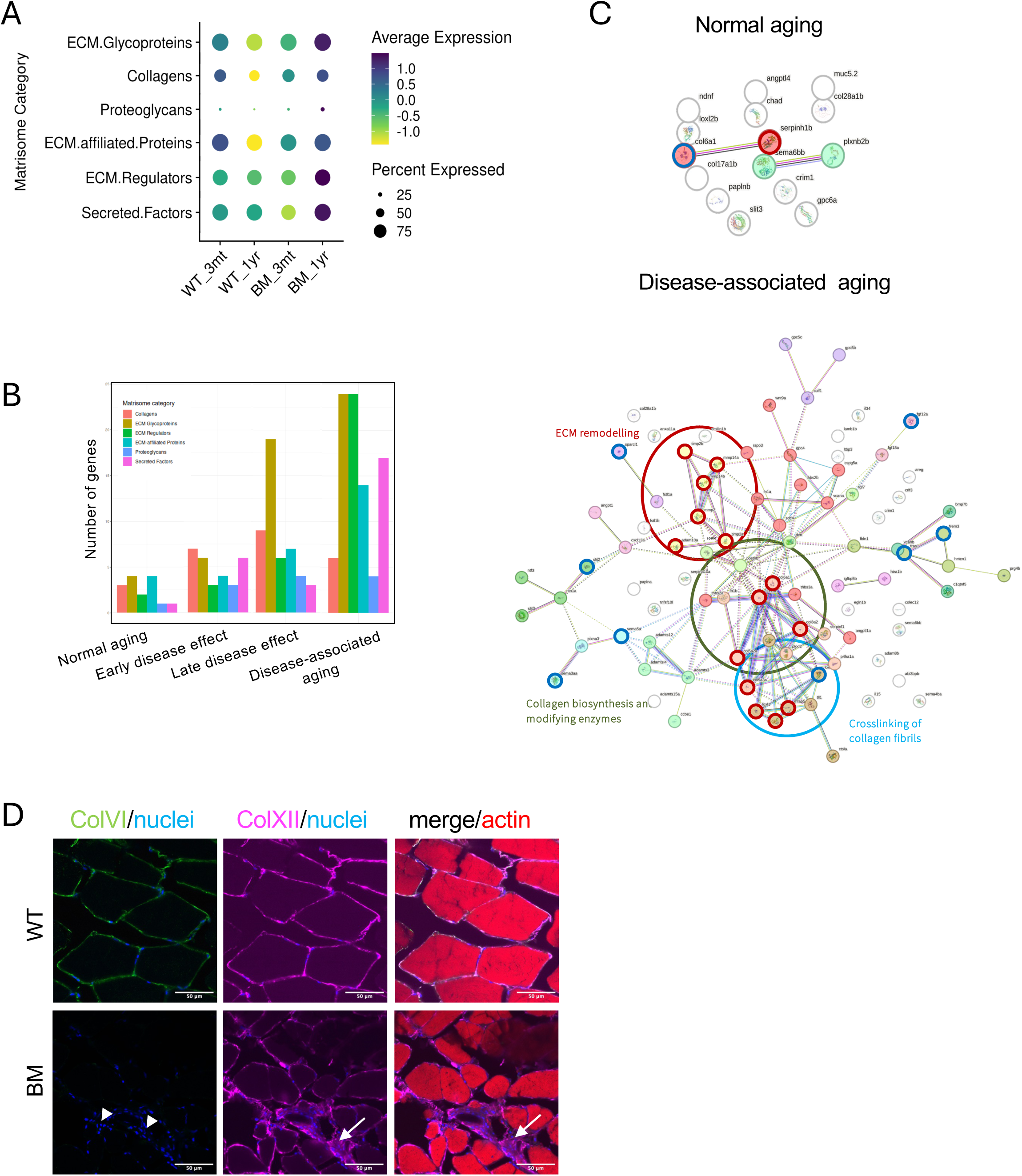
BM fibroblasts drive fibrotic ECM remodeling and altered muscle cross-talk at late stages. **A.** Dot plot showing expression levels of matrisome category genes in BM and WT fibroblasts at 3 months and 1-year. **B.** Number of differentially expressed (DEGs) per matrisome category genes in fibroblasts across all conditions (normal aging, early disease, late disease and disease-associated aging). **C.** Protein-protein interaction network analysis of the matrisome DEGs expressed by fibroblasts in normal ageing (left panel, 35 genes) and disease-associated aging (right panel, 194 genes) with the STRING database. The protein interaction networks were created with matrisome DEGs (p_val_adj < 0.05 and avg_log2FC > 0.25 and ran STRING for MCL clustering of threshold 0.3). Line edge thickness reflects interaction confidence score. The top few up-regulated genes (avg_log2FC > 0 at 1year condition) circled in red and down-regulated (avg_log2FC < 0 at 1year condition) circled in blue. Three hubs of interest are highlighted: ECM remodeling (red), collagen biosynthesis and modifying enzymes (green) and cross-linking of collagen fibrils genes (blue). **D**. Immunofluorescent staining of 1-year old WT and BM cryosections with anti-ColXII (magenta) and anti-ColVI (green). Nuclei are in blue, merge images show staining with anti-ColXII (magenta), anti-ColVI (green), and phalloidin-alexa647 (actin, red). Arrowheads indicate clusters of nuclei in the fibrotic area; arrows point to ColXII patches in BM fish. Scale bars = 50 µm.

### Fibroblast–muscle crosstalk is progressively dysregulated in BM fish during aging

Given the central role of fibroblasts in shaping the extracellular matrix and mediating intercellular signaling, we next investigated how fibroblast–muscle communication evolves during disease progression. Using CellChat, we first assessed global ligand–receptor interactions across all cell types. In wild-type (WT) fish, the total number of inferred interactions increased modestly with age (from 330 to 422). In contrast, BM fish exhibited a marked increase, from 260 interactions at 3 months to 1,261 at 1 year, indicating a substantial amplification of intercellular communication during disease progression (Figure 6A and B). A similar trend was observed for overall interaction strength, which was reduced in BM compared to WT at 3 months but markedly increased at 1 year (Figure 6A).

**Figure 6:**
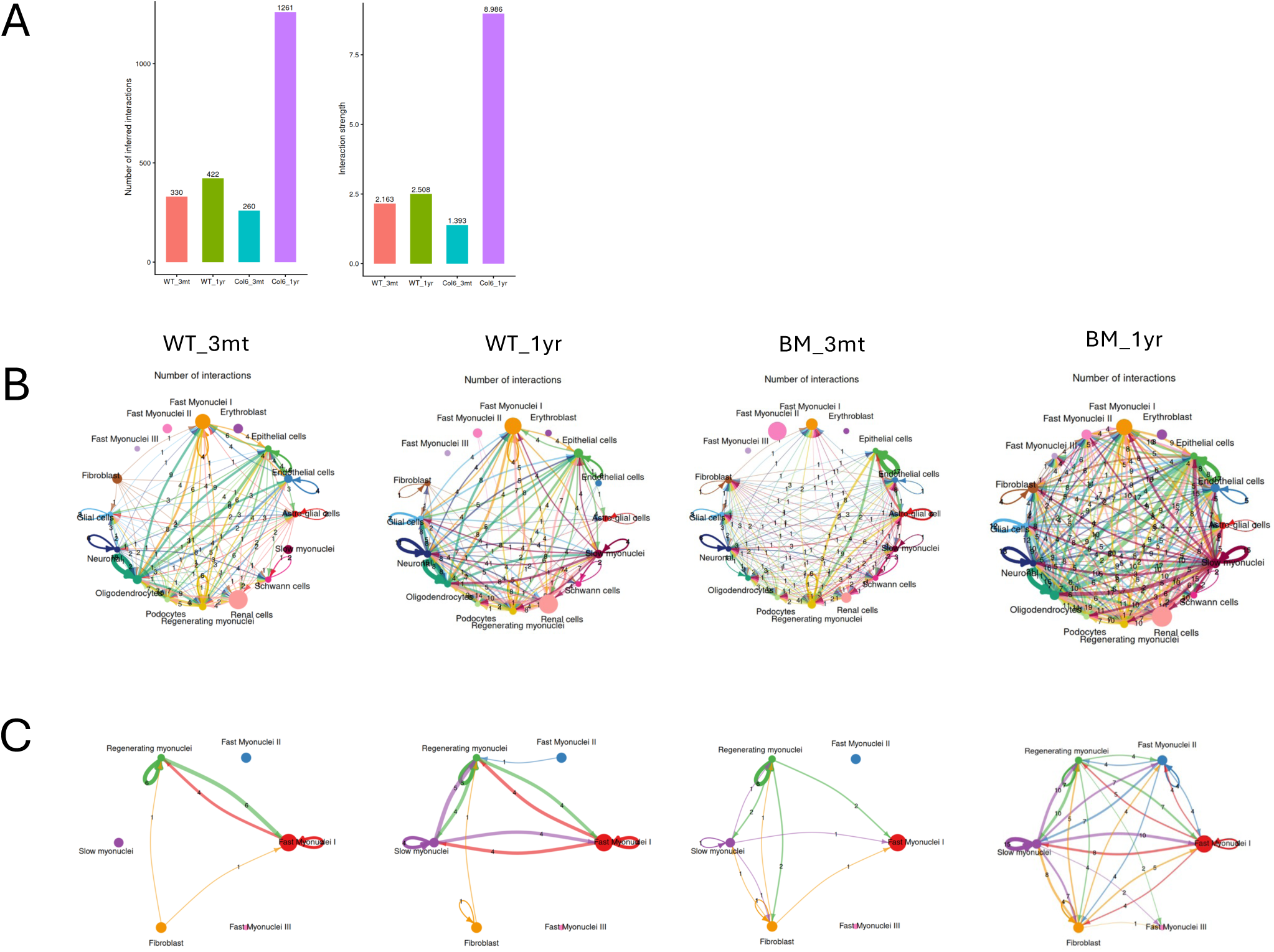
Fibroblast-myofibre crosstalk is progressively amplified in BM fish. **A.** CellChat analysis of inferred interactions, including the number of interactions (left histogram) and interaction strength (right histogram). **B.** CellChat circle plots showing the number of inferred interactions between all cell populations in each condition. Node size reflects the number of cells in each population; edge thickness reflects the number of interactions; edge color indicates the sending cell type, in WT and BM samples at 3 months and 1 year. **C.** Predicted ligand–receptor interactions between myonuclear populations (clusters 0, 1, 3, 4, and 13) and fibroblasts (cluster 5) in WT and BM conditions at 3 months and 1 year.

To specifically examine fibroblast–muscle communication, we focused on interactions between fibroblasts and myonuclear populations (Figure 6C). In WT fish, predicted crosstalk between fibroblasts and myofibers remained limited and relatively stable with age. Age-related changes in WT were primarily restricted to interactions involving regenerating, fast, and slow myonuclei, consistent with a role for fibroblast signaling in supporting muscle maintenance and repair during physiological aging. In contrast, BM fish displayed a pronounced and progressive increase in predicted interactions between fibroblasts and all myonuclear populations (Figure 6C). This increase was already evident at 3 months compared to age-matched WT and became substantially amplified at 1 year. These findings suggest persistent and widespread signaling between fibroblasts and muscle cells in response to ongoing muscle damage. This is consistent with a persistent attempt to maintain muscle integrity in the context of collagen VI deficiency within the myomatrix (Figure 1A and B), despite continued expression of *col6a1–3* genes by fibroblasts (Figure 4D). Notably, the marked increase in fibroblast–myofiber communication may also reflect qualitative alterations in signaling.

Although fibroblasts appear to enhance extracellular matrix production, the remodeled myomatrix may be functionally altered and unable to properly mediate signaling to muscle fibers. Together, these results indicate that fibroblast–muscle communication is not simply reduced or lost in BM fish but becomes progressively amplified and dysregulated with age.

## DISCUSSION

Bethlem myopathy (BM) is a progressive muscle disorder in which the mechanisms driving age-dependent disease worsening remain incompletely understood. Clinically, BM patients display features reminiscent of premature aging. Zebrafish offer genetic tractability, rapid aging, suitability for pharmacological studies, and the possibility to investigate disease progression *in vivo* (Patton et al., 2021). Here, using a zebrafish model carrying a pathogenic collagen VI mutation (Idoux et al., 2025; Radev et al., 2015), we show that this model not only reproduces hallmark features of the human disease (Radev et al., 2015), but also recapitulates its progressive nature (Deconinck et al., 2015).

Although COL6-related myopathies (COL6-RM) have primarily been studied as skeletal muscle disorders, skeletal abnormalities are increasingly recognized as part of the disease spectrum (Bönnemann, 2011; Di Martino et al., 2023b). Mouse models lacking *Col6a1 or Col6a2* exhibit reduced trabecular bone mass, altered osteoblast connectivity, and increased osteoclast activity (Christensen et al., 2012; Pham et al., 2020), although the mechanistic basis of these alterations and their relationship to muscle pathology remain unresolved. Collagen VI is primarily expressed by fibroblasts in skeletal muscle, but is also present in bone, where it contributes to maintain bone mass (Izu et al., 2012). In zebrafish, ColVI is expressed in craniofacial elements during embryonic development (Tonelotto et al., 2019). In the present study, BM fish exhibited progressive muscle wasting, increased myofiber size variability, and secondary skeletal deformities, including scoliosis and jaw abnormalities, consistent with clinical observations in patients (Baker et al., 2007; Bönnemann, 2011; Briñas et al., 2010). Importantly, our morphological and gene expression analyses suggest that progressive muscle weakness is more likely the primary driver of the late-onset skeletal phenotype than intrinsic bone defects. Supporting this interpretation, skeletal development appeared normal in BM larvae and juveniles based on histological analyses. In addition, expression levels of key bone remodeling markers involved in osteogenesis and remodeling (Valenti et al., 2020), including runx2 (early osteogenic marker), sparc (early-to-intermediate osteogenic marker), and bglap (late osteogenic marker), were unchanged between 1-year old BM and WT fish. This interpretation is consistent with the concept of muscle-bone coupling described in disorders such as osteogenesis imperfecta, where skeletal abnormalities partly result from impaired mechanotransduction between muscle and bone (Gremminger & Phillips, 2021). It also aligns with clinical observations in BM patients, in whom scoliosis and skeletal deformities generally emerge progressively beyond childhood (Bönnemann, 2011). Together, these observations suggest that the progressive musculoskeletal phenotype is accompanied by early molecular adaptations within skeletal muscle itself.

At the molecular level, single-nuclei RNA sequencing revealed that collagen VI deficiency triggers an early and coordinated stress response in skeletal muscle. Myonuclei displayed enrichment of pathways related to autophagy, mitophagy, and ubiquitin-mediated proteolysis, consistent with activation of proteostatic and organelle quality-control mechanisms. Such adaptive responses are well documented in stressed muscle and muscular dystrophies (Grumati et al., 2010; Romanello & Sandri, 2016). Concomitant metabolic rewiring, including enrichment of “oxidative phosphorylation” and “fatty acid metabolism” pathways (KEEG analysis), further suggests an attempt to sustain energy demands under chronic stress conditions (Argilés et al., 2016). Notably, enrichment of neuroactive ligand–receptor interaction pathways in regenerating myonuclei points to altered neuromuscular communication, a feature previously implicated in COL6-RM progression (Cescon et al., 2018).

Disease muscle fibroblasts appear to acquire signaling functions that may actively shape the tissue microenvironment, suggesting that, on their side, they follow pathways that are distinct from those associated with diseased muscle. Upregulation of axon guidance molecules, including semaphorins, plexins, netrins, and Slit proteins, suggests potential roles in neuromuscular remodeling and vascular patterning. These molecules are increasingly recognized to function beyond the nervous system, including in angiogenesis and tissue repair (Adams & Eichmann, 2010; Tamagnone, 2012). Beyond extracellular matrix (ECM) production, d

A central finding of this study is the early and sustained involvement of fibroblasts in disease progression. One-year-old BM skeletal muscle displayed a marked imbalance between myonuclei and fibroblasts, characterized by increased fibroblast abundance and reduced myonuclear representation. This shift suggests a transition from regenerative myogenesis toward fibrogenesis and is consistent with the observed loss of body weight and pronounced variability in myofiber size, already detectable at early stages and becoming more severe with age. These findings are in agreement with studies in other muscular dystrophies, where fibro-adipogenic progenitors and fibroblasts contribute to fibrosis and impaired regeneration (Pessina et al., 2015; Uezumi et al., 2011).

The early enrichment of ribosomal pathways in fibroblasts suggests that these cells are transcriptionally and biosynthetically activated. As disease progresses, however, fibroblasts appear to transition from activation to dysfunction. We observed reduced enrichment of ER protein-processing pathways in 1-year BM fish together with increased expression of ER stress markers such as *capn1*, *calr3a*, and *edem1*. This profile is consistent with impaired proteostasis and activation of the unfolded protein response (UPR), hallmarks of chronic cellular stress and aging (Hetz & Saxena, 2017; Kaushik & Cuervo, 2015). Given that collagen VI is predominantly produced by fibroblasts, intracellular accumulation of misfolded mutant protein likely represents a major source of proteotoxic stress. Similar mechanisms have been described in other connective tissue disorders involving collagen misfolding and ER stress (Ishikawa & Bächinger, 2013). Our *in vivo* imaging data further support this model, demonstrating intracellular retention and aberrant deposition of mutant collagen VI from early developmental stages through adulthood (this study; Idoux et al., 2025). In parallel, the progressive increase in *col6a1* expression despite defective protein processing suggests a maladaptive compensatory feedback loop in which fibroblasts attempt to restore ECM integrity but ultimately exacerbate cellular stress.

Chemical chaperones such as 4-PBA and TUDCA, both FDA-approved compounds, can modulate protein folding and secretion by alleviating ER quality-control constraints and promoting export of misfolded proteins (Balch et al., 2008; Sitia & Braakman, 2003). Such approaches have been explored in several connective tissue disorders associated with collagen mutations. For example, 4-PBA, but not CBZ, reduced ER stress in fibroblasts derived from patients with vascular Ehlers–Danlos syndrome caused by *COL3A1* mutations (Omar et al., 2025). In a zebrafish model of osteogenesis imperfecta (*Chihuahua* mutant), characterized by intracellular accumulation of misfolded collagen I, prolonged treatment with 4-PBA, but not TUDCA, reduced disease-associated defects (Gioia et al., 2017). In a mouse model of chondrodysplasia caused by *Col10a1* mutations, stimulation of intracellular proteolysis with CBZ ameliorated abnormalities associated with hypertrophic chondrocyte differentiation (Mullan et al., 2017). In our zebrafish BM model, intracellular accumulation of collagen VI was observed both at early stages (this study) and later stages (Idoux et al., 2025), while extracellular collagen VI progressively disappeared from skeletal muscle. Under these conditions, TUDCA, but not CBZ or 4-PBA, partially restored collagen VI secretion. The relatively short treatment duration (4 days) in larvae may have limited our ability to detect additional therapeutic effects, and longer treatments or combinatorial approaches may reveal greater efficacy. Although preliminary, these findings suggest that modulation of proteostasis pathways may represent a promising therapeutic avenue for BM.

At later stages, fibroblasts acquired a pronounced pro-fibrotic phenotype characterized by activation of ECM remodeling pathways and formation of a complex matrisome network. This is in agreement with previous histological analysis showing fibrotic tissue in the skeletal muscle of 5-monthold BM fish (Radev et al., 2015). Functional hubs involving metalloproteinases (MMPs and ADAMTS family members), collagen biosynthesis enzymes, and crosslinking factors of the LOX family were identified, consistent with established mechanisms of fibrosis (Theocharis et al., 2016; Wynn & Ramalingam, 2012). The interplay between calpain-1 and MMP2, previously implicated in ECM remodeling and fibrosis (Jiang et al., 2012), may represent an important driver of this process in BM muscle.

Importantly, immunofluorescence analyses confirmed increased collagen XII deposition and fibrotic tissue accumulation, validating the transcriptomic findings at the tissue level. Collagen XII is expressed in zebrafish fascia during development (Bader et al., 2009), and fascia-associated fibroblasts constitute a major cellular source during tissue repair (Knoedler et al., 2023). Moreover, mutations in *COL12A1* cause myopathic Ehlers–Danlos syndrome (mEDS), an early-onset disorder characterized by connective tissue abnormalities and muscle weakness (Izu & Birk, 2023). Increased collagen XII expression has also been reported during tissue regeneration in both the heart (Marro et al., 2016) and regenerating zebrafish spinal tissue, where it was proposed to promote regenerative processes (Wehner et al., 2017).

Interestingly, CellChat analysis revealed a striking amplification of fibroblast–muscle communication during disease progression. Whereas physiological aging was associated with relatively modest and selective intercellular interactions, BM muscle exhibited widespread and intensified signaling between fibroblasts and all myonuclear populations. This likely reflects a chronic attempt to preserve tissue integrity in the face of persistent damage. However, the qualitative nature of these interactions may also be altered, as ECM remodeling is known to profoundly influence mechanical signal transduction and cellular behavior (Humphrey et al., 2014). Thus, enhanced intercellular communication may not be protective, but instead contribute to disease progression by reinforcing maladaptive signaling loops within a structurally altered ECM environment.

Overall, our findings support a model in which collagen VI deficiency initiates an early stress response in muscle fibers, rapidly followed by fibroblast activation. Over time, the regenerative myonuclear population declines, while fibroblasts become proteostatically stressed and functionally impaired, simultaneously driving extracellular matrix remodeling and fibrosis. This dual role positions fibroblasts as central regulators of disease progression in Bethlem myopathy and highlights them as particularly attractive therapeutic targets, especially given their responsiveness to chemical chaperone treatment.

## MATERIALS AND METHODS

### Zebrafish Maintenance and Ethical Statement

Zebrafish (AB/TU) maintenance and embryo collection were performed at the zebrafish PRECI facility (Plateau de Recherche Expérimentale de Criblage In vivo, UMS CNRS 3444 Lyon Biosciences, Gerland) in compliance with French government guidelines for animal welfare (agreement number C693870602). Embryos obtained from natural spawning were raised following standard conditions. Zebrafish (Danio rerio) were maintained in a recirculating system at 28°C under a 12-h light/12-h dark cycle. Adult fish were fed thrice daily with a combination of dry food and fresh Artemia salina. Developmental stages are given in hour post-fertilization (hpf) at 28.5°C according to morphological criteria (Kimmel et al, 1995). Tricaine (ms-222, Sigma-Aldrich, St Louis, Missouri, USA) was used to anesthetize the fish before the larvae were embedded in LM agarose. For experimentation, 24 hpf embryos were treated with phenylthiourea (P7629, Sigma-Aldrich, St Louis, Missouri, USA) to prevent pigmentation. All animal manipulations were performed in agreement with EU Directive 2010/63/EU and approved by the French ethical committee (APAFIS#28791-2020102114261517 v4 and APAFIS #52184-2024062116505409 v8). The creation of the *col6a1^τιex14-/-^* line (BM fish), fin clipping, DNA extraction and genotyping were previously described in (Radev et al, 2015).

### Morphometric Analysis

For all imaging and morphological assessments, fish were anesthetized using Tricaine methane sulfonate (MS-222). Anesthesia was confirmed by the absence of opercular movement and response to physical stimulation. Body weight was measured to the nearest 0.01 g using an analytical balance. Standard and total lengths were recorded using a standardized ruler, with the caudal fin positioned in a relaxed, resting state.

Craniofacial imaging and geometric morphometrics were performed using High-resolution imaging was performed using a Keyence VHX-7000 digital microscope. Fish were imaged using a VHX-E20 low-magnification lens (20x–100x). Lighting was standardized using epi-illumination with a full ring setting at 80% brightness. Images were captured using the serial recording method with 3D image stitching. The Z-stage was calibrated to the top-most focal plane (apex of the body) and the bottom-most focal plane (caudal fin), with a vertical pitch of 100 µm. A compressed 2D image was generated, calibrated with a scale bar, and saved as a TIFF file. Both lateral sides of the fish were imaged to evaluate asymmetry.

Geometric morphometrics were performed using TPSUtil to format images into TPS files. Landmarks were assigned on 2D images using TPSDig2. Eight anatomical landmarks were selected, modified from Miyashita et al. (2020), to capture the craniofacial configuration: (1) anterior orbit, (2) posterior orbit, (3) ventral hypobranchial region, (4) posterior lower lip, (5) anterior lower lip, (6) lip junction, (7) anterior upper lip, and (8) trunk–head boundary. Data were analyzed in MorphoJ (Klingenberg, 2011) via Procrustes superimposition, followed by Principal Component Analysis (PCA) and Canonical Variate Analysis (CVA).

### Μicro-Computed Tomography (µCT)

Skeletal structure was assessed using a SKYSCAN 1272 CMOS Edition (Bruker). Samples were fixed in 4% PFA with 0.9 mM CaCl_2_ and 0.49 mM MgCl_2_ for 24 h at 4°C, then dehydrated through a graded ethanol series to a final concentration of 70% ethanol. Samples were scanned immersed in 70% ethanol to prevent dehydration.

Scanning parameters included: voxel size 5 µm; voltage 60 kV; 0.71 mm Al filter; 1800 ms exposure; 0.71° rotation step; and 3-frame averaging over 180°. Data were reconstructed using NRecon software (Bruker) with the following settings: smoothing 0, ring artifact reduction 3, and beam-hardening correction 40%. Skeletons were visualized using DataViewer (Bruker), segmented in CTAn, and volume measurements were computed using Avizo (Thermo Fisher Scientific).

### Whole-Mount Staining (Alizarin Red/Alcian Blue)

This protocol was adapted from Gioia, R., et al. (2017). 4 dpf larvae were stained with Alcian blue and 1-month juveniles were double stained (Alcian Blue/Alizarin Red). For double staining, fish were anesthetized, scales and internal organs were manually removed, and specimens were fixed in 4% PFA in PBS (pH 7.4) supplemented with CaCl2 and MgCl2 at 4°C. Samples were washed in tap water for 30 min. Cartilage staining was performed by overnight incubation in 0.04% Alcian Blue 8GX (Sigma-Aldrich) in 80% EtOH containing 10 mM MgCl2. Samples were rehydrated in an ethanol series (80%, 50%, 25%) and water (1 h each). Pigmentation was bleached using 1% $H2O2 /1% KOH. Following a 30-min water wash, specimens were incubated in 30% saturated sodium tetraborate for 12 h. Bone staining was performed using 1 mg/mL Alizarin Red S in 1% KOH overnight. All specimens were cleared in a glycerol/0.1% KOH series (50% and 70%) and stored in 80% glycerol.

### Gene Expression Analysis (RT-qPCR)

Heads and trunks were dissected and separated. Cranial tissue was isolated after careful removal of the brain. Tissues were stored in RNAlater until processing. Total RNA was extracted using the RNeasy Mini Kit (Qiagen) following homogenization via QIAshredder (Qiagen). cDNA synthesis was synthesized from 1 μg of RNA using the SuperScript First-Strand Synthesis System (Thermo Fisher Scientific), according to the manufacturer’s instructions. Quantitative PCR was performed with PowerUp SYBR Green Master Mix (A25777, Thermo Fisher Scientific). Gene expression was normalized to *actb* expression. Primers used include: *runx2* F: 5’-CTTCAATGACCTGCGCTTTGT-3’, R: 5’-TCGGAGAGTCATCCAGCTTC*-3’; sparc* F: 5’-CCCTCTGCGTGCTCCTCTTA-3’, R: 5’-GCATCGCACTGCTCAAAGAA-3’; *bglap F: 5’-*CTGCTGCCTGATGACTGTGT-3’, R: 5’-CACGCTTCACAAACACACCT-3’; *actb* F: 5’-GCCAACAGGGAAAAGATGAC-3’, R: 5’-GACACCATCACCAGAGTCCA-3’.

### snRNA-seq analysis

Tissue Dissociation and Nuclei Isolation: The samples were euthanized using an overdose of tricaine and dissected in Tyrode solution. Two zebrafish per genotype (WT and BM mutant) at 3 months and 1 year of age were processed. Briefly, the head, caudal fin, dorsal fin, and anal fin were removed, followed by careful excision of the internal visceral organs. The skin was then gently peeled from both sides of the fish to expose the trunk musculature while minimizing tissue damage. The trunk muscle was divided into rostral and caudal portions by sectioning at the level of the anal opening. The rostral trunk muscle was snap-frozen on aluminum foil placed over liquid nitrogen and stored at −80 °C until further processing.

Frozen tissue (≤200 mg) was minced on dry ice and dissociated in a gentleMACS C-tube using ice-cold lysis buffer supplemented with 0.2 U/µL RNase inhibitor. Tissue was dissociated using the “4C_nuclei_1” program on a gentleMACS Dissociator (Miltenyi Biotec), filtered (70 µm), centrifuged at 300 x g (4°C, 5 min), and resuspended in nuclei-resuspension buffer. Nuclei were magnetically labeled with Anti-Nucleus MicroBeads and processed through LS columns. Final concentrations were adjusted to ∼8 × 10⁵ nuclei/mL. Encapsulation was performed using the 10X Chromium Controller. cDNA amplification (12 cycles) was followed by SPRIselect size selection. Libraries were sequenced as Dual-Index libraries (14 cycles). Bioinformatics of the data were performed by demultiplexing and alignment were performed using Cell Ranger v8.0.0 (reference: GRCz11). Matrix filtering (Seurat) included: genes ≥ 3 nuclei, nuclei ≥ 50 genes, and ≤ 2% mitochondrial reads. Data were SCT-normalized, integrated, and scaled prior to PCA and UMAP clustering. Raw reads were deposited to Array Express data base (access number pending).

### NGFP Line Generation (CRISPR/HDR)

A single guide RNA (sgRNA) targeting exon 35 of col6a1 (5′-AGATTTCCCTGGAGACGAGA-3′; PAM: GGG) was designed using CRISPOR v5.01 (Concordet and Haeussler, 2018) to induce a double-strand break two codons upstream of the stop codon. The sgRNA was synthesised using the EnGen sgRNA Synthesis Kit (New England Biolabs, NEB) from the oligonucleotide 5′-TTCTAATACGACTCACTATAGAGATTTCCCTGGAGACGAGAGTTTTAGAGCTAGA-3′, purified, and quality-verified by gel electrophoresis. The mNeonGreen donor template was prepared by a two-round nested PCR strategy.

Round 1 amplified the mNeonGreen open reading frame from a plasmid template (1 ng per 50 µL reaction) using Phusion high-fidelity polymerase with internal primers under the following cycling conditions: 98°C for 4 min; 5 cycles of 98°C/30 s, 64°C/20 s, 72°C/20 s; 35 cycles of 98°C/30 s, 72°C/20 s; final extension 72°C for 5 min. The Round 1 product was purified using a NucleoSpin PCR Clean-Up Kit (Macherey-Nagel) and quantified by NanoDrop.

Round 2 used 20 ng of the purified Round 1 product as template across eight parallel 50 µL reactions with biotinylated external primers under identical cycling conditions but with 30 extension cycles. All eight reactions were pooled, resolved on a 1% agarose gel, and the target band was excised and gel-purified, yielding a final donor concentration of approximately 300 ng/µL in nuclease-free water.

For microinjection, the sgRNA and Cas9 protein (NEB) were pre-assembled into a ribonucleoprotein (RNP) complex by incubation at room temperature for 5 minutes, then combined with the biotinylated donor template to produce an injection mix (500 pg gRNA -RNP complex, 50 pg donor template - round 2 PCR, 600pg Cas9 protein; Nuclease-free water (qsp to 15 µL). One-cell-stage embryos from wild-type (AB/TU) and col6a1Δex14 homozygous mutant backgrounds were injected at the one-cell stage and raised at 28°C in E3 medium. Successful knock-in founders were initially identified by screening injected larvae for mNeonGreen fluorescence. Integration was subsequently confirmed by junction PCR across the upstream and downstream junctions, and verified by Sanger sequencing.

### Immunofluorescence staining

Adult fishes were euthanized terminally anesthetized using 0.2% MS-222 (tricaine methanesulfonate, Sigma-Aldrich) and the trunk part from the mid gut to the end of dorsal fin is dissected. The trunk section is fixed overnight at 4 °C in 4% paraformaldehyde, followed by rinsing in 1× PBS. Samples are cryoprotected overnight in 30% sucrose. Blocks were then embedded in Tissue-Tek OCT, rapidly frozen in cold isopentane, and stored at −80 °C. Cryosections (10 µm) were obtained using a Leica CM3050 cryostat and mounted onto Superfrost Plus slides and stored at -20 till the time of staining. Sections were brought to room temperature, rehydrated in PBS, and blocked in 1% BSA and 3% sheep serum in PBS. Primary antibodies and phalloidin-647, when indicated, were applied in blocking solution and incubated either for 1 hour at room temperature or overnight at 4 °C. WGA staining was performed by incubating the samples with Wheat Germ Agglutinin (WGA; Alexa Fluor™ 488 Conjugate, Thermo Fisher Scientific, W11261) diluted 1:1000 in blocking solution for 1 hour at room temperature (RT), followed by three washes of 5 minutes each in PBS. After washing in PBS, sections were incubated with appropriate fluorescent secondary antibodies. Nuclei were stained using st 33342 (Invitrogen Hoechst 33342) for 10 mins in PBS. Following final washes, sections were mounted in 50% glycerol or DAKO mounting medium (Dako Faramount Aqueous Mounting Medium, Code S3025) in PBS and stored at 4 °C protected from light until imaging.

For whole-mount immunostaining, larvae were fixed with 4% PFA in PBS overnight at 4°C and processed to immunofluorescence staining without MeOH storage. Permeabilization step included a 10 min treatment to 5% triton in PBS for 10 min followed with proteinase K (Roche, 12 µg/ml) incubation at 37°C for 45 min. Larvae were then incubated in blocking buffer (1% BSA, 3% sheep serum) in PBDTT (1% DMSO, 0.1 % Tween-20, 0.5 % Triton X-100 in PBS) for 4 h at RT. Larvae were incubated in primary and secondary antibodies overnight at 4°. Heads and yolk sacs were removed and trunks were mounted in DAKO medium before imaging.

Primary and secondary antibodies were used at indicated dilutions: guinea pig polyclonal anti-ColVI⍺1 (1:1,500) (Idoux et al, 2025), rabbit anti-ColXII (1:400) (Bader et al, 2009), goat anti-guinea pig AlexaFLuor-488 and anti-rabbit AlexaFluor 546 (1:500, Invitrogen, France), phalloidin-647 (1:500, Invitrogen, France).

All samples were imaged using a Zeiss LSM 780 spectral confocal microscope. Whole-mount images represent single optical section.

### Protein extraction and western blot analysis

Protein extraction and Western blot analysis were described in (Idoux et al, 2025). Briefly, 3 months old fish were euthanized as indicated above. Skeletal muscle was dissected after removal of the skin and snap-frozen and store at -80°C until further use. Samples were incubated in extraction solution containing 50 mM Tris–HCl (pH 6.8), 5% glycerol, 1% SDS, 4 M urea, 50 mM DTT, and 0.1% bromophenol blue at a ratio of 1:20 (mg tissue/μL buffer) and homogenized using a mortar and pestle. After centrifugation, the supernatant was heated at 95 °C for 3 min, separated by 5% SDS–PAGE and transferred onto PVDF membranes. After blocking in skimmed milk, membranes were incubated with guinea pig anti–ColVI (1:1,000; Idoux et al, 2025) and mouse monoclonal anti-vinculin (1:8,000; V4505, Sigma-Aldrich) as a loading control. Immunoreactive bands were detected using HRP-conjugated secondary antibodies and the ImmunStar WesternC substrate (Bio-Rad). The band intensities were quantified by densitometry using ImageJ software and protein levels were normalized to vinculin.

### Cellpose segmentation

Muscle fibre cross-sectional area (CSA) was quantified from fluorescence microscopy images of WGA using an automated segmentation pipeline followed by manual quality control. Single-channel images of muscle cross-sections were used for analysis. When Z-stack images were obtained, they were first collapsed into a single plane using maximum intensity projection prior to segmentation. Images were then imported directly into Cellpose for fibre detection. Muscle fibres were segmented using the cytoplasm model (Cyto3) implemented in Cellpose. The average fibre diameter was first calibrated using the built-in diameter estimation tool, after which automated segmentation was performed. Where segmentation missed fibres or included incorrect masks, manual corrections were carried out by adding or removing regions to ensure accurate representation of individual fibres. Following segmentation, the processed segmentated masks were imported into FIJI (ImageJ) using the Labels-to-ROI plugin to generate individual ROIs for each muscle fibre. For each ROI, morphometric parameters were measured using the standard FIJI measurement settings. The primary parameter used in this study area, perimeter and Feret’s diameter. For each fish sample, multiple image frames were analyzed to minimize variability due to sectioning or imaging differences. Fibre-level measurements obtained from each frame were first compiled into a single dataset per frame. The mean value across all frames was used to represent each individual fish sample. Statistical analyses were performed using GraphPad Prism. For each experimental group, the mean CSA value calculated from frame-averaged measurements for each fish was used as a single biological data point. Data distribution was assessed prior to statistical testing. Group comparisons were performed depending on data normality. Results are presented as mean ± standard deviation (SD), and statistical significance was defined as p < 0.05.

### Drug treatment

WT and BM embryos were collected after natural mating and kept in Petri dish in E3 for 1 day at 28°C. At 24hpf, manually dechorionated embryos were transferred into 96-well plates (1 embryo per well) with 200 µl of E3, DMSO (0.125%, Sigma-Aldrich), 4-BPA (0.01 mM in E3, Sodium 4-Phenylbutyrate, Cayman chemical, 11323) TUDCA (0.1 mM in E3, Tauroursodeoxycholic Acid, Sodium Salt, Millipore, 580549) or carbamazepine (CBZ, 0.25 mM in 0.125% DMSO, Sigma-Aldrich, C4024). Half the media were replaced with fresh solution every day until 5 dpf. All treatment included PTU at 0.003% final concentration to prevent pigmentation. Larvae were then immunostained with anti-ColVI and imaged as described above.

### Statistical analysis

All statistical analyses were performed using GraphPad Prism (Version 10.2.2). Prior to statistical testing, data distribution was assessed for normality using the Shapiro–Wilk test. For comparisons involving more than two groups or two independent variables (genotype and age), a two-way analysis of variance (ANOVA) was applied, followed by Tukey’s or Šídák’s post-hoc test for multiple comparisons, as appropriate. Sex-stratified morphometric comparisons (body weight and standard length within each genotype across timepoints) were performed using one-way ANOVA with Tukey’s post-hoc correction. Geometric morphometric data were analysed in MorphoJ (Klingenberg, 2011) using Procrustes superimposition followed by Principal Component Analysis (PCA) and Canonical Variate Analysis (CVA) to assess shape variation between genotypes and timepoints. For RT-qPCR data, gene expression was normalised to the reference gene actb using the 2^−ΔΔCt^ method and group comparisons were performed as described above. For muscle fibre cross-sectional area (CSA) analysis, the mean CSA value derived from frame-averaged measurements was used as a single biological replicate per fish. Micro-CT volumetric and bone mineral density data were analysed using two-way ANOVA with post-hoc correction for multiple comparisons across genotype and age. For single-nucleus RNA sequencing, differential gene expression between conditions was performed using the Seurat FindMarkers function with the Wilcoxon rank-sum test; adjusted p-values were computed using Bonferroni correction. Gene set enrichment analysis (GSEA) was performed using the fgsea package in R, with normalised enrichment scores (NES) and adjusted p-values reported. Cell-cell communication analysis was performed using CellChat v2; differences in interaction number and strength between conditions were assessed using the built-in differential interaction analysis framework. All bar graph data are presented as mean ± standard error of the mean (SEM) unless otherwise stated. Statistical significance was defined as p < 0.05, with significance levels indicated as follows: * p < 0.05, ** p < 0.01, *** p < 0.001, **** p < 0.0001; ns = not significant.

## Supporting information

Spplemental Figures S1-S4 and Tables S1-17

## AUTHOR CONTRIBUTION

S.S.: Experiment Design, Methodology, Investigation, Formal analysis, Visualization, Writing-original draft, Writing-review & editing. L.G.: Experiment Design, Methodology, Investigation, Formal analysis, Writing-review & editing. F.S.: Experiment Design, Methodology, Investigation, Formal analysis, Writing-review & editing. A.F.: Methodology, Formal analysis, Writing-review & editing. L.L-M.: Resources, Writing-review & editing. E.D.: Experiment Design, Formal analysis, Resources, Writing-review & editing. S.B.: Experiment Design, Methodology, Investigation, Resources, Supervision, Writing-review & editing. F.R.: Conceptualization, Funding acquisition, Project administration, Supervision, Resources, Writing-original draft, Writing-review & editing. All authors read and approved the final manuscript.

## ACKNOWLEDGMENTS

We are grateful to Prof Tom van Agtmael (Univ Glasgow, Scotland) for useful discussion on drug testing. We acknowledge the contribution of the SFR Biosciences (UAR3444/CNRS, US8/Inserm, ENS de Lyon, Université Claude Bernard Lyon 1, Lyon, France) facilities, notably the zebrafish facility (PRECI, Laure Bernard and Robert Renard), the IGFL imaging platform (Marilyne Malbouyres) and the CRCL cancer genomics platform (Cyril Degletagne). We thank Arthur Gairin-Calvo for his assistance in morphometrics analysis. This study was funded by the European Union within the Horizon Europe MSCA program under grant agreement N° 101072766 (CHANGE project). SS is a recipient of a doctoral fellowship (CHANGE project) and a 4^th^ year doctoral fellowship from the FRM (Fondation pour la Recherche Médicale; FDT202504020714); RF received a financial support of Graduate Initiative MuSkLE, Lyon Saint-Etienne Universities Graduate+ project, funded by the French National Research Agency under the France 2030 program (ANR-21-SFRI-0001). The funder played no role in study design, data collection, analysis and interpretation of data, or the writing of this manuscript.

**SUPPLEMENTAL INFORMATION**

**Figures S1-S4**

**Tables S1-S7**

