## Supplementary material for "Rewiring Fibroblast–Muscle Axis Drives Progressive Pathology in Bethlem Myopathy": Spplemental Figures S1-S4 and Tables S1-17

**Supplemental Information**

Progressive Collagen VI Deficiency Disrupts the Myomatrix–Fibroblast–Muscle Trio in a Zebrafish Model of Bethlem Myopathy

Shivashakthi Shivaraman, Laurent Gilquin, Frederic Sohm, Laurence Legeais-Mallet, Antonella Forlino, Emilie Dambroise, Sandrine Bretaud* and Florence Ruggiero*

**Supplemental Figures**

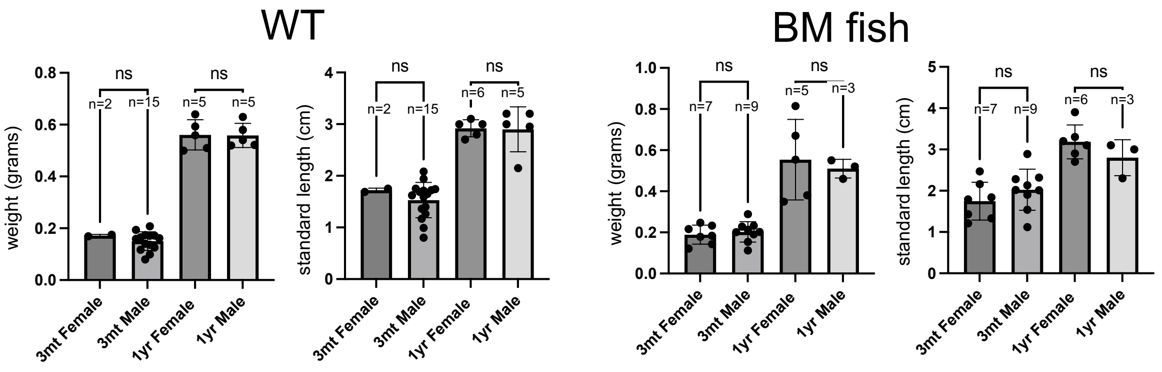

**Figure S1 related to Figure 1: Growth differences in BM fish are independent of sex.** Standard body length and body weight of male or female WT (left) and BM (right) fish at 3 month and 1 year of age. No significant sex-dependent differences were detected within either genotype at either time point. Statistical analysis was performed using ordinary one-way ANOVA test. ns: no significance. Error bars are SEM.

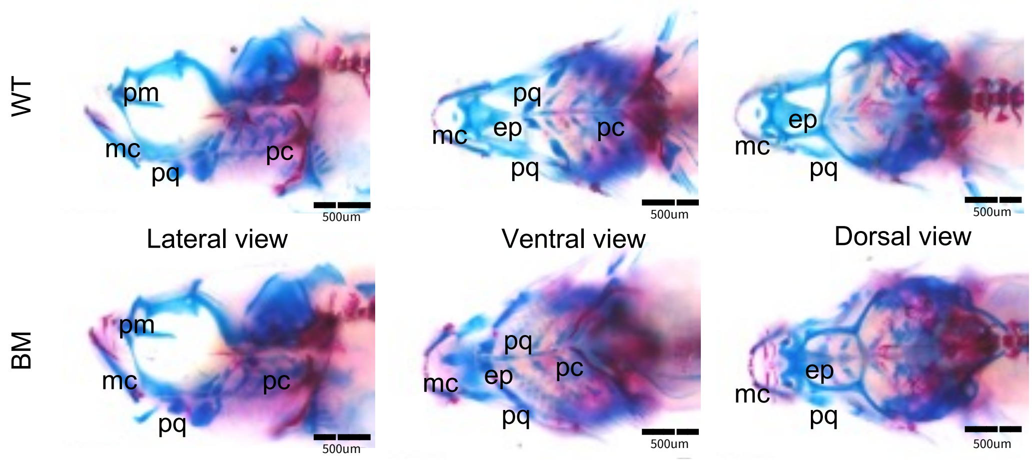

**Figure S2 related to Figure 2: The BM bone phenotype is not attributable to developmental defects.** Alizarin red (mineralized bone) and Alcian blue (cartilage) staining of WT and BM fish at 1 month. Lateral, ventral and dorsal views of the head are presented. n=4 for each condition. mc, Meckel's cartilage; pq, platoquadrate; pc, pharyngeal cartilage; ep, ethmoid plate. Scale bars = 500 µm.

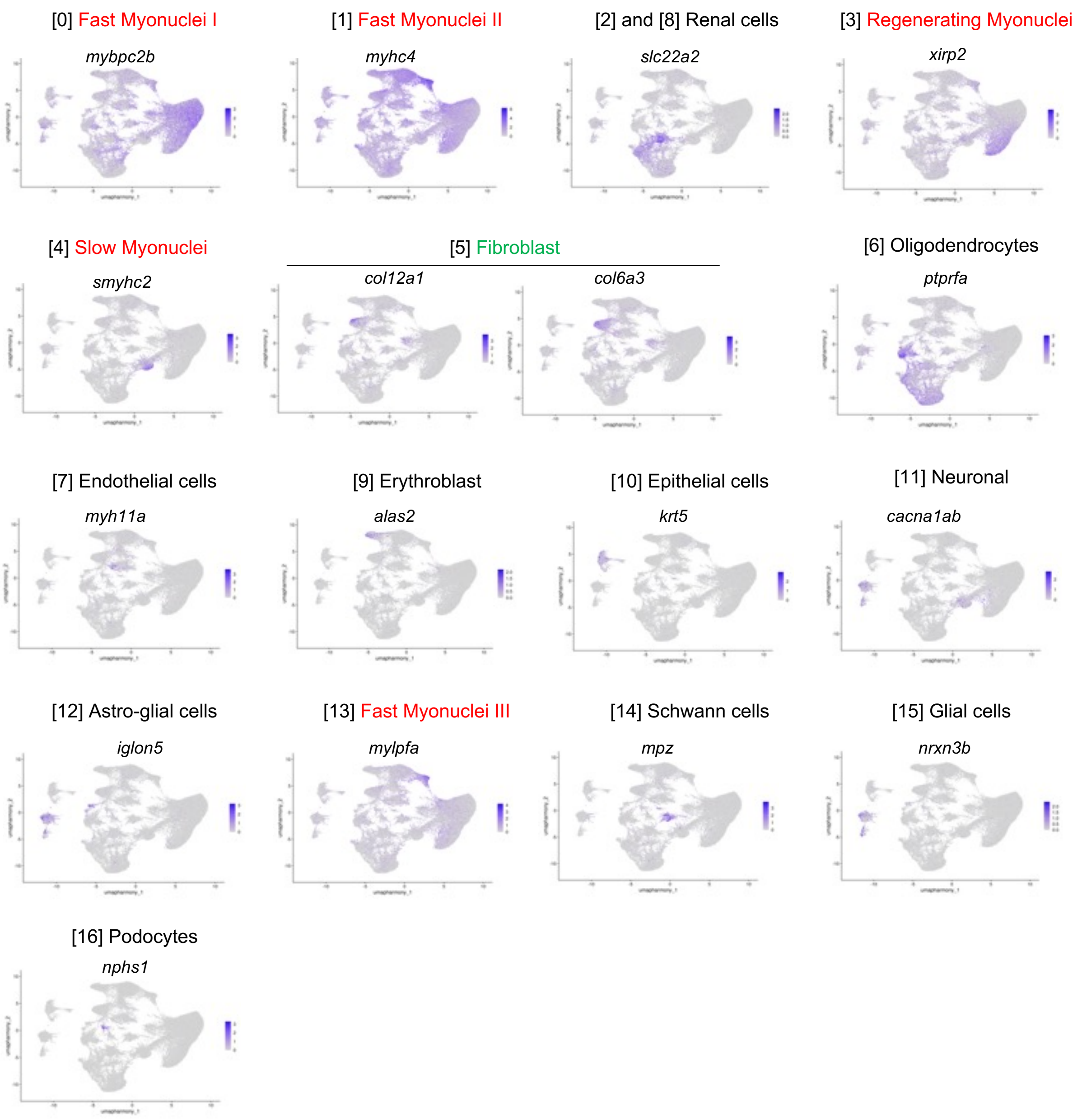

**Figure S3 related to Figure 3: Assignment of cell types and myonuclear populations to clusters 0-16 based on gene maker expression.** UMAP visualization of the expression of curated representative gene markers for each cluster (0-16). Colour gradient bar indicates the average gene expression (log-normalised expression values).

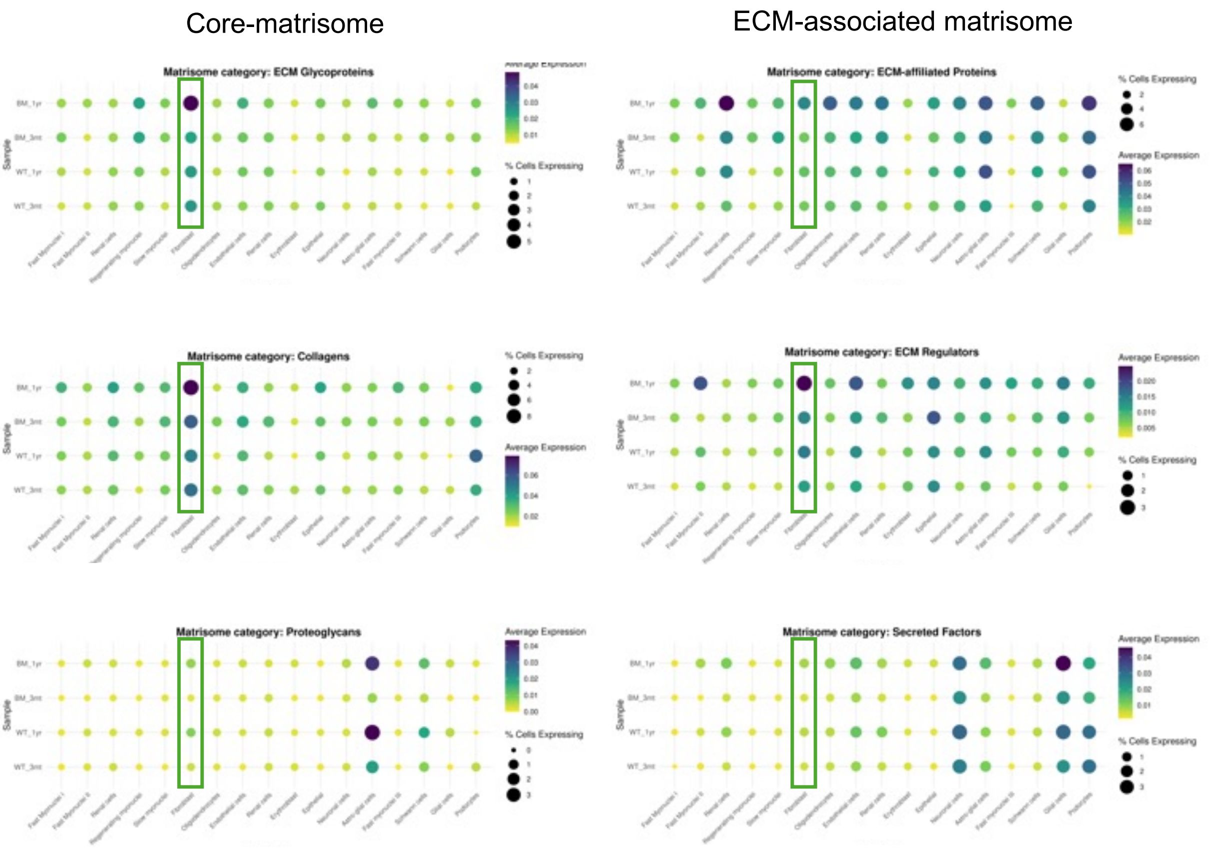

**Figure S4 related to Figure 5: Matrisome gene expression in the different clusters (0-16).** Dot plots representing matrisome gene expression across different categories (core matrisome and ECM-associated matrisome) for each cluster (0–16). Green boxes indicate cluster 5 (fibroblasts). Dot color indicates the average gene expression (log-normalized expression values) and dot size corresponds to the percentage of cells expressing the gene in each cluster.

### Supplemental Tables

**Table S1: Top marker genes used for cell type and nuclei population annotation.** Top differentially expressed marker genes identified for each cluster determined by the Seurat *FindAllMarkers* function (Wilcoxon rank-sum test). For each gene, the table reports the cluster identity, average log2 fold change, percentage of nuclei expressing the gene within the cluster (pct.1) and outside the cluster (pct.2), adjusted p-value, and the cell type identity assigned to each cluster based on marker gene expression.

| Cluster | Gene_Symbol | Avg_Log2fc | Pct.1 | Pct.2 | P_Val_Adj |
|---|---|---|---|---|---|
| Fast Myonuclei I | *pdlim5a* | 1.923798 | 0.734 | 0.307 | 0 |
| Fast Myonuclei I | *phka1b* | 1.920008 | 0.588 | 0.205 | 0 |
| Fast Myonuclei I | *UCKL1* | 1.894115 | 0.679 | 0.288 | 0 |
| Fast Myonuclei I | *CA12* | 1.889012 | 0.272 | 0.086 | 0 |
| Fast Myonuclei I | *sbk3* | 1.835288 | 0.407 | 0.154 | 0 |
| Fast Myonuclei I | *cacng6b* | 1.824051 | 0.428 | 0.142 | 0 |
| Fast Myonuclei I | *nt5c1aa* | 1.818222 | 0.669 | 0.314 | 0 |
| Fast Myonuclei I | *ampd3b* | 1.75794 | 0.316 | 0.121 | 0 |
| Fast Myonuclei I | *fgf12b* | 1.74837 | 0.274 | 0.091 | 0 |
| Fast Myonuclei I | *igfn1.1* | 1.695351 | 0.869 | 0.48 | 0 |
| Fast Myonuclei I | *CR384075.1* | 1.663995 | 0.48 | 0.184 | 0 |
| Fast Myonuclei I | *dennd4a* | 1.651078 | 0.348 | 0.131 | 0 |
| Fast Myonuclei I | *synpo* | 1.647162 | 0.331 | 0.128 | 0 |
| Fast Myonuclei I | *CHST8* | 1.637869 | 0.694 | 0.327 | 0 |
| Fast Myonuclei II | *BX548011.2* | 1.709583 | 0.661 | 0.318 | 0 |
| Fast Myonuclei II | *wu:fi09b08* | 1.674863 | 0.599 | 0.275 | 0 |
| Fast Myonuclei II | *myha* | 1.476523 | 0.416 | 0.169 | 0 |
| Fast Myonuclei II | *myhc4* | 1.177914 | 0.989 | 0.901 | 0 |
| Fast Myonuclei II | *FO704772.3* | 1.065107 | 0.572 | 0.338 | 0 |
| Fast Myonuclei II | *myhz1.1l* | 0.654027 | 0.612 | 0.396 | 0 |
| Fast Myonuclei II | *pvalb2* | 0.386172 | 0.305 | 0.192 | 6.9E-146 |
| Fast Myonuclei II | *tpma* | 0.358534 | 0.315 | 0.24 | 7.74E-58 |
| Renal cells I | *P4HA3* | 4.799312 | 0.43 | 0.044 | 0 |
| Renal cells I | *SLC34A1* | 4.612285 | 0.258 | 0.022 | 0 |
| Renal cells I | *lrp2a* | 4.497835 | 0.824 | 0.199 | 0 |
| Renal cells I | *slc5a2* | 4.461003 | 0.402 | 0.045 | 0 |
| Renal cells I | *slc22a13b* | 4.441431 | 0.285 | 0.019 | 0 |
| Renal cells I | *ucp1* | 4.415637 | 0.324 | 0.023 | 0 |
| Renal cells I | *slc6a19b* | 4.351655 | 0.335 | 0.037 | 0 |
| Renal cells I | *slc5a12* | 4.348037 | 0.429 | 0.038 | 0 |
| Renal cells I | *SLC5A10* | 4.297323 | 0.256 | 0.016 | 0 |
| Renal cells I | *rgl1* | 4.180821 | 0.755 | 0.121 | 0 |
| Renal cells I | *nrip2* | 4.16255 | 0.35 | 0.032 | 0 |
| Renal cells I | *SLC24A3* | 4.135208 | 0.448 | 0.117 | 0 |
| Renal cells I | *NBEA* | 4.133819 | 0.421 | 0.048 | 0 |

| | | | | | |
|---|---|---|---|---|---|
| Renal cells I | *cltrn* | 4.118323 | 0.6 | 0.078 | 0 |
| Regenerating myonuclei | *sparcl1* | 3.814974 | 0.284 | 0.023 | 0 |
| Regenerating myonuclei | *galntl6* | 3.585359 | 0.453 | 0.07 | 0 |
| Regenerating myonuclei | *sema5a* | 3.271487 | 0.432 | 0.075 | 0 |
| Regenerating myonuclei | *csmd1a* | 3.160562 | 0.294 | 0.055 | 0 |
| Regenerating myonuclei | *FHOD3* | 3.079508 | 0.85 | 0.254 | 0 |
| Regenerating myonuclei | *fras1* | 2.864697 | 0.684 | 0.166 | 0 |
| Regenerating myonuclei | *CDH13* | 2.733882 | 0.778 | 0.276 | 0 |
| Regenerating myonuclei | *frem3* | 2.68809 | 0.36 | 0.078 | 0 |
| Regenerating myonuclei | *CSPG4* | 2.665767 | 0.599 | 0.141 | 0 |
| Regenerating myonuclei | *runx2a* | 2.492359 | 0.322 | 0.076 | 0 |
| Regenerating myonuclei | *scamp5a* | 2.470296 | 0.257 | 0.051 | 0 |
| Regenerating myonuclei | *CLCN2* | 2.468322 | 0.289 | 0.062 | 0 |
| Regenerating myonuclei | *XIRP2* | 2.426719 | 0.773 | 0.251 | 0 |
| Regenerating myonuclei | *clstn1* | 2.389669 | 0.326 | 0.073 | 0 |
| Regenerating myonuclei | *eya1* | 2.384843 | 0.637 | 0.179 | 0 |
| Slow myonuclei | *mybpc1* | 5.092267 | 0.594 | 0.072 | 0 |
| Slow myonuclei | *mybpc3* | 4.879688 | 0.49 | 0.039 | 0 |
| Slow myonuclei | *myh7bb* | 4.817924 | 0.315 | 0.026 | 0 |
| Slow myonuclei | *atp2a2a* | 4.800437 | 0.616 | 0.08 | 0 |
| Slow myonuclei | *tnnt2e* | 4.773924 | 0.367 | 0.031 | 0 |
| Slow myonuclei | *RYR1* | 4.632211 | 0.722 | 0.065 | 0 |
| Slow myonuclei | *tnni4b.1* | 4.581993 | 0.315 | 0.025 | 0 |
| Slow myonuclei | *bpgm* | 4.512621 | 0.416 | 0.032 | 0 |
| Slow myonuclei | *cacna1sa* | 4.471207 | 0.347 | 0.019 | 0 |
| Slow myonuclei | *myh7ba* | 4.302037 | 0.645 | 0.072 | 0 |
| Slow myonuclei | *myoz2b* | 4.116171 | 0.348 | 0.024 | 0 |
| Slow myonuclei | *KCNB2* | 4.044717 | 0.45 | 0.06 | 0 |
| Slow myonuclei | *fgf11a* | 3.977147 | 0.28 | 0.032 | 0 |
| Slow myonuclei | *tpm2* | 3.695617 | 0.582 | 0.089 | 0 |
| Slow myonuclei | *smyhc2* | 3.677265 | 0.425 | 0.081 | 0 |
| Fibroblast | *cd34* | 3.856388 | 0.48 | 0.089 | 0 |
| Fibroblast | *htra1b* | 3.806448 | 0.304 | 0.047 | 0 |
| Fibroblast | *COL12A1* | 3.623206 | 0.483 | 0.111 | 0 |
| Fibroblast | *col28a1b* | 3.248436 | 0.257 | 0.037 | 0 |
| Fibroblast | *COL6A3* | 3.244884 | 0.625 | 0.137 | 0 |
| Fibroblast | *COL5A1* | 3.201068 | 0.533 | 0.098 | 0 |
| Fibroblast | *col5a2a* | 3.147793 | 0.349 | 0.056 | 0 |
| Fibroblast | *loxl2a* | 3.122151 | 0.311 | 0.062 | 0 |
| Fibroblast | *col6a1* | 3.10427 | 0.347 | 0.06 | 0 |
| Fibroblast | *COL12A1.1* | 2.962203 | 0.456 | 0.123 | 0 |
| Fibroblast | *LRP1.1* | 2.943891 | 0.41 | 0.075 | 0 |
| Fibroblast | *col11a1b* | 2.938374 | 0.284 | 0.058 | 0 |
| Fibroblast | *NHSL2* | 2.921095 | 0.412 | 0.098 | 0 |
| Fibroblast | *COL6A2* | 2.897594 | 0.279 | 0.049 | 0 |

| | | | | | |
|---|---|---|---|---|---|
| Fibroblast | *c6.1* | 2.85087 | 0.385 | 0.088 | 0 |
| Oligodendrocytes | *prdm16* | 5.054762 | 0.838 | 0.099 | 0 |
| Oligodendrocytes | *rhcgb* | 4.792307 | 0.419 | 0.029 | 0 |
| Oligodendrocytes | *cysltr1* | 4.545223 | 0.255 | 0.014 | 0 |
| Oligodendrocytes | *unc5b* | 4.517392 | 0.546 | 0.047 | 0 |
| Oligodendrocytes | *esrrb* | 4.486041 | 0.501 | 0.047 | 0 |
| Oligodendrocytes | *prdm16* | 4.300116 | 0.263 | 0.017 | 0 |
| Oligodendrocytes | *wnt9a* | 4.199362 | 0.361 | 0.037 | 0 |
| Oligodendrocytes | *dusp8a* | 3.993434 | 0.31 | 0.024 | 0 |
| Oligodendrocytes | *slc24a2* | 3.888172 | 0.256 | 0.021 | 0 |
| Oligodendrocytes | *BRSK2.1* | 3.806816 | 0.385 | 0.044 | 0 |
| Oligodendrocytes | *sstr5* | 3.547602 | 0.402 | 0.044 | 0 |
| Oligodendrocytes | *glis3* | 3.528567 | 0.301 | 0.029 | 0 |
| Oligodendrocytes | *linc-mir30e-2* | 3.492889 | 0.551 | 0.078 | 0 |
| Oligodendrocytes | *CFAP57* | 3.415802 | 0.292 | 0.034 | 0 |
| Endothelial cells | *fgd5a* | 4.855883 | 0.336 | 0.021 | 0 |
| Endothelial cells | *ptprb* | 4.670653 | 0.272 | 0.017 | 0 |
| Endothelial cells | *kdrl* | 4.636962 | 0.301 | 0.021 | 0 |
| Endothelial cells | *ldb2a* | 4.480142 | 0.289 | 0.026 | 0 |
| Endothelial cells | *cxcl12a* | 4.279899 | 0.292 | 0.032 | 0 |
| Endothelial cells | *stab1* | 4.193169 | 0.276 | 0.04 | 0 |
| Endothelial cells | *myh11a* | 3.939258 | 0.289 | 0.046 | 0 |
| Endothelial cells | *mtss1la* | 3.890429 | 0.339 | 0.039 | 0 |
| Endothelial cells | *epas1b* | 3.793748 | 0.258 | 0.033 | 0 |
| Endothelial cells | *cavin1b* | 3.469398 | 0.257 | 0.037 | 0 |
| Endothelial cells | *sema4c* | 3.1819 | 0.313 | 0.047 | 0 |
| Endothelial cells | *PREX2* | 3.077741 | 0.389 | 0.058 | 0 |
| Endothelial cells | *myo10l3* | 2.983787 | 0.25 | 0.046 | 0 |
| Endothelial cells | *spns2* | 2.922672 | 0.427 | 0.104 | 0 |
| Endothelial cells | *PLPP2* | 2.810015 | 0.257 | 0.054 | 0 |
| Renal cells II | *rap1gapa* | 5.778909 | 0.91 | 0.12 | 0 |
| Renal cells II | *AVPR2* | 5.294852 | 0.421 | 0.017 | 0 |
| Renal cells II | *hmg20a* | 5.234819 | 0.351 | 0.02 | 0 |
| Renal cells II | *slco1f2* | 5.026209 | 0.547 | 0.036 | 0 |
| Renal cells II | *abcb11a* | 4.942709 | 0.533 | 0.033 | 0 |
| Renal cells II | *ddc* | 4.877412 | 0.325 | 0.029 | 0 |
| Renal cells II | *slc6a6a* | 4.616397 | 0.794 | 0.078 | 0 |
| Renal cells II | *PDE8B* | 4.247821 | 0.487 | 0.038 | 0 |
| Renal cells II | *mfsd2al1* | 3.981226 | 0.444 | 0.042 | 0 |
| Renal cells II | *acot17* | 3.880885 | 0.558 | 0.06 | 0 |
| Renal cells II | *slc12a9* | 3.778791 | 0.262 | 0.025 | 0 |
| Renal cells II | *CEP152* | 3.721116 | 0.329 | 0.031 | 0 |
| Renal cells II | *slc2a9l1* | 3.701082 | 0.446 | 0.049 | 0 |
| Erythroblast | *hif1al2* | 5.403769 | 0.287 | 0.01 | 0 |
| Erythroblast | *si:ch73-55i23.1* | 5.035259 | 0.276 | 0.013 | 0 |

| | | | | | |
|---|---|---|---|---|---|
| Erythroblast | *si:ch211-250g4.3* | 4.934714 | 0.695 | 0.057 | 0 |
| Erythroblast | *alas2* | 4.750482 | 0.343 | 0.018 | 0 |
| Erythroblast | *nt5c2l1* | 4.744459 | 0.516 | 0.041 | 0 |
| Erythroblast | *epb41b* | 4.668019 | 0.519 | 0.034 | 0 |
| Erythroblast | *si:ch211-5k11.8* | 4.196167 | 0.294 | 0.03 | 0 |
| Erythroblast | *ank1a* | 4.194044 | 0.616 | 0.062 | 0 |
| Erythroblast | *slc4a1a* | 4.166111 | 0.427 | 0.056 | 0 |
| Erythroblast | *hbba1.1* | 4.144263 | 0.348 | 0.037 | 0 |
| Erythroblast | *ripor3* | 4.012917 | 0.263 | 0.021 | 0 |
| Erythroblast | *add2* | 3.977851 | 0.267 | 0.022 | 0 |
| Erythroblast | *si:ch211-227m13.1* | 3.90374 | 0.455 | 0.05 | 0 |
| Erythroblast | *zgc:66433* | 3.894668 | 0.393 | 0.041 | 0 |
| Erythroblast | *mibp* | 3.835322 | 0.342 | 0.036 | 0 |
| Epithelial cells | *LOC100333235* | 6.445484 | 0.382 | 0.007 | 0 |
| Epithelial cells | *CABZ01072532.1* | 6.419622 | 0.332 | 0.006 | 0 |
| Epithelial cells | *si:ch211-69b7.6* | 6.340099 | 0.556 | 0.023 | 0 |
| Epithelial cells | *pkp1b* | 6.205051 | 0.302 | 0.007 | 0 |
| Epithelial cells | *FAM83B* | 6.06302 | 0.433 | 0.01 | 0 |
| Epithelial cells | *evpla* | 6.05907 | 0.389 | 0.012 | 0 |
| Epithelial cells | *scel* | 5.97648 | 0.26 | 0.007 | 0 |
| Epithelial cells | *znf185* | 5.869225 | 0.308 | 0.009 | 0 |
| Epithelial cells | *FAT2* | 5.863304 | 0.352 | 0.009 | 0 |
| Epithelial cells | *pkp3a* | 5.840188 | 0.781 | 0.044 | 0 |
| Epithelial cells | *ppl* | 5.82925 | 0.367 | 0.012 | 0 |
| Epithelial cells | *FO681323.1* | 5.802902 | 0.286 | 0.008 | 0 |
| Epithelial cells | *ano1a* | 5.722416 | 0.42 | 0.017 | 0 |
| Epithelial cells | *dsc2l* | 5.636944 | 0.649 | 0.035 | 0 |
| Epithelial cells | *si:dkey-262k9.2* | 5.507708 | 0.449 | 0.017 | 0 |
| Neuronal | *GRID1.1* | 6.313143 | 0.65 | 0.035 | 0 |
| Neuronal | *slc12a5b* | 5.913491 | 0.291 | 0.006 | 0 |
| Neuronal | *GRIK2* | 5.68884 | 0.295 | 0.01 | 0 |
| Neuronal | *GRIN2A* | 5.615303 | 0.343 | 0.01 | 0 |
| Neuronal | *hs3st4* | 5.428693 | 0.289 | 0.014 | 0 |
| Neuronal | *cntnap2a* | 5.319932 | 0.57 | 0.037 | 0 |
| Neuronal | *kcnd2* | 5.264747 | 0.251 | 0.01 | 0 |
| Neuronal | *gria3b* | 5.175283 | 0.514 | 0.024 | 0 |
| Neuronal | *kcnc3a* | 5.17093 | 0.344 | 0.013 | 0 |
| Neuronal | *CAMTA1.1* | 5.169405 | 0.363 | 0.014 | 0 |
| Neuronal | *gria4b* | 5.158609 | 0.346 | 0.013 | 0 |
| Neuronal | *atp2b3b* | 5.141948 | 0.313 | 0.011 | 0 |
| Neuronal | *csmd2* | 5.136984 | 0.406 | 0.019 | 0 |
| Neuronal | *gfra4a* | 5.125147 | 0.305 | 0.015 | 0 |
| Neuronal | *ncam2* | 5.121916 | 0.542 | 0.033 | 0 |
| Astro-glial cells | *QKI* | 3.3894 | 0.463 | 0.111 | 0 |
| Astro-glial cells | *NPAS3* | 3.064946 | 0.352 | 0.108 | 2.2E-223 |

| | | | | | |
|---|---|---|---|---|---|
| Astro-glial cells | *iglon5* | 2.906159 | 0.303 | 0.06 | 0 |
| Astro-glial cells | *clstn2a* | 2.665591 | 0.254 | 0.093 | 7.6E-109 |
| Astro-glial cells | *ZNF536* | 2.592583 | 0.293 | 0.073 | 7.6E-239 |
| Astro-glial cells | *DOCK3* | 2.589849 | 0.262 | 0.051 | 7.2E-288 |
| Astro-glial cells | *znf385b* | 2.465495 | 0.402 | 0.195 | 1.7E-116 |
| Astro-glial cells | *ncam1a* | 2.358314 | 0.279 | 0.081 | 3.1E-177 |
| Astro-glial cells | *ntm* | 2.147852 | 0.289 | 0.088 | 3.7E-169 |
| Astro-glial cells | *LRP1.1* | 2.061413 | 0.33 | 0.089 | 2.5E-230 |
| Astro-glial cells | *NHSL2* | 1.712218 | 0.286 | 0.112 | 1.3E-102 |
| Astro-glial cells | *DDR1* | 1.621614 | 0.313 | 0.168 | 3.1E-57 |
| Astro-glial cells | *tgfbr3* | 1.540219 | 0.371 | 0.156 | 3.3E-122 |
| Astro-glial cells | *LRP1* | 1.536003 | 0.296 | 0.122 | 1.21E-95 |
| Astro-glial cells | *CSPG4* | 1.533895 | 0.333 | 0.17 | 4.32E-69 |
| Fast Myonuclei III | *pvalb2* | 3.777975 | 0.763 | 0.197 | 0 |
| Fast Myonuclei III | *nme2b.2* | 3.749106 | 0.932 | 0.332 | 0 |
| Fast Myonuclei III | *pvalb3* | 3.594343 | 0.638 | 0.148 | 0 |
| Fast Myonuclei III | *mylpfa* | 3.433775 | 0.94 | 0.383 | 0 |
| Fast Myonuclei III | *ak1* | 3.37214 | 0.279 | 0.037 | 0 |
| Fast Myonuclei III | *tnnc2.2* | 3.318753 | 0.617 | 0.115 | 0 |
| Fast Myonuclei III | *pvalb1* | 3.259007 | 0.813 | 0.258 | 0 |
| Fast Myonuclei III | *tpma* | 3.11741 | 0.815 | 0.239 | 0 |
| Fast Myonuclei III | *actc1b* | 3.093753 | 0.922 | 0.419 | 0 |
| Fast Myonuclei III | *ckmb* | 2.926528 | 0.818 | 0.261 | 0 |
| Fast Myonuclei III | *myhc4* | 2.919887 | 1 | 0.913 | 0 |
| Fast Myonuclei III | *si:ch73-367p23.2* | 2.676224 | 0.631 | 0.18 | 0 |
| Fast Myonuclei III | *gapdh* | 2.638233 | 0.481 | 0.128 | 0 |
| Fast Myonuclei III | *pvalb4* | 2.491928 | 0.712 | 0.282 | 0 |
| Fast Myonuclei III | *myhz1.1l* | 2.463208 | 0.594 | 0.426 | 9.12E-61 |
| Schwann cells | *dhrs12la* | 6.639445 | 0.413 | 0.008 | 0 |
| Schwann cells | *si:dkey-200l5.4* | 6.626664 | 0.519 | 0.014 | 0 |
| Schwann cells | *mag* | 6.467547 | 0.549 | 0.015 | 0 |
| Schwann cells | *mbpa* | 6.167375 | 0.849 | 0.045 | 0 |
| Schwann cells | *mpz* | 5.790668 | 0.852 | 0.064 | 0 |
| Schwann cells | *cldn19* | 5.575551 | 0.434 | 0.014 | 0 |
| Schwann cells | *plp1b* | 5.152513 | 0.364 | 0.015 | 0 |
| Schwann cells | *cracd* | 5.023822 | 0.358 | 0.019 | 0 |
| Schwann cells | *map4l* | 5.003555 | 0.371 | 0.02 | 0 |
| Schwann cells | *rnf220a* | 4.999433 | 0.51 | 0.035 | 0 |
| Schwann cells | *vim* | 4.815802 | 0.253 | 0.014 | 0 |
| Schwann cells | *CR848833.1* | 4.737819 | 0.556 | 0.052 | 0 |
| Schwann cells | *DSCAML1* | 4.6815 | 0.361 | 0.022 | 0 |
| Schwann cells | *nfasca* | 4.467867 | 0.466 | 0.038 | 0 |
| Schwann cells | *prom1a* | 4.464459 | 0.335 | 0.022 | 0 |
| Glial cells | *kcnt2a* | 7.510285 | 0.295 | 0.003 | 0 |
| Glial cells | *PHF24* | 6.753021 | 0.323 | 0.009 | 0 |

| | | | | | |
|---|---|---|---|---|---|
| Glial cells | *scn1laa* | 6.71844 | 0.343 | 0.006 | 0 |
| Glial cells | *cacna1ha* | 6.410928 | 0.652 | 0.025 | 0 |
| Glial cells | *kcnd3* | 6.375467 | 0.443 | 0.013 | 0 |
| Glial cells | *si:cabz01061351.1* | 6.198824 | 0.308 | 0.006 | 0 |
| Glial cells | *st8sia5* | 6.152325 | 0.34 | 0.006 | 0 |
| Glial cells | *PLCB1* | 6.145686 | 0.377 | 0.013 | 0 |
| Glial cells | *ank2a* | 5.988821 | 0.542 | 0.018 | 0 |
| Glial cells | *kcnd1* | 5.932588 | 0.367 | 0.015 | 0 |
| Glial cells | *FRMD3* | 5.675472 | 0.38 | 0.012 | 0 |
| Glial cells | *KCNAB2* | 5.513679 | 0.572 | 0.054 | 0 |
| Glial cells | *elavl4* | 5.464489 | 0.72 | 0.033 | 0 |
| Glial cells | *hpca* | 5.464341 | 0.527 | 0.04 | 0 |
| Glial cells | *cacna1ba* | 5.255507 | 0.45 | 0.018 | 0 |
| Podocytes | *nphs2* | 8.675163 | 0.386 | 0.001 | 0 |
| Podocytes | *si:ch73-205h11.1* | 8.570791 | 0.576 | 0.004 | 0 |
| Podocytes | *nphs1* | 8.447292 | 0.855 | 0.013 | 0 |
| Podocytes | *col4a3* | 8.177364 | 0.287 | 0.002 | 0 |
| Podocytes | *ptprq* | 8.174331 | 0.661 | 0.006 | 0 |
| Podocytes | *ANKDD1B* | 7.452526 | 0.319 | 0.003 | 0 |
| Podocytes | *wt1a* | 7.237547 | 0.669 | 0.011 | 0 |
| Podocytes | *grid2ipb* | 7.096753 | 0.602 | 0.01 | 0 |
| Podocytes | *tmem132e* | 7.006737 | 0.687 | 0.019 | 0 |
| Podocytes | *magi2a* | 6.934166 | 0.947 | 0.068 | 0 |
| Podocytes | *kirrel1b* | 6.544079 | 0.487 | 0.01 | 0 |
| Podocytes | *b3gat2* | 6.327025 | 0.394 | 0.008 | 0 |
| Podocytes | *lmx1bb* | 6.158109 | 0.747 | 0.028 | 0 |
| Podocytes | *CHRM3* | 5.945819 | 0.293 | 0.008 | 0 |
| Podocytes | *podxl* | 5.41412 | 0.39 | 0.012 | 0 |

**Table S2**: **Core matrisome genes (collagen, glycoproteins and proteoglycans) expressed by fibroblasts** (cluster 5) in each condition and their average expression (log-normalized expression values). Gene classification follows the MatrisomeDB v2.0 annotation framework (Naba et al., 2016).

| Gene_Symbol | Matrisome_Category | Matrisome_Division | WT-3mt | BM-3mt | WT-1yr | BM-1yr |
|---|---|---|---|---|---|---|
| *col12a1a* | Collagens | Core matrisome | 1.77570093 | 1.82972136 | 0.94480519 | 1.97291808 |
| *col6a3* | Collagens | Core matrisome | 0.87757009 | 1.40479876 | 1.47402597 | 2.28639133 |
| *col5a1* | Collagens | Core matrisome | 1.28878505 | 0.73993808 | 1.25324675 | 1.06702776 |
| *col1a1b* | Collagens | Core matrisome | 0.89158879 | 0.5247678 | 1.14285714 | 0.59783345 |
| *col4a2* | Collagens | Core matrisome | 1.01495327 | 0.44195046 | 0.5487013 | 0.56872038 |
| *col1a1a* | Collagens | Core matrisome | 0.90280374 | 0.38931889 | 0.86363636 | 0.4041977 |
| *col1a2* | Collagens | Core matrisome | 0.68504673 | 0.35294118 | 0.7987013 | 0.48138118 |
| *col5a2a* | Collagens | Core matrisome | 0.50373832 | 0.35758514 | 0.22727273 | 0.80162492 |
| *col11a1b* | Collagens | Core matrisome | 0.57570093 | 0.48839009 | 0.28246753 | 0.48882871 |
| *col5a3a* | Collagens | Core matrisome | 0.3046729 | 0.19736842 | 0.21428571 | 0.74678402 |
| *col6a2* | Collagens | Core matrisome | 0.32242991 | 0.30185759 | 0.21753247 | 0.54705484 |
| *col28a1b* | Collagens | Core matrisome | 0.42429907 | 0.23142415 | 0.18506494 | 0.50914015 |
| *col4a1* | Collagens | Core matrisome | 0.42990654 | 0.1493808 | 0.35714286 | 0.25592417 |
| *col27a1b* | Collagens | Core matrisome | 0.21588785 | 0.28405573 | 0.11038961 | 0.23899797 |
| *col18a1* | Collagens | Core matrisome | 0.08037383 | 0.12306502 | 0.04545455 | 0.22884225 |
| *col19a1* | Collagens | Core matrisome | 0.12523364 | 0.06656347 | 0.13311688 | 0.06838186 |
| *col8a2* | Collagens | Core matrisome | 0.09626168 | 0.0503096 | 0.09090909 | 0.15098172 |
| *col11a2* | Collagens | Core matrisome | 0.12056075 | 0.03095975 | 0.10714286 | 0.01895735 |
| *col10a1b* | Collagens | Core matrisome | 0.04859813 | 0.00696594 | 0.20454545 | 0.00067705 |
| *col14a1a* | Collagens | Core matrisome | 0.04859813 | 0.0503096 | 0.08441558 | 0.03926879 |
| *col10a1a* | Collagens | Core matrisome | 0.0317757 | 0.00696594 | 0.10714286 | 0.00541638 |
| *col11a1a* | Collagens | Core matrisome | 0.03551402 | 0.01780186 | 0.07792208 | 0.01963439 |
| *col8a1b* | Collagens | Core matrisome | 0.03084112 | 0.01625387 | 0.02272727 | 0.06770481 |
| *col2a1b* | Collagens | Core matrisome | 0.03271028 | 0.02708978 | 0.01948052 | 0.03249831 |
| *col15a1b* | Collagens | Core matrisome | 0.01214953 | 0.01780186 | 0.00974026 | 0.02979012 |
| *col4a5* | Collagens | Core matrisome | 0.01495327 | 0.01006192 | 0.01948052 | 0.02437373 |
| *col9a2* | Collagens | Core matrisome | 0.01495327 | 0.00619195 | 0.01298701 | 0.01286391 |
| *col2a1a* | Collagens | Core matrisome | 0.00841121 | 0.00541796 | 0.01298701 | 0.01015572 |
| *col9a1b* | Collagens | Core matrisome | 0.0046729 | 0.00077399 | 0.00324675 | 0.00270819 |
| *col9a3* | Collagens | Core matrisome | 0.00093458 | 0.00309598 | 0.00324675 | 0.00203114 |
| *col5a3b* | Collagens | Core matrisome | 0.00093458 | 0.00386997 | 0 | 0.00203114 |
| *col4a4* | Collagens | Core matrisome | 0 | 0 | 0.00324675 | 0.00270819 |
| *col9a1a* | Collagens | Core matrisome | 0.00093458 | 0 | 0.00324675 | 0.00067705 |
| *col28a2a* | Collagens | Core matrisome | 0.00093458 | 0.00077399 | 0 | 0.00270819 |
| *col17a1a* | Collagens | Core matrisome | 0.00093458 | 0 | 0 | 0.00067705 |
| *crim1* | ECM Glycoproteins | Core matrisome | 0.89439252 | 0.37074303 | 0.41233766 | 0.61340555 |
| *lamc1* | ECM Glycoproteins | Core matrisome | 0.5588785 | 0.54411765 | 0.3538961 | 0.6844956 |
| *agrn* | ECM Glycoproteins | Core matrisome | 0.40280374 | 0.45201238 | 0.55194805 | 0.49898443 |
| *abi3bpb* | ECM Glycoproteins | Core matrisome | 0.35046729 | 0.86068111 | 0.10064935 | 0.47935003 |

| | | | | | | |
|---|---|---|---|---|---|---|
| *slit3* | ECM Glycoproteins | Core matrisome | 0.18130841 | 0.24922601 | 0.45779221 | 0.77454299 |
| *fn1b* | ECM Glycoproteins | Core matrisome | 0.23551402 | 0.34365325 | 0.12012987 | 0.88219364 |
| *edil3a* | ECM Glycoproteins | Core matrisome | 0.34766355 | 0.32430341 | 0.24675325 | 0.63303995 |
| *postnb* | ECM Glycoproteins | Core matrisome | 0.32523364 | 0.21826625 | 0.25 | 0.71496276 |
| *igfbp5b* | ECM Glycoproteins | Core matrisome | 0.24392523 | 0.1501548 | 0.25 | 0.39471903 |
| *sparc* | ECM Glycoproteins | Core matrisome | 0.2271028 | 0.13157895 | 0.27922078 | 0.21665538 |
| *pxdn* | ECM Glycoproteins | Core matrisome | 0.20093458 | 0.19504644 | 0.17532468 | 0.27014218 |
| *fbln2* | ECM Glycoproteins | Core matrisome | 0.18411215 | 0.08204334 | 0.17857143 | 0.35951253 |
| *crispld2* | ECM Glycoproteins | Core matrisome | 0.15700935 | 0.19659443 | 0.12012987 | 0.2972241 |
| *slit2* | ECM Glycoproteins | Core matrisome | 0.21588785 | 0.23606811 | 0.14285714 | 0.14624238 |
| *ltbp1* | ECM Glycoproteins | Core matrisome | 0.23738318 | 0.16795666 | 0.09090909 | 0.2281652 |
| *fn1a* | ECM Glycoproteins | Core matrisome | 0.10841121 | 0.17569659 | 0.05519481 | 0.36086662 |
| *npnt* | ECM Glycoproteins | Core matrisome | 0.2 | 0.12306502 | 0.1038961 | 0.25118483 |
| *lamb1b* | ECM Glycoproteins | Core matrisome | 0.13271028 | 0.10681115 | 0.11688312 | 0.32092079 |
| *fras1* | ECM Glycoproteins | Core matrisome | 0.21962617 | 0.26083591 | 0.08441558 | 0.0690589 |
| *cyr61* | ECM Glycoproteins | Core matrisome | 0.11214953 | 0.1493808 | 0.0974026 | 0.27285037 |
| *svep1* | ECM Glycoproteins | Core matrisome | 0.14018692 | 0.12848297 | 0.10714286 | 0.24576845 |
| *thbs3a* | ECM Glycoproteins | Core matrisome | 0.1364486 | 0.10835913 | 0.05519481 | 0.24712255 |
| *thbs4b* | ECM Glycoproteins | Core matrisome | 0.22242991 | 0.06656347 | 0.18831169 | 0.03317536 |
| *fndc1* | ECM Glycoproteins | Core matrisome | 0.13457944 | 0.11532508 | 0.05844156 | 0.15301286 |
| *igfbp3* | ECM Glycoproteins | Core matrisome | 0.17663551 | 0.11996904 | 0.09415584 | 0.06770481 |
| *nid1a* | ECM Glycoproteins | Core matrisome | 0.10186916 | 0.05340557 | 0.07792208 | 0.20853081 |
| *ntn1a* | ECM Glycoproteins | Core matrisome | 0.04859813 | 0.08359133 | 0.02597403 | 0.2647258 |
| *cilp2* | ECM Glycoproteins | Core matrisome | 0.19158879 | 0.05572755 | 0.10714286 | 0.06431957 |
| *colq* | ECM Glycoproteins | Core matrisome | 0.09252336 | 0.06811146 | 0.10064935 | 0.13879485 |
| *lama4* | ECM Glycoproteins | Core matrisome | 0.09252336 | 0.08978328 | 0.07467532 | 0.12525389 |
| *emilin2a* | ECM Glycoproteins | Core matrisome | 0.02990654 | 0.05650155 | 0.05194805 | 0.2281652 |
| *mfge8b* | ECM Glycoproteins | Core matrisome | 0.14672897 | 0.06578947 | 0.06168831 | 0.09207854 |
| *paplnb* | ECM Glycoproteins | Core matrisome | 0.04392523 | 0.00154799 | 0.31493506 | 0.00203114 |
| *cilp* | ECM Glycoproteins | Core matrisome | 0.08598131 | 0.08591331 | 0.02922078 | 0.15030467 |
| *fbn2b* | ECM Glycoproteins | Core matrisome | 0.06448598 | 0.09055728 | 0.07467532 | 0.11035884 |
| *lamb2* | ECM Glycoproteins | Core matrisome | 0.07196262 | 0.11609907 | 0.04220779 | 0.07244414 |
| *aebp1* | ECM Glycoproteins | Core matrisome | 0.03084112 | 0.04876161 | 0.04220779 | 0.15233582 |
| *postna* | ECM Glycoproteins | Core matrisome | 0.03738318 | 0.07585139 | 0.01948052 | 0.13337847 |
| *spon1b* | ECM Glycoproteins | Core matrisome | 0.04672897 | 0.04798762 | 0.08441558 | 0.04468517 |
| *tnc* | ECM Glycoproteins | Core matrisome | 0.10841121 | 0.03637771 | 0.06168831 | 0.01083277 |
| *mfap5* | ECM Glycoproteins | Core matrisome | 0.04579439 | 0.02786378 | 0.03571429 | 0.09884902 |
| *nid2a* | ECM Glycoproteins | Core matrisome | 0.05607477 | 0.03328173 | 0.02597403 | 0.07921462 |
| *lama5* | ECM Glycoproteins | Core matrisome | 0.02616822 | 0.04566563 | 0.03571429 | 0.07718348 |
| *coch* | ECM Glycoproteins | Core matrisome | 0.0271028 | 0.00773994 | 0.13311688 | 0.01354096 |
| *vwa1* | ECM Glycoproteins | Core matrisome | 0.03084112 | 0.04566563 | 0.02272727 | 0.08056872 |
| *igsf10* | ECM Glycoproteins | Core matrisome | 0.02523364 | 0.03637771 | 0.01623377 | 0.08937035 |
| *tinagl1* | ECM Glycoproteins | Core matrisome | 0.04485981 | 0.0247678 | 0.02922078 | 0.06635071 |
| *pcolce2b* | ECM Glycoproteins | Core matrisome | 0.04018692 | 0.05572755 | 0.01948052 | 0.04536222 |
| *elnb* | ECM Glycoproteins | Core matrisome | 0.0046729 | 0.01702786 | 0.00324675 | 0.13540961 |

| | | | | | | |
|---|---|---|---|---|---|---|
| *lamb1a* | ECM Glycoproteins | Core matrisome | 0.06448598 | 0.02631579 | 0.03571429 | 0.02031144 |
| *thbs1a* | ECM Glycoproteins | Core matrisome | 0.02242991 | 0.01547988 | 0.02272727 | 0.08327691 |
| *smoc1* | ECM Glycoproteins | Core matrisome | 0.02990654 | 0.02708978 | 0.02922078 | 0.05754909 |
| *anos1b* | ECM Glycoproteins | Core matrisome | 0.0635514 | 0.02321981 | 0.02922078 | 0.01963439 |
| *spp1* | ECM Glycoproteins | Core matrisome | 0.03738318 | 0.02089783 | 0.06168831 | 0.01557211 |
| *rspo1* | ECM Glycoproteins | Core matrisome | 0.01962617 | 0.00851393 | 0.01948052 | 0.07718348 |
| *ctgfb* | ECM Glycoproteins | Core matrisome | 0.05420561 | 0.01934985 | 0.01298701 | 0.02301963 |
| *thbs4a* | ECM Glycoproteins | Core matrisome | 0.02336449 | 0.02167183 | 0.02272727 | 0.04129993 |
| *spon1a* | ECM Glycoproteins | Core matrisome | 0.03457944 | 0.01857585 | 0.02597403 | 0.02979012 |
| *spon2a* | ECM Glycoproteins | Core matrisome | 0.08037383 | 0.02244582 | 0 | 0.00541638 |
| *rspo2* | ECM Glycoproteins | Core matrisome | 0.05233645 | 0.01780186 | 0.00974026 | 0.02775897 |
| *ntn1b* | ECM Glycoproteins | Core matrisome | 0.01214953 | 0.02244582 | 0.01948052 | 0.0534868 |
| *ctgfa* | ECM Glycoproteins | Core matrisome | 0.03364486 | 0.01547988 | 0.02922078 | 0.02911307 |
| *igfbp2b* | ECM Glycoproteins | Core matrisome | 0.02242991 | 0.02321981 | 0.00324675 | 0.05484089 |
| *sparcl1* | ECM Glycoproteins | Core matrisome | 0.04579439 | 0.04179567 | 0.00649351 | 0.00473934 |
| *ndnf* | ECM Glycoproteins | Core matrisome | 0.00934579 | 0.01160991 | 0.06818182 | 0.00338524 |
| *smoc2* | ECM Glycoproteins | Core matrisome | 0.02242991 | 0.01625387 | 0.01623377 | 0.03452945 |
| *bmper* | ECM Glycoproteins | Core matrisome | 0.02429907 | 0.01470588 | 0.02922078 | 0.02031144 |
| *spon2b* | ECM Glycoproteins | Core matrisome | 0.01682243 | 0.01702786 | 0.01298701 | 0.04062288 |
| *srpx* | ECM Glycoproteins | Core matrisome | 0.02616822 | 0.01702786 | 0.00974026 | 0.03182126 |
| *fgl2a* | ECM Glycoproteins | Core matrisome | 0.01588785 | 0.00851393 | 0.04220779 | 0.01624915 |
| *igfbp1a* | ECM Glycoproteins | Core matrisome | 0.01775701 | 0.01857585 | 0.02597403 | 0.01895735 |
| *tnr* | ECM Glycoproteins | Core matrisome | 0.01588785 | 0.01780186 | 0.01623377 | 0.02640487 |
| *dpt* | ECM Glycoproteins | Core matrisome | 0.00934579 | 0.00851393 | 0.03246753 | 0.02505078 |
| *efemp2a* | ECM Glycoproteins | Core matrisome | 0.01869159 | 0.01160991 | 0.01298701 | 0.02572783 |
| *mgp* | ECM Glycoproteins | Core matrisome | 0.02056075 | 0.00541796 | 0.01623377 | 0.02640487 |
| *bglapl* | ECM Glycoproteins | Core matrisome | 0.0364486 | 0.01083591 | 0.01298701 | 0.00270819 |
| *crispld1a* | ECM Glycoproteins | Core matrisome | 0.01308411 | 0.00773994 | 0.03246753 | 0.00947867 |
| *lgi3* | ECM Glycoproteins | Core matrisome | 0.02149533 | 0.00773994 | 0.01623377 | 0.01624915 |
| *fbln1* | ECM Glycoproteins | Core matrisome | 0.01121495 | 0.00309598 | 0.00974026 | 0.03723764 |
| *bglap* | ECM Glycoproteins | Core matrisome | 0.01869159 | 0.00696594 | 0.02272727 | 0.01286391 |
| *rspo3* | ECM Glycoproteins | Core matrisome | 0.00280374 | 0.00851393 | 0 | 0.04874746 |
| *mmrn2a* | ECM Glycoproteins | Core matrisome | 0.00654206 | 0.0123839 | 0.02272727 | 0.0182803 |
| *slit1a* | ECM Glycoproteins | Core matrisome | 0.01962617 | 0.0123839 | 0.01298701 | 0.01489506 |
| *mxra5a* | ECM Glycoproteins | Core matrisome | 0.01028037 | 0.00541796 | 0.02597403 | 0.0169262 |
| *anos1a* | ECM Glycoproteins | Core matrisome | 0.00747664 | 0.00464396 | 0.01948052 | 0.02505078 |
| *pcolceb* | ECM Glycoproteins | Core matrisome | 0.01401869 | 0.00851393 | 0.02597403 | 0.00812458 |
| *tgfbi* | ECM Glycoproteins | Core matrisome | 0.01775701 | 0.01780186 | 0.00649351 | 0.01421801 |
| *ddx26b* | ECM Glycoproteins | Core matrisome | 0.01121495 | 0.0123839 | 0.00974026 | 0.02234259 |
| *gas6* | ECM Glycoproteins | Core matrisome | 0.01401869 | 0.00928793 | 0.00974026 | 0.02098849 |
| *srpx2* | ECM Glycoproteins | Core matrisome | 0.02242991 | 0.00696594 | 0 | 0.02234259 |
| *mfap1* | ECM Glycoproteins | Core matrisome | 0.01214953 | 0.00773994 | 0.01298701 | 0.01421801 |
| *wisp1a* | ECM Glycoproteins | Core matrisome | 0.00747664 | 0.00154799 | 0.03571429 | 0.0013541 |
| *matn3a* | ECM Glycoproteins | Core matrisome | 0.0046729 | 0.00851393 | 0.01623377 | 0.01624915 |
| *tnfaip6* | ECM Glycoproteins | Core matrisome | 0.00280374 | 0.00773994 | 0.01623377 | 0.01760325 |

| | | | | | | |
|---|---|---|---|---|---|---|
| *cthrc1a* | ECM Glycoproteins | Core matrisome | 0.0046729 | 0.00232198 | 0.02272727 | 0.01421801 |
| *tnw* | ECM Glycoproteins | Core matrisome | 0.01495327 | 0.00154799 | 0.01623377 | 0.01083277 |
| *igfbp7* | ECM Glycoproteins | Core matrisome | 0.00934579 | 0.01006192 | 0.00974026 | 0.01354096 |
| *matn4* | ECM Glycoproteins | Core matrisome | 0.01121495 | 0.0123839 | 0.00324675 | 0.01557211 |
| *comp* | ECM Glycoproteins | Core matrisome | 0.00934579 | 0.01470588 | 0 | 0.01624915 |
| *ntng1a* | ECM Glycoproteins | Core matrisome | 0.00747664 | 0.00077399 | 0.00974026 | 0.02166554 |
| *tsku* | ECM Glycoproteins | Core matrisome | 0.0046729 | 0.01315789 | 0.00649351 | 0.01286391 |
| *mfge8a* | ECM Glycoproteins | Core matrisome | 0.01121495 | 0.00464396 | 0.00649351 | 0.01421801 |
| *gldn* | ECM Glycoproteins | Core matrisome | 0.01401869 | 0.00232198 | 0.00649351 | 0.01150982 |
| *ntng2a* | ECM Glycoproteins | Core matrisome | 0.00934579 | 0.00541796 | 0.00649351 | 0.01286391 |
| *efemp2b* | ECM Glycoproteins | Core matrisome | 0.00654206 | 0.00696594 | 0.00324675 | 0.0169262 |
| *pcolcea* | ECM Glycoproteins | Core matrisome | 0.00747664 | 0.00386997 | 0.00649351 | 0.01489506 |
| *cthrc1b* | ECM Glycoproteins | Core matrisome | 0.00747664 | 0.00928793 | 0.00974026 | 0.00609343 |
| *fbn2a* | ECM Glycoproteins | Core matrisome | 0.00280374 | 0.00464396 | 0.01948052 | 0.00541638 |
| *VWA5A (1 to many)* | ECM Glycoproteins | Core matrisome | 0.00373832 | 0.00851393 | 0.00974026 | 0.01015572 |
| *creld2* | ECM Glycoproteins | Core matrisome | 0.0046729 | 0.00464396 | 0.00649351 | 0.01489506 |
| *otog* | ECM Glycoproteins | Core matrisome | 0.0046729 | 0.00232198 | 0.00649351 | 0.0169262 |
| *ch1073-291c23.1* | ECM Glycoproteins | Core matrisome | 0.00841121 | 0.00619195 | 0.00324675 | 0.01150982 |
| *matn3b* | ECM Glycoproteins | Core matrisome | 0.02242991 | 0.00154799 | 0.00324675 | 0.00203114 |
| *lamb4* | ECM Glycoproteins | Core matrisome | 0.01401869 | 0.00309598 | 0.00974026 | 0.0013541 |
| *emilin1a* | ECM Glycoproteins | Core matrisome | 0.00373832 | 0.00154799 | 0.00974026 | 0.01083277 |
| *wisp1b* | ECM Glycoproteins | Core matrisome | 0.01495327 | 0.00154799 | 0.00324675 | 0.00541638 |
| *tecta* | ECM Glycoproteins | Core matrisome | 0.00654206 | 0.00309598 | 0.00974026 | 0.00473934 |
| *fbln5* | ECM Glycoproteins | Core matrisome | 0.00560748 | 0.00464396 | 0.00649351 | 0.00609343 |
| *vwf* | ECM Glycoproteins | Core matrisome | 0.00560748 | 0.00464396 | 0 | 0.01083277 |
| *igfbp1b* | ECM Glycoproteins | Core matrisome | 0.00841121 | 0.00232198 | 0.00324675 | 0.00677048 |
| *wisp2* | ECM Glycoproteins | Core matrisome | 0.00280374 | 0.00309598 | 0 | 0.01354096 |
| *matn1* | ECM Glycoproteins | Core matrisome | 0.00186916 | 0.00309598 | 0.00974026 | 0.00067705 |
| *igfbp2a* | ECM Glycoproteins | Core matrisome | 0.00280374 | 0.00077399 | 0.00649351 | 0.00473934 |
| *slit1b* | ECM Glycoproteins | Core matrisome | 0.0046729 | 0.00232198 | 0.00324675 | 0.00270819 |
| *cyr61l2* | ECM Glycoproteins | Core matrisome | 0.0046729 | 0.00232198 | 0.00324675 | 0.00270819 |
| *ntn4* | ECM Glycoproteins | Core matrisome | 0 | 0.00154799 | 0.00974026 | 0.00067705 |
| *sbspon* | ECM Glycoproteins | Core matrisome | 0 | 0.00309598 | 0.00649351 | 0.0013541 |
| *lama1* | ECM Glycoproteins | Core matrisome | 0 | 0.00541796 | 0.00324675 | 0.0013541 |
| *vwa9* | ECM Glycoproteins | Core matrisome | 0.00093458 | 0.00154799 | 0.00324675 | 0.00338524 |
| *tectb* | ECM Glycoproteins | Core matrisome | 0 | 0.00773994 | 0 | 0 |
| *igfbp6a* | ECM Glycoproteins | Core matrisome | 0.00280374 | 0.00154799 | 0 | 0.00338524 |
| *adipoqa* | ECM Glycoproteins | Core matrisome | 0.00093458 | 0.00309598 | 0.00324675 | 0 |
| *nell2a* | ECM Glycoproteins | Core matrisome | 0.00093458 | 0.00154799 | 0 | 0.00406229 |
| *vwde* | ECM Glycoproteins | Core matrisome | 0.00373832 | 0.00077399 | 0 | 0.00203114 |
| *thbs3b* | ECM Glycoproteins | Core matrisome | 0.00093458 | 0.00077399 | 0.00324675 | 0.0013541 |
| *fgb* | ECM Glycoproteins | Core matrisome | 0 | 0.00077399 | 0.00324675 | 0.0013541 |
| *zp3c* | ECM Glycoproteins | Core matrisome | 0 | 0.00077399 | 0.00324675 | 0.0013541 |
| *lgi2a* | ECM Glycoproteins | Core matrisome | 0.00093458 | 0 | 0.00324675 | 0.00067705 |

| | | | | | | |
|---|---|---|---|---|---|---|
| *lgi1a* | ECM Glycoproteins | Core matrisome | 0.00186916 | 0.00154799 | 0 | 0.0013541 |
| *igfals* | ECM Glycoproteins | Core matrisome | 0.00093458 | 0.00077399 | 0 | 0.00270819 |
| *vwa2* | ECM Glycoproteins | Core matrisome | 0.00280374 | 0.00077399 | 0 | 0.00067705 |
| *otol1a* | ECM Glycoproteins | Core matrisome | 0 | 0 | 0.00324675 | 0.00067705 |
| *adipoqb* | ECM Glycoproteins | Core matrisome | 0.00093458 | 0.00154799 | 0 | 0.0013541 |
| *creld1* | ECM Glycoproteins | Core matrisome | 0 | 0.00232198 | 0 | 0.0013541 |
| *mfap4* | ECM Glycoproteins | Core matrisome | 0 | 0.00077399 | 0 | 0.00270819 |
| *igfbp5a* | ECM Glycoproteins | Core matrisome | 0.00186916 | 0.00154799 | 0 | 0 |
| *igfbp6b* | ECM Glycoproteins | Core matrisome | 0.00186916 | 0.00077399 | 0 | 0.00067705 |
| *tspeara* | ECM Glycoproteins | Core matrisome | 0 | 0.00232198 | 0 | 0.00067705 |
| *cyr61l1* | ECM Glycoproteins | Core matrisome | 0 | 0 | 0 | 0.00203114 |
| *lgi2b* | ECM Glycoproteins | Core matrisome | 0.00186916 | 0 | 0 | 0 |
| *egflam* | ECM Glycoproteins | Core matrisome | 0.00093458 | 0.00077399 | 0 | 0 |
| *efemp1* | ECM Glycoproteins | Core matrisome | 0.00093458 | 0 | 0 | 0.00067705 |
| *pcolce2a* | ECM Glycoproteins | Core matrisome | 0.00093458 | 0 | 0 | 0.00067705 |
| *zp3b* | ECM Glycoproteins | Core matrisome | 0.00093458 | 0 | 0 | 0 |
| *lgi1b* | ECM Glycoproteins | Core matrisome | 0 | 0.00077399 | 0 | 0 |
| *rspo4* | ECM Glycoproteins | Core matrisome | 0 | 0.00077399 | 0 | 0 |
| *hspg2* | Proteoglycans | Core matrisome | 0.50747664 | 0.52244582 | 0.40584416 | 0.58090724 |
| *dcn* | Proteoglycans | Core matrisome | 0.29906542 | 0.15402477 | 0.1525974 | 0.34326337 |
| *vcanb* | Proteoglycans | Core matrisome | 0.10747664 | 0.23142415 | 0.12662338 | 0.3324306 |
| *hapln1a* | Proteoglycans | Core matrisome | 0.17009346 | 0.11842105 | 0.0974026 | 0.22477996 |
| *bgna* | Proteoglycans | Core matrisome | 0.0635514 | 0.01780186 | 0.14935065 | 0.01354096 |
| *prelp* | Proteoglycans | Core matrisome | 0.05233645 | 0.03482972 | 0.04545455 | 0.09884902 |
| *prg4b* | Proteoglycans | Core matrisome | 0.0317757 | 0.01083591 | 0.06818182 | 0.09952607 |
| *bgnb* | Proteoglycans | Core matrisome | 0.03271028 | 0.02863777 | 0.0487013 | 0.04739336 |
| *podn* | Proteoglycans | Core matrisome | 0.01869159 | 0.01702786 | 0.01298701 | 0.07786053 |
| *bcan* | Proteoglycans | Core matrisome | 0.03925234 | 0.01547988 | 0.05194805 | 0.01354096 |
| *chad* | Proteoglycans | Core matrisome | 0.01495327 | 0.00386997 | 0.07467532 | 0.00744753 |
| *spock1* | Proteoglycans | Core matrisome | 0.02523364 | 0.02089783 | 0.00974026 | 0.02708192 |
| *aspn* | Proteoglycans | Core matrisome | 0.02616822 | 0.01393189 | 0.00649351 | 0.02031144 |
| *srgn* | Proteoglycans | Core matrisome | 0.00560748 | 0.01857585 | 0.02272727 | 0.01963439 |
| *hapln1b* | Proteoglycans | Core matrisome | 0.00280374 | 0.00541796 | 0.03246753 | 0.01421801 |
| *lum* | Proteoglycans | Core matrisome | 0.01495327 | 0.0123839 | 0.01623377 | 0.01015572 |
| *spock3* | Proteoglycans | Core matrisome | 0.01028037 | 0.01780186 | 0.00974026 | 0.01557211 |
| *ncana* | Proteoglycans | Core matrisome | 0.00747664 | 0.00464396 | 0.00649351 | 0.00270819 |
| *ogn* | Proteoglycans | Core matrisome | 0.00560748 | 0.00154799 | 0 | 0.00744753 |
| *hapln3* | Proteoglycans | Core matrisome | 0.00280374 | 0.00386997 | 0 | 0.00609343 |
| *spock2* | Proteoglycans | Core matrisome | 0.00280374 | 0.00386997 | 0.00324675 | 0.00270819 |
| *omd* | Proteoglycans | Core matrisome | 0.00186916 | 0.00077399 | 0.00324675 | 0.00203114 |
| *prg4a* | Proteoglycans | Core matrisome | 0 | 0.00464396 | 0 | 0.0013541 |
| *hapln2* | Proteoglycans | Core matrisome | 0.00280374 | 0 | 0 | 0.00203114 |
| *chadlb* | Proteoglycans | Core matrisome | 0 | 0 | 0.00324675 | 0.00067705 |
| *fmodb* | Proteoglycans | Core matrisome | 0 | 0.00386997 | 0 | 0 |
| *epyc* | Proteoglycans | Core matrisome | 0 | 0.00077399 | 0 | 0.00203114 |

| | | | | | | |
|---|---|---|---|---|---|---|
| *esm1* | Proteoglycans | Core matrisome | 0.00093458 | 0 | 0 | 0.00067705 |
| *fmoda* | Proteoglycans | Core matrisome | 0.00093458 | 0 | 0 | 0 |
| *kera* | Proteoglycans | Core matrisome | 0.00093458 | 0 | 0 | 0 |

**Table S3**: **ECM regulators matrisome genes expressed by fibroblasts** (cluster 5) in each condition and their average expression (log-normalized expression values). Gene classifiction follows the v2.0 annotation framework (Naba et al., 2016).

| Gene_Symbol | Matrisome_Category | Matrisome_Division | Wt-3mt | BM-3mt | Wt-1yr | BM-1yr |
|---|---|---|---|---|---|---|
| *htra1b* | ECM Regulators | Matrisome-associated | 0.28598131 | 0.44117647 | 0.38636364 | 1.5788761 |
| *loxl2a* | ECM Regulators | Matrisome-associated | 0.64953271 | 0.63622291 | 0.43181818 | 0.67433988 |
| *sulf1* | ECM Regulators | Matrisome-associated | 0.2364486 | 0.22678019 | 0.16558442 | 0.94854435 |
| *adamtsl3* | ECM Regulators | Matrisome-associated | 0.25140187 | 0.2755418 | 0.18506494 | 0.57075152 |
| *bmp1a* | ECM Regulators | Matrisome-associated | 0.40841121 | 0.27708978 | 0.23701299 | 0.35003385 |
| *egln2* | ECM Regulators | Matrisome-associated | 0.70373832 | 0.15325077 | 0.27922078 | 0.12660799 |
| *fam20ca* | ECM Regulators | Matrisome-associated | 0.3 | 0.15866873 | 0.51623377 | 0.23967502 |
| *loxl2b* | ECM Regulators | Matrisome-associated | 0.12523364 | 0.29721362 | 0.43181818 | 0.1746784 |
| *pcsk5b* | ECM Regulators | Matrisome-associated | 0.38878505 | 0.14705882 | 0.17207792 | 0.16249154 |
| *loxl3b* | ECM Regulators | Matrisome-associated | 0.16728972 | 0.11842105 | 0.33766234 | 0.21936357 |
| *plod2* | ECM Regulators | Matrisome-associated | 0.25607477 | 0.13390093 | 0.14285714 | 0.28842248 |
| *egln1b* | ECM Regulators | Matrisome-associated | 0.3 | 0.20665635 | 0.18506494 | 0.0690589 |
| *adamts17* | ECM Regulators | Matrisome-associated | 0.21869159 | 0.23529412 | 0.09415584 | 0.12322275 |
| *adam10a* | ECM Regulators | Matrisome-associated | 0.11682243 | 0.10990712 | 0.14285714 | 0.28909953 |
| *adamts15a* | ECM Regulators | Matrisome-associated | 0.10934579 | 0.0998452 | 0.16233766 | 0.21530129 |
| *loxa* | ECM Regulators | Matrisome-associated | 0.11214953 | 0.06733746 | 0.1461039 | 0.19092756 |
| *adam9* | ECM Regulators | Matrisome-associated | 0.08785047 | 0.06424149 | 0.08766234 | 0.27623561 |
| *timp2a* | ECM Regulators | Matrisome-associated | 0.11214953 | 0.05727554 | 0.15909091 | 0.11103588 |
| *htra3b* | ECM Regulators | Matrisome-associated | 0.06542056 | 0.05804954 | 0.09415584 | 0.19498984 |
| *adamts3* | ECM Regulators | Matrisome-associated | 0.13084112 | 0.05108359 | 0.0974026 | 0.10968179 |
| *adam12* | ECM Regulators | Matrisome-associated | 0.03271028 | 0.04256966 | 0.1038961 | 0.20920785 |
| *mmp14a* | ECM Regulators | Matrisome-associated | 0.08785047 | 0.0495356 | 0.07792208 | 0.12051456 |
| *sulf2b* | ECM Regulators | Matrisome-associated | 0.08691589 | 0.06501548 | 0.0487013 | 0.11577522 |
| *hpse2* | ECM Regulators | Matrisome-associated | 0.06728972 | 0.05108359 | 0.02597403 | 0.15842925 |
| *adamts10* | ECM Regulators | Matrisome-associated | 0.08691589 | 0.06965944 | 0.05844156 | 0.06770481 |
| *plod1a* | ECM Regulators | Matrisome-associated | 0.09719626 | 0.05263158 | 0.06493506 | 0.05484089 |
| *mmp14b* | ECM Regulators | Matrisome-associated | 0.03457944 | 0.04411765 | 0.06493506 | 0.11035884 |
| *pappaa* | ECM Regulators | Matrisome-associated | 0.07757009 | 0.03328173 | 0.07467532 | 0.06770481 |
| *mmp2* | ECM Regulators | Matrisome-associated | 0.04953271 | 0.02708978 | 0.07467532 | 0.09952607 |
| *ctsla* | ECM Regulators | Matrisome-associated | 0.04392523 | 0.04643963 | 0.05844156 | 0.10155721 |
| *ctsd* | ECM Regulators | Matrisome-associated | 0.03084112 | 0.05959752 | 0.09415584 | 0.06025728 |
| *adamts8a* | ECM Regulators | Matrisome-associated | 0.09065421 | 0.04024768 | 0.0487013 | 0.06161137 |
| *tll1* | ECM Regulators | Matrisome-associated | 0.03551402 | 0.03018576 | 0.0487013 | 0.11848341 |
| *adamts7* | ECM Regulators | Matrisome-associated | 0.0682243 | 0.07275542 | 0.03246753 | 0.05687204 |
| *p4ha3* | ECM Regulators | Matrisome-associated | 0.06448598 | 0.05650155 | 0.05844156 | 0.05077861 |
| *adam15* | ECM Regulators | Matrisome-associated | 0.03925234 | 0.04489164 | 0.05194805 | 0.08666215 |
| *p4ha1b* | ECM Regulators | Matrisome-associated | 0.07757009 | 0.03018576 | 0.05844156 | 0.0534868 |
| *mmp15a* | ECM Regulators | Matrisome-associated | 0.04485981 | 0.07972136 | 0.03896104 | 0.0534868 |
| *serpinh1b* | ECM Regulators | Matrisome-associated | 0.02056075 | 0.02863777 | 0.1038961 | 0.0365606 |
| *serpinf1* | ECM Regulators | Matrisome-associated | 0.05514019 | 0.0255418 | 0.03571429 | 0.07312119 |

| | | | | | | |
|---|---|---|---|---|---|---|
| *fam20b* | ECM Regulators | Matrisome-associated | 0.05233645 | 0.04024768 | 0.03246753 | 0.05619499 |
| *serpine1* | ECM Regulators | Matrisome-associated | 0.03457944 | 0.04798762 | 0.03571429 | 0.04739336 |
| *p4ha2* | ECM Regulators | Matrisome-associated | 0.04579439 | 0.03482972 | 0.03571429 | 0.04536222 |
| *itih5* | ECM Regulators | Matrisome-associated | 0.02429907 | 0.00928793 | 0.01298701 | 0.11035884 |
| *loxl1* | ECM Regulators | Matrisome-associated | 0.02523364 | 0.02941176 | 0.00974026 | 0.08463101 |
| *plat* | ECM Regulators | Matrisome-associated | 0.01962617 | 0.02786378 | 0.03246753 | 0.06770481 |
| *timp2b* | ECM Regulators | Matrisome-associated | 0.03925234 | 0.01393189 | 0.00324675 | 0.0859851 |
| *f13a1b* | ECM Regulators | Matrisome-associated | 0.01214953 | 0.01702786 | 0.02272727 | 0.07853758 |
| *serpine2* | ECM Regulators | Matrisome-associated | 0.03364486 | 0.02012384 | 0.03246753 | 0.04197698 |
| *p4ha1a* | ECM Regulators | Matrisome-associated | 0.02523364 | 0.01315789 | 0.03571429 | 0.0534868 |
| *ctsba* | ECM Regulators | Matrisome-associated | 0.01308411 | 0.0255418 | 0.02922078 | 0.05619499 |
| *adam17a* | ECM Regulators | Matrisome-associated | 0.01775701 | 0.01857585 | 0.0487013 | 0.03791469 |
| *adamts6* | ECM Regulators | Matrisome-associated | 0.03738318 | 0.0255418 | 0.02597403 | 0.03182126 |
| *loxl4* | ECM Regulators | Matrisome-associated | 0.05233645 | 0.01006192 | 0.03896104 | 0.01760325 |
| *pamr1* | ECM Regulators | Matrisome-associated | 0.02803738 | 0.03018576 | 0.01948052 | 0.03994584 |
| *pappab* | ECM Regulators | Matrisome-associated | 0.03271028 | 0.00928793 | 0.02597403 | 0.04468517 |
| *adam22* | ECM Regulators | Matrisome-associated | 0.04018692 | 0.01702786 | 0.02597403 | 0.02505078 |
| *adamts14* | ECM Regulators | Matrisome-associated | 0.01869159 | 0.01083591 | 0.04220779 | 0.02775897 |
| *prss12* | ECM Regulators | Matrisome-associated | 0.01869159 | 0.02863777 | 0.02922078 | 0.0169262 |
| *adam17b* | ECM Regulators | Matrisome-associated | 0.02242991 | 0.01702786 | 0.02597403 | 0.02775897 |
| *tgm2b* | ECM Regulators | Matrisome-associated | 0.01214953 | 0.01702786 | 0.03246753 | 0.02843602 |
| *mmp17a* | ECM Regulators | Matrisome-associated | 0.02149533 | 0.02167183 | 0.01623377 | 0.02640487 |
| *hyal4* | ECM Regulators | Matrisome-associated | 0.0317757 | 0.00541796 | 0.03571429 | 0.00947867 |
| *adamts5* | ECM Regulators | Matrisome-associated | 0.01682243 | 0.01393189 | 0.02272727 | 0.02843602 |
| *ctsf* | ECM Regulators | Matrisome-associated | 0.02149533 | 0.00851393 | 0.02922078 | 0.02166554 |
| *fam20a* | ECM Regulators | Matrisome-associated | 0.02149533 | 0.00541796 | 0.04545455 | 0.00812458 |
| *htra1a* | ECM Regulators | Matrisome-associated | 0.01682243 | 0.00464396 | 0.0487013 | 0.00677048 |
| *ctsc* | ECM Regulators | Matrisome-associated | 0.01588785 | 0.02012384 | 0.00974026 | 0.02979012 |
| *cpamd8* | ECM Regulators | Matrisome-associated | 0.02056075 | 0.01625387 | 0.01948052 | 0.0169262 |
| *bmp1b* | ECM Regulators | Matrisome-associated | 0.04672897 | 0.00541796 | 0.01298701 | 0.00677048 |
| *adamts9* | ECM Regulators | Matrisome-associated | 0.01682243 | 0.01934985 | 0.01948052 | 0.01489506 |
| *serpinh1a* | ECM Regulators | Matrisome-associated | 0.00747664 | 0.01006192 | 0.02272727 | 0.02775897 |
| *mmp23ba* | ECM Regulators | Matrisome-associated | 0.01588785 | 0.02708978 | 0.00649351 | 0.0182803 |
| *ctsk* | ECM Regulators | Matrisome-associated | 0.01308411 | 0.00851393 | 0.00649351 | 0.03588355 |
| *serpina10a* | ECM Regulators | Matrisome-associated | 0.00934579 | 0.00619195 | 0.01298701 | 0.03249831 |
| *pcsk6* | ECM Regulators | Matrisome-associated | 0.01495327 | 0.01006192 | 0.01623377 | 0.01963439 |
| *egln3* | ECM Regulators | Matrisome-associated | 0.02149533 | 0.01702786 | 0.00974026 | 0.01218687 |
| *cst3* | ECM Regulators | Matrisome-associated | 0.01682243 | 0.00619195 | 0.01948052 | 0.01624915 |
| *adam23a* | ECM Regulators | Matrisome-associated | 0.01121495 | 0.01160991 | 0.02272727 | 0.01286391 |
| *plod3* | ECM Regulators | Matrisome-associated | 0.00747664 | 0.00464396 | 0.02597403 | 0.01895735 |
| *kazald3* | ECM Regulators | Matrisome-associated | 0.01308411 | 0.01393189 | 0.02272727 | 0.00677048 |
| *ctso* | ECM Regulators | Matrisome-associated | 0.01962617 | 0.01393189 | 0.00324675 | 0.01760325 |
| *ctsz* | ECM Regulators | Matrisome-associated | 0.01308411 | 0.01625387 | 0 | 0.02505078 |
| *adamts16* | ECM Regulators | Matrisome-associated | 0.0271028 | 0.00773994 | 0.01298701 | 0.00338524 |
| *adam8a* | ECM Regulators | Matrisome-associated | 0.00373832 | 0.00386997 | 0.02272727 | 0.0182803 |

| | | | | | | |
|---|---|---|---|---|---|---|
| *ctsa* | ECM Regulators | Matrisome-associated | 0.00934579 | 0.01393189 | 0.00324675 | 0.01895735 |
| *htra3a* | ECM Regulators | Matrisome-associated | 0.00934579 | 0.00619195 | 0.01623377 | 0.01286391 |
| *hyal1* | ECM Regulators | Matrisome-associated | 0.00841121 | 0 | 0.02272727 | 0.01083277 |
| *loxl3a* | ECM Regulators | Matrisome-associated | 0.01214953 | 0.00386997 | 0.00649351 | 0.01895735 |
| *f13a1a.1* | ECM Regulators | Matrisome-associated | 0.00560748 | 0.01006192 | 0 | 0.02572783 |
| *pappa2* | ECM Regulators | Matrisome-associated | 0.00280374 | 0.00541796 | 0 | 0.03182126 |
| *f10* | ECM Regulators | Matrisome-associated | 0.00373832 | 0.00541796 | 0.02272727 | 0.00812458 |
| *adam28* | ECM Regulators | Matrisome-associated | 0.00654206 | 0.00386997 | 0.01948052 | 0.00744753 |
| *adamts1* | ECM Regulators | Matrisome-associated | 0.00280374 | 0.01160991 | 0.00974026 | 0.01150982 |
| *ctsh* | ECM Regulators | Matrisome-associated | 0.00747664 | 0.00541796 | 0.00974026 | 0.01150982 |
| *hyal2a* | ECM Regulators | Matrisome-associated | 0.00654206 | 0.00154799 | 0.01623377 | 0.00473934 |
| *sulf2a* | ECM Regulators | Matrisome-associated | 0.00373832 | 0.00696594 | 0.00324675 | 0.01286391 |
| *st14a* | ECM Regulators | Matrisome-associated | 0.00280374 | 0.00619195 | 0.00324675 | 0.01218687 |
| *mmp24* | ECM Regulators | Matrisome-associated | 0.00560748 | 0.00232198 | 0.00324675 | 0.01218687 |
| *ngly1* | ECM Regulators | Matrisome-associated | 0.00373832 | 0.00851393 | 0 | 0.01083277 |
| *adam11* | ECM Regulators | Matrisome-associated | 0.00280374 | 0.00386997 | 0.00649351 | 0.00947867 |
| *plaub* | ECM Regulators | Matrisome-associated | 0.00280374 | 0.00309598 | 0.00324675 | 0.01083277 |
| *hyal3* | ECM Regulators | Matrisome-associated | 0.00560748 | 0.00309598 | 0.00324675 | 0.00677048 |
| *tgm2a* | ECM Regulators | Matrisome-associated | 0.00093458 | 0.01006192 | 0.00324675 | 0.00270819 |
| *ctss2.2* | ECM Regulators | Matrisome-associated | 0.00093458 | 0.00309598 | 0.00649351 | 0.00609343 |
| *adamts18* | ECM Regulators | Matrisome-associated | 0.01028037 | 0.00154799 | 0 | 0.00473934 |
| *ogfod1* | ECM Regulators | Matrisome-associated | 0.0046729 | 0.00077399 | 0.00324675 | 0.00744753 |
| *kng1* | ECM Regulators | Matrisome-associated | 0.00186916 | 0.00232198 | 0.00649351 | 0.00473934 |
| *mmp9* | ECM Regulators | Matrisome-associated | 0.0046729 | 0.00232198 | 0.00649351 | 0.0013541 |
| *ctss2.1* | ECM Regulators | Matrisome-associated | 0.00093458 | 0.00232198 | 0.00324675 | 0.00812458 |
| *tgm1* | ECM Regulators | Matrisome-associated | 0.00186916 | 0.00154799 | 0.00974026 | 0.0013541 |
| *spam1* | ECM Regulators | Matrisome-associated | 0.00373832 | 0.00077399 | 0.00649351 | 0.00203114 |
| *st14b* | ECM Regulators | Matrisome-associated | 0 | 0.00309598 | 0.00649351 | 0.00338524 |
| *mmp11a* | ECM Regulators | Matrisome-associated | 0 | 0.00154799 | 0.00324675 | 0.00812458 |
| *f2* | ECM Regulators | Matrisome-associated | 0.00280374 | 0.00232198 | 0.00324675 | 0.00406229 |
| *serpind1* | ECM Regulators | Matrisome-associated | 0.00186916 | 0.00154799 | 0.00649351 | 0.0013541 |
| *adamtsl2* | ECM Regulators | Matrisome-associated | 0.00841121 | 0.00077399 | 0 | 0.00203114 |
| *ctss1* | ECM Regulators | Matrisome-associated | 0.00280374 | 0.00077399 | 0.00649351 | 0.00067705 |
| *cd109* | ECM Regulators | Matrisome-associated | 0.00280374 | 0.00077399 | 0.00649351 | 0.00067705 |
| *serpinb1l2* | ECM Regulators | Matrisome-associated | 0 | 0.00386997 | 0.00649351 | 0 |
| *timp4* | ECM Regulators | Matrisome-associated | 0.00747664 | 0 | 0 | 0.00270819 |
| *egln1a* | ECM Regulators | Matrisome-associated | 0.00186916 | 0.00386997 | 0 | 0.00406229 |
| *p4htm* | ECM Regulators | Matrisome-associated | 0.00186916 | 0.00154799 | 0.00324675 | 0.00270819 |
| *timp4a* | ECM Regulators | Matrisome-associated | 0.00280374 | 0.00309598 | 0 | 0.00338524 |
| *hpse* | ECM Regulators | Matrisome-associated | 0.00280374 | 0.00232198 | 0 | 0.00406229 |
| *hyal6* | ECM Regulators | Matrisome-associated | 0.0046729 | 0.00309598 | 0 | 0.0013541 |
| *tgm5l* | ECM Regulators | Matrisome-associated | 0.00186916 | 0 | 0.00649351 | 0.00067705 |
| *serpini1* | ECM Regulators | Matrisome-associated | 0.0046729 | 0.00077399 | 0 | 0.00338524 |
| *mmp21* | ECM Regulators | Matrisome-associated | 0.00186916 | 0.00154799 | 0.00324675 | 0.00203114 |
| *mmp23bb* | ECM Regulators | Matrisome-associated | 0.00093458 | 0 | 0.00649351 | 0.00067705 |

| | | | | | | |
|---|---|---|---|---|---|---|
| *itih2* | ECM Regulators | Matrisome-associated | 0.00093458 | 0.00077399 | 0.00324675 | 0.00270819 |
| *f13a1a.2* | ECM Regulators | Matrisome-associated | 0.00093458 | 0 | 0.00324675 | 0.00338524 |
| *mmp28* | ECM Regulators | Matrisome-associated | 0.00560748 | 0.00077399 | 0 | 0.00067705 |
| *serpinf2b* | ECM Regulators | Matrisome-associated | 0 | 0.00077399 | 0.00324675 | 0.00270819 |
| *masp1* | ECM Regulators | Matrisome-associated | 0 | 0.00077399 | 0.00324675 | 0.00203114 |
| *mep1b* | ECM Regulators | Matrisome-associated | 0.00093458 | 0.00232198 | 0 | 0.00203114 |
| *plg* | ECM Regulators | Matrisome-associated | 0 | 0 | 0.00324675 | 0.0013541 |
| *serpinc1* | ECM Regulators | Matrisome-associated | 0 | 0 | 0 | 0.00406229 |
| *mmp13a* | ECM Regulators | Matrisome-associated | 0 | 0 | 0.00324675 | 0.00067705 |
| *nots* | ECM Regulators | Matrisome-associated | 0 | 0.00154799 | 0 | 0.00203114 |
| *ogfod2* | ECM Regulators | Matrisome-associated | 0.00280374 | 0 | 0 | 0.00067705 |
| *serpine3* | ECM Regulators | Matrisome-associated | 0.00280374 | 0 | 0 | 0.00067705 |
| *ctsll* | ECM Regulators | Matrisome-associated | 0.00093458 | 0.00077399 | 0 | 0.0013541 |
| *habp2* | ECM Regulators | Matrisome-associated | 0 | 0 | 0 | 0.00270819 |
| *mmp19* | ECM Regulators | Matrisome-associated | 0.00093458 | 0.00154799 | 0 | 0 |
| *mmp16a* | ECM Regulators | Matrisome-associated | 0.00093458 | 0.00077399 | 0 | 0.00067705 |
| *mep1a.1* | ECM Regulators | Matrisome-associated | 0 | 0.00232198 | 0 | 0 |
| *serping1* | ECM Regulators | Matrisome-associated | 0 | 0.00232198 | 0 | 0 |
| *ctsbb* | ECM Regulators | Matrisome-associated | 0 | 0.00077399 | 0 | 0.0013541 |
| *ambp* | ECM Regulators | Matrisome-associated | 0.00093458 | 0.00077399 | 0 | 0 |
| *serpinb1* | ECM Regulators | Matrisome-associated | 0 | 0.00077399 | 0 | 0.00067705 |
| *agt* | ECM Regulators | Matrisome-associated | 0 | 0 | 0 | 0.0013541 |
| *itih3a* | ECM Regulators | Matrisome-associated | 0.00093458 | 0 | 0 | 0 |
| *ctsl.1* | ECM Regulators | Matrisome-associated | 0 | 0.00077399 | 0 | 0 |
| *mep1b* | ECM Regulators | Matrisome-associated | 0 | 0.00077399 | 0 | 0 |
| *plaua* | ECM Regulators | Matrisome-associated | 0 | 0 | 0 | 0.00067705 |
| *tgm1l4* | ECM Regulators | Matrisome-associated | 0 | 0 | 0 | 0.00067705 |

**Table S4**: **ECM-affiliated proteins matrisome genes expressed by fibroblasts** (cluster 5) in each condition and their average expression (log-normalized expression values). Gene classification follows the MatrisomeDB v2.0 annotation framework (Naba et al., 2016).

| Gene_Symbol | Matrisome_Category | Matrisome_Division | Wt-3mt | BM-3mt | Wt-1yr | BM-1yr |
|---|---|---|---|---|---|---|
| *sema4ba* | ECM-affiliated Proteins | Matrisome-associated | 0.65514019 | 0.54489164 | 0.58766234 | 0.96073121 |
| *gpc6a* | ECM-affiliated Proteins | Matrisome-associated | 0.66915888 | 0.64164087 | 0.30194805 | 0.6127285 |
| *cspg4* | ECM-affiliated Proteins | Matrisome-associated | 0.51214953 | 0.3003096 | 0.31493506 | 0.281652 |
| *plxna3* | ECM-affiliated Proteins | Matrisome-associated | 0.27383178 | 0.29102167 | 0.21753247 | 0.51997292 |
| *gpc1b* | ECM-affiliated Proteins | Matrisome-associated | 0.37196262 | 0.34055728 | 0.21103896 | 0.3337847 |
| *plxdc2* | ECM-affiliated Proteins | Matrisome-associated | 0.18411215 | 0.1501548 | 0.16558442 | 0.38794854 |
| *plxna2* | ECM-affiliated Proteins | Matrisome-associated | 0.30934579 | 0.18575851 | 0.21428571 | 0.13134733 |
| *sema3aa* | ECM-affiliated Proteins | Matrisome-associated | 0.23364486 | 0.25464396 | 0.19155844 | 0.10155721 |
| *sdc2* | ECM-affiliated Proteins | Matrisome-associated | 0.19158879 | 0.1501548 | 0.17857143 | 0.20040623 |
| *plxna4* | ECM-affiliated Proteins | Matrisome-associated | 0.20654206 | 0.14396285 | 0.11363636 | 0.19498984 |
| *sema3c* | ECM-affiliated Proteins | Matrisome-associated | 0.19252336 | 0.07894737 | 0.10064935 | 0.19566689 |
| *sema5a* | ECM-affiliated Proteins | Matrisome-associated | 0.18971963 | 0.14551084 | 0.16883117 | 0.05145565 |
| *sema4c* | ECM-affiliated Proteins | Matrisome-associated | 0.11775701 | 0.0874613 | 0.14285714 | 0.09884902 |
| *gpc5b* | ECM-affiliated Proteins | Matrisome-associated | 0.11495327 | 0.17647059 | 0.0487013 | 0.02234259 |
| *anxa2a* | ECM-affiliated Proteins | Matrisome-associated | 0.03084112 | 0.11764706 | 0.05844156 | 0.14827353 |
| *frem3* | ECM-affiliated Proteins | Matrisome-associated | 0.04579439 | 0.17182663 | 0.07792208 | 0.04129993 |
| *sema3fb* | ECM-affiliated Proteins | Matrisome-associated | 0.05700935 | 0.0874613 | 0.06168831 | 0.10968179 |
| *colec12* | ECM-affiliated Proteins | Matrisome-associated | 0.04579439 | 0.06114551 | 0.06168831 | 0.13608666 |
| *anxa11a* | ECM-affiliated Proteins | Matrisome-associated | 0.05233645 | 0.04102167 | 0.07142857 | 0.11780636 |
| *sema3d* | ECM-affiliated Proteins | Matrisome-associated | 0.07196262 | 0.05185759 | 0.09415584 | 0.05619499 |
| *sema3ab* | ECM-affiliated Proteins | Matrisome-associated | 0.08130841 | 0.08204334 | 0.05194805 | 0.05551794 |
| *sema3fa* | ECM-affiliated Proteins | Matrisome-associated | 0.05420561 | 0.08591331 | 0.05844156 | 0.06770481 |
| *plxna1a* | ECM-affiliated Proteins | Matrisome-associated | 0.08224299 | 0.04102167 | 0.05519481 | 0.08192282 |
| *sema3b* | ECM-affiliated Proteins | Matrisome-associated | 0.02429907 | 0.06733746 | 0.05194805 | 0.11577522 |
| *gpc4* | ECM-affiliated Proteins | Matrisome-associated | 0.0588785 | 0.02941176 | 0.09090909 | 0.07582938 |
| *anxa6* | ECM-affiliated Proteins | Matrisome-associated | 0.13925234 | 0.00386997 | 0.07142857 | 0.00203114 |
| *sema4f* | ECM-affiliated Proteins | Matrisome-associated | 0.12616822 | 0.02786378 | 0.02922078 | 0.02437373 |
| *sdc4* | ECM-affiliated Proteins | Matrisome-associated | 0.03084112 | 0.02321981 | 0.03246753 | 0.07853758 |
| *sema5ba* | ECM-affiliated Proteins | Matrisome-associated | 0.02523364 | 0.01857585 | 0.00649351 | 0.10494245 |
| *clec19a* | ECM-affiliated Proteins | Matrisome-associated | 0.01869159 | 0.04024768 | 0.04545455 | 0.04603927 |
| *sema3gb* | ECM-affiliated Proteins | Matrisome-associated | 0.04392523 | 0.02244582 | 0.05844156 | 0.02301963 |
| *sema6dl* | ECM-affiliated Proteins | Matrisome-associated | 0.02897196 | 0.01315789 | 0.05844156 | 0.0352065 |
| *anxa4* | ECM-affiliated Proteins | Matrisome-associated | 0.02149533 | 0.02012384 | 0.02922078 | 0.04739336 |
| *gpc5a* | ECM-affiliated Proteins | Matrisome-associated | 0.02803738 | 0.02708978 | 0.01623377 | 0.04536222 |
| *sema4bb* | ECM-affiliated Proteins | Matrisome-associated | 0.0317757 | 0.01547988 | 0.02597403 | 0.04197698 |
| *anxa1a* | ECM-affiliated Proteins | Matrisome-associated | 0.02149533 | 0.02012384 | 0.02597403 | 0.04400812 |
| *anxa5b* | ECM-affiliated Proteins | Matrisome-associated | 0.02056075 | 0.01393189 | 0.02922078 | 0.04468517 |
| *plxnb2b* | ECM-affiliated Proteins | Matrisome-associated | 0.01214953 | 0.00696594 | 0.06818182 | 0.01760325 |
| *frem2b* | ECM-affiliated Proteins | Matrisome-associated | 0.01962617 | 0.01780186 | 0.02272727 | 0.03791469 |
| *lgals3a* | ECM-affiliated Proteins | Matrisome-associated | 0.02242991 | 0.01547988 | 0.02272727 | 0.03588355 |

| | | | | | | |
|---|---|---|---|---|---|---|
| *plxnc1* | ECM-affiliated Proteins | Matrisome-associated | 0.02242991 | 0.02089783 | 0.00649351 | 0.03723764 |
| *sema3e* | ECM-affiliated Proteins | Matrisome-associated | 0.01401869 | 0.00464396 | 0.04545455 | 0.0182803 |
| *anxa11b* | ECM-affiliated Proteins | Matrisome-associated | 0.02429907 | 0.00773994 | 0.02272727 | 0.02098849 |
| *anxa3b* | ECM-affiliated Proteins | Matrisome-associated | 0.01401869 | 0.01547988 | 0.00649351 | 0.03926879 |
| *c1qtnf5* | ECM-affiliated Proteins | Matrisome-associated | 0.01588785 | 0.00077399 | 0.01298701 | 0.04062288 |
| *c1qtnf1* | ECM-affiliated Proteins | Matrisome-associated | 0.00560748 | 0.00773994 | 0.03896104 | 0.01760325 |
| *anxa13l* | ECM-affiliated Proteins | Matrisome-associated | 0.01214953 | 0.00773994 | 0.03571429 | 0.01083277 |
| *c1qtnf7* | ECM-affiliated Proteins | Matrisome-associated | 0.01401869 | 0.01702786 | 0 | 0.0352065 |
| *sema3ga* | ECM-affiliated Proteins | Matrisome-associated | 0.00747664 | 0.01006192 | 0.01948052 | 0.02708192 |
| *c1qtnf9* | ECM-affiliated Proteins | Matrisome-associated | 0.01121495 | 0.01780186 | 0.00649351 | 0.02843602 |
| *sema4ga* | ECM-affiliated Proteins | Matrisome-associated | 0.0046729 | 0.00773994 | 0.01948052 | 0.0182803 |
| *lman1* | ECM-affiliated Proteins | Matrisome-associated | 0.01028037 | 0.00851393 | 0.01298701 | 0.01760325 |
| *lgals3b* | ECM-affiliated Proteins | Matrisome-associated | 0.01028037 | 0.00773994 | 0.00649351 | 0.02437373 |
| *anxa13* | ECM-affiliated Proteins | Matrisome-associated | 0.0046729 | 0.00619195 | 0.00974026 | 0.02505078 |
| *lgals2a* | ECM-affiliated Proteins | Matrisome-associated | 0.01308411 | 0.00773994 | 0 | 0.02301963 |
| *gpc1a* | ECM-affiliated Proteins | Matrisome-associated | 0.01308411 | 0.00928793 | 0.01298701 | 0.00812458 |
| *lgalsla* | ECM-affiliated Proteins | Matrisome-associated | 0.00654206 | 0.01625387 | 0 | 0.01895735 |
| *cspg5b* | ECM-affiliated Proteins | Matrisome-associated | 0.01495327 | 0.01083591 | 0 | 0.01489506 |
| *c1qtnf6a* | ECM-affiliated Proteins | Matrisome-associated | 0.00186916 | 0.00309598 | 0.02597403 | 0.00338524 |
| *lgals8a* | ECM-affiliated Proteins | Matrisome-associated | 0.00841121 | 0.00232198 | 0.00974026 | 0.01083277 |
| *c1qb* | ECM-affiliated Proteins | Matrisome-associated | 0.00373832 | 0.00232198 | 0.00974026 | 0.01286391 |
| *lgals9l3* | ECM-affiliated Proteins | Matrisome-associated | 0.00373832 | 0.00464396 | 0.00324675 | 0.01557211 |
| *frem1b* | ECM-affiliated Proteins | Matrisome-associated | 0.00373832 | 0.00773994 | 0 | 0.01421801 |
| *sema3bl* | ECM-affiliated Proteins | Matrisome-associated | 0.00373832 | 0.00541796 | 0.00324675 | 0.01218687 |
| *gpc6b* | ECM-affiliated Proteins | Matrisome-associated | 0.00841121 | 0.00232198 | 0.00649351 | 0.00541638 |
| *sema7a* | ECM-affiliated Proteins | Matrisome-associated | 0.00560748 | 0.00541796 | 0 | 0.01150982 |
| *sema6d* | ECM-affiliated Proteins | Matrisome-associated | 0.00186916 | 0.00386997 | 0 | 0.01624915 |
| *frem1a* | ECM-affiliated Proteins | Matrisome-associated | 0.00186916 | 0.00309598 | 0.01298701 | 0.00338524 |
| *clec3a* | ECM-affiliated Proteins | Matrisome-associated | 0.00747664 | 0.00309598 | 0.00649351 | 0.00406229 |
| *gpc3* | ECM-affiliated Proteins | Matrisome-associated | 0.00560748 | 0.00464396 | 0.00324675 | 0.00677048 |
| *elfn1a* | ECM-affiliated Proteins | Matrisome-associated | 0.00280374 | 0.00464396 | 0.00324675 | 0.00947867 |
| *c1qtnf2* | ECM-affiliated Proteins | Matrisome-associated | 0.00280374 | 0.00154799 | 0.00324675 | 0.01218687 |
| *sema6a* | ECM-affiliated Proteins | Matrisome-associated | 0.00280374 | 0.00232198 | 0.00324675 | 0.00947867 |
| *lgals9l1* | ECM-affiliated Proteins | Matrisome-associated | 0 | 0.00154799 | 0.00649351 | 0.00744753 |
| *frem2a* | ECM-affiliated Proteins | Matrisome-associated | 0.00560748 | 0.00309598 | 0.00324675 | 0.00203114 |
| *c1qc* | ECM-affiliated Proteins | Matrisome-associated | 0.00186916 | 0.00541796 | 0.00324675 | 0.00203114 |
| *colec10* | ECM-affiliated Proteins | Matrisome-associated | 0 | 0.00309598 | 0.00324675 | 0.00338524 |
| *plxdc1* | ECM-affiliated Proteins | Matrisome-associated | 0.00186916 | 0.00541796 | 0 | 0.00203114 |
| *sftpbb* | ECM-affiliated Proteins | Matrisome-associated | 0.00186916 | 0.00309598 | 0 | 0.00406229 |
| *anxa1b* | ECM-affiliated Proteins | Matrisome-associated | 0 | 0 | 0.00649351 | 0.0013541 |
| *clec14a* | ECM-affiliated Proteins | Matrisome-associated | 0.00093458 | 0.00077399 | 0.00324675 | 0.0013541 |
| *c1qa* | ECM-affiliated Proteins | Matrisome-associated | 0.00186916 | 0.00154799 | 0 | 0.00270819 |
| *itln1* | ECM-affiliated Proteins | Matrisome-associated | 0 | 0.00154799 | 0 | 0.00338524 |
| *c1qtnf4* | ECM-affiliated Proteins | Matrisome-associated | 0.00186916 | 0.00232198 | 0 | 0.00067705 |
| *elfn2a* | ECM-affiliated Proteins | Matrisome-associated | 0.00093458 | 0 | 0.00324675 | 0 |

| | | | | | | |
|---|---|---|---|---|---|---|
| *anxa5a* | ECM-affiliated Proteins | Matrisome-associated | 0 | 0.00077399 | 0.00324675 | 0 |
| *colec11* | ECM-affiliated Proteins | Matrisome-associated | 0.00093458 | 0.00154799 | 0 | 0.0013541 |
| *lgalslb* | ECM-affiliated Proteins | Matrisome-associated | 0.00093458 | 0.00077399 | 0 | 0.0013541 |
| *clec3ba* | ECM-affiliated Proteins | Matrisome-associated | 0.00093458 | 0 | 0 | 0.00203114 |
| *anxa2b* | ECM-affiliated Proteins | Matrisome-associated | 0 | 0.00077399 | 0 | 0.0013541 |
| *lgals1l1* | ECM-affiliated Proteins | Matrisome-associated | 0.00186916 | 0 | 0 | 0 |
| *hbl3* | ECM-affiliated Proteins | Matrisome-associated | 0.00093458 | 0.00077399 | 0 | 0 |
| *lgals9l4* | ECM-affiliated Proteins | Matrisome-associated | 0.00093458 | 0 | 0 | 0.00067705 |
| *anxa3a* | ECM-affiliated Proteins | Matrisome-associated | 0 | 0.00077399 | 0 | 0.00067705 |
| *cd209* | ECM-affiliated Proteins | Matrisome-associated | 0 | 0.00077399 | 0 | 0.00067705 |
| *c1ql3b* | ECM-affiliated Proteins | Matrisome-associated | 0 | 0 | 0 | 0.0013541 |
| *c1ql4a* | ECM-affiliated Proteins | Matrisome-associated | 0 | 0.00077399 | 0 | 0 |
| *anxa1c* | ECM-affiliated Proteins | Matrisome-associated | 0 | 0 | 0 | 0.00067705 |
| *c1ql3a* | ECM-affiliated Proteins | Matrisome-associated | 0 | 0 | 0 | 0.00067705 |

**Table S5**: **ECM-secreted factor matrisome genes expressed by fibroblasts** (cluster 5) in each condition and their average expression (log-normalized expression values). Gene classification follows the MatrisomeDB v2.0 annotation framework (Naba et al., 2016).

| Gene_Symbol | Matrisome_Category | Matrisome_Division | Wt-3mt | BM-3mt | Wt-1yr | BM-1yr |
|---|---|---|---|---|---|---|
| *tgfb2* | Secreted Factors | Matrisome-associated | 0.33271028 | 0.37925697 | 0.31168831 | 0.44685173 |
| *fstl1b* | Secreted Factors | Matrisome-associated | 0.27757009 | 0.25464396 | 0.14935065 | 0.4915369 |
| *angptl1a* | Secreted Factors | Matrisome-associated | 0.20186916 | 0.20123839 | 0.15909091 | 0.36492891 |
| *vegfaa* | Secreted Factors | Matrisome-associated | 0.20186916 | 0.20123839 | 0.27272727 | 0.23019634 |
| *pik3ip1* | Secreted Factors | Matrisome-associated | 0.21121495 | 0.07585139 | 0.17207792 | 0.11983751 |
| *nrg2a* | Secreted Factors | Matrisome-associated | 0.10934579 | 0.12925697 | 0.10714286 | 0.17535545 |
| *nrg1* | Secreted Factors | Matrisome-associated | 0.12523364 | 0.08591331 | 0.1038961 | 0.17197021 |
| *fgf12b* | Secreted Factors | Matrisome-associated | 0.14859813 | 0.04256966 | 0.16233766 | 0.05551794 |
| *tnfsf10l* | Secreted Factors | Matrisome-associated | 0.08411215 | 0.08049536 | 0.07142857 | 0.17264726 |
| *angpt1* | Secreted Factors | Matrisome-associated | 0.10934579 | 0.04179567 | 0.08766234 | 0.15504401 |
| *il15* | Secreted Factors | Matrisome-associated | 0.07476636 | 0.05882353 | 0.05844156 | 0.18483412 |
| *pdgfc* | Secreted Factors | Matrisome-associated | 0.08504673 | 0.05650155 | 0.17857143 | 0.05619499 |
| *vegfab* | Secreted Factors | Matrisome-associated | 0.1 | 0.08823529 | 0.08441558 | 0.08327691 |
| *wnt5b* | Secreted Factors | Matrisome-associated | 0.07102804 | 0.04566563 | 0.13311688 | 0.09952607 |
| *cxcl12a* | Secreted Factors | Matrisome-associated | 0.02803738 | 0.06811146 | 0.11688312 | 0.12322275 |
| *angptl4* | Secreted Factors | Matrisome-associated | 0.20841121 | 0.03095975 | 0.02597403 | 0.06973595 |
| *fgf13a* | Secreted Factors | Matrisome-associated | 0.06635514 | 0.05108359 | 0.11038961 | 0.07853758 |
| *ccbe1* | Secreted Factors | Matrisome-associated | 0.05700935 | 0.0379257 | 0.03246753 | 0.16790792 |
| *wnt9a* | Secreted Factors | Matrisome-associated | 0.05514019 | 0.05263158 | 0.06818182 | 0.11103588 |
| *eda* | Secreted Factors | Matrisome-associated | 0.02897196 | 0.0619195 | 0.07467532 | 0.11848341 |
| *scube1* | Secreted Factors | Matrisome-associated | 0.02523364 | 0.04876161 | 0.0974026 | 0.06161137 |
| *il16* | Secreted Factors | Matrisome-associated | 0.05607477 | 0.08359133 | 0.02597403 | 0.05619499 |
| *ngfb* | Secreted Factors | Matrisome-associated | 0.03925234 | 0.04334365 | 0.01948052 | 0.11238998 |
| *hcfc1b* | Secreted Factors | Matrisome-associated | 0.04485981 | 0.04876161 | 0.03246753 | 0.08056872 |
| *fgf18a* | Secreted Factors | Matrisome-associated | 0.03084112 | 0.0379257 | 0.02272727 | 0.09681787 |
| *megf8* | Secreted Factors | Matrisome-associated | 0.03271028 | 0.03018576 | 0.05844156 | 0.06567366 |
| *tgfb3* | Secreted Factors | Matrisome-associated | 0.05140187 | 0.02786378 | 0.02922078 | 0.05958023 |
| *megf10* | Secreted Factors | Matrisome-associated | 0.03551402 | 0.02631579 | 0.03896104 | 0.04874746 |
| *pdgfd* | Secreted Factors | Matrisome-associated | 0.0271028 | 0.02631579 | 0.03571429 | 0.05280975 |
| *s100a10b* | Secreted Factors | Matrisome-associated | 0.02242991 | 0.02089783 | 0.04220779 | 0.05551794 |
| *egfl6* | Secreted Factors | Matrisome-associated | 0.03738318 | 0.04643963 | 0.02922078 | 0.02369668 |
| *ntf3* | Secreted Factors | Matrisome-associated | 0.0411215 | 0.02012384 | 0.00974026 | 0.06364252 |
| *pdgfab* | Secreted Factors | Matrisome-associated | 0.03738318 | 0.03095975 | 0.02597403 | 0.03994584 |
| *hgfb* | Secreted Factors | Matrisome-associated | 0.01588785 | 0.01547988 | 0.01623377 | 0.0859851 |
| *csf1a* | Secreted Factors | Matrisome-associated | 0.02803738 | 0.01315789 | 0.02272727 | 0.06838186 |
| *angptl2a* | Secreted Factors | Matrisome-associated | 0.0317757 | 0.03947368 | 0.00974026 | 0.04603927 |
| *bmp7b* | Secreted Factors | Matrisome-associated | 0.01775701 | 0.02012384 | 0.01298701 | 0.07244414 |
| *crlf1a* | Secreted Factors | Matrisome-associated | 0.02242991 | 0.01547988 | 0.02922078 | 0.05484089 |
| *fam132a* | Secreted Factors | Matrisome-associated | 0.03551402 | 0.01625387 | 0.05519481 | 0.01286391 |
| *vegfc* | Secreted Factors | Matrisome-associated | 0.03364486 | 0.01470588 | 0.01948052 | 0.04739336 |

| | | | | | | |
|---|---|---|---|---|---|---|
| *bmp6* | Secreted Factors | Matrisome-associated | 0.02056075 | 0.01547988 | 0.05194805 | 0.01557211 |
| *crlf3* | Secreted Factors | Matrisome-associated | 0.02336449 | 0.01006192 | 0.01623377 | 0.0534868 |
| *hbegfa* | Secreted Factors | Matrisome-associated | 0.02242991 | 0.01702786 | 0.02597403 | 0.0365606 |
| *angptl7* | Secreted Factors | Matrisome-associated | 0.02990654 | 0.0247678 | 0.01948052 | 0.02234259 |
| *fstl1a* | Secreted Factors | Matrisome-associated | 0.01028037 | 0.01470588 | 0.00974026 | 0.06161137 |
| *tgfb1a* | Secreted Factors | Matrisome-associated | 0.01214953 | 0.01470588 | 0.02597403 | 0.04265403 |
| *igf2b* | Secreted Factors | Matrisome-associated | 0.01401869 | 0.01547988 | 0.01623377 | 0.04468517 |
| *angpt4* | Secreted Factors | Matrisome-associated | 0.01495327 | 0.0255418 | 0.00649351 | 0.04265403 |
| *scube3* | Secreted Factors | Matrisome-associated | 0.01214953 | 0.01857585 | 0.02272727 | 0.03114421 |
| *megf6a* | Secreted Factors | Matrisome-associated | 0.02803738 | 0.02399381 | 0 | 0.02843602 |
| *bmp4* | Secreted Factors | Matrisome-associated | 0.01308411 | 0.00928793 | 0.01623377 | 0.0352065 |
| *fgf12a* | Secreted Factors | Matrisome-associated | 0.00373832 | 0.0495356 | 0.00974026 | 0.00609343 |
| *angptl2b* | Secreted Factors | Matrisome-associated | 0.02616822 | 0.01083591 | 0.01623377 | 0.01557211 |
| *inhbaa* | Secreted Factors | Matrisome-associated | 0.01121495 | 0.01934985 | 0.00974026 | 0.02505078 |
| *fgf11a* | Secreted Factors | Matrisome-associated | 0.01121495 | 0.01547988 | 0.00649351 | 0.03182126 |
| *hcfc1a* | Secreted Factors | Matrisome-associated | 0.01214953 | 0.01934985 | 0.01298701 | 0.01557211 |
| *gdnfa* | Secreted Factors | Matrisome-associated | 0.02242991 | 0.00232198 | 0.01298701 | 0.02098849 |
| *il34* | Secreted Factors | Matrisome-associated | 0.00373832 | 0.00541796 | 0.00649351 | 0.04265403 |
| *fgf6a* | Secreted Factors | Matrisome-associated | 0.01869159 | 0.01934985 | 0.00649351 | 0.01015572 |
| *kitlga* | Secreted Factors | Matrisome-associated | 0.00747664 | 0.00464396 | 0.01948052 | 0.02301963 |
| *csf1b* | Secreted Factors | Matrisome-associated | 0.00186916 | 0.0123839 | 0.02597403 | 0.01421801 |
| *nrg2b* | Secreted Factors | Matrisome-associated | 0.01588785 | 0.01702786 | 0.00974026 | 0.01083277 |
| *ccl25b* | Secreted Factors | Matrisome-associated | 0.00280374 | 0.01625387 | 0.01298701 | 0.01895735 |
| *figf* | Secreted Factors | Matrisome-associated | 0.00560748 | 0.00773994 | 0 | 0.03588355 |
| *igf1* | Secreted Factors | Matrisome-associated | 0.01495327 | 0.01083591 | 0.00324675 | 0.01963439 |
| *ptn* | Secreted Factors | Matrisome-associated | 0.00280374 | 0.03328173 | 0.00649351 | 0.00541638 |
| *angptl5* | Secreted Factors | Matrisome-associated | 0.0046729 | 0.01315789 | 0.01298701 | 0.0169262 |
| *ism1* | Secreted Factors | Matrisome-associated | 0.00280374 | 0.00386997 | 0.02272727 | 0.01760325 |
| *brinp2* | Secreted Factors | Matrisome-associated | 0.00654206 | 0.01393189 | 0.01298701 | 0.01218687 |
| *cxcl12b* | Secreted Factors | Matrisome-associated | 0.01588785 | 0.00696594 | 0.00324675 | 0.01624915 |
| *fstb* | Secreted Factors | Matrisome-associated | 0.00373832 | 0.00619195 | 0.01948052 | 0.01286391 |
| *mdka* | Secreted Factors | Matrisome-associated | 0.01028037 | 0.01393189 | 0.00324675 | 0.01421801 |
| *frzb* | Secreted Factors | Matrisome-associated | 0.01214953 | 0.00464396 | 0.00974026 | 0.01489506 |
| *clcf1* | Secreted Factors | Matrisome-associated | 0.00841121 | 0.00773994 | 0.01298701 | 0.01218687 |
| *egfl7* | Secreted Factors | Matrisome-associated | 0.01121495 | 0.01083591 | 0.00649351 | 0.01218687 |
| *wnt7ba* | Secreted Factors | Matrisome-associated | 0.00186916 | 0.00541796 | 0.01298701 | 0.02031144 |
| *ccl19a.1* | Secreted Factors | Matrisome-associated | 0.00841121 | 0.00773994 | 0.00649351 | 0.01760325 |
| *ngfa* | Secreted Factors | Matrisome-associated | 0.00560748 | 0.00619195 | 0.01298701 | 0.01421801 |
| *igf2a* | Secreted Factors | Matrisome-associated | 0.01028037 | 0.00541796 | 0.00649351 | 0.01624915 |
| *hhip* | Secreted Factors | Matrisome-associated | 0.01121495 | 0.00309598 | 0.00649351 | 0.0169262 |
| *sfrp1b* | Secreted Factors | Matrisome-associated | 0.00186916 | 0.00309598 | 0.01298701 | 0.01895735 |
| *sfrp1a* | Secreted Factors | Matrisome-associated | 0.00560748 | 0.00928793 | 0.00324675 | 0.01760325 |
| *btc* | Secreted Factors | Matrisome-associated | 0.01308411 | 0.01315789 | 0 | 0.00880162 |
| *angpt2b* | Secreted Factors | Matrisome-associated | 0.01028037 | 0.00851393 | 0.01298701 | 0.00203114 |
| *brinp3a-1* | Secreted Factors | Matrisome-associated | 0.00560748 | 0.00851393 | 0.00324675 | 0.01489506 |

| | | | | | | |
|---|---|---|---|---|---|---|
| *sfrp5* | Secreted Factors | Matrisome-associated | 0.00280374 | 0.00464396 | 0.00324675 | 0.02031144 |
| *crhbp* | Secreted Factors | Matrisome-associated | 0.0046729 | 0.00541796 | 0.00324675 | 0.01760325 |
| *fsta* | Secreted Factors | Matrisome-associated | 0.00934579 | 0.00386997 | 0.00324675 | 0.01421801 |
| *gdf11* | Secreted Factors | Matrisome-associated | 0.00186916 | 0.00773994 | 0.00649351 | 0.01421801 |
| *fgf7* | Secreted Factors | Matrisome-associated | 0.00186916 | 0 | 0 | 0.02640487 |
| *fstl3* | Secreted Factors | Matrisome-associated | 0.00093458 | 0.00464396 | 0.01298701 | 0.00880162 |
| *cntf* | Secreted Factors | Matrisome-associated | 0.0046729 | 0.00309598 | 0.00649351 | 0.01286391 |
| *chd* | Secreted Factors | Matrisome-associated | 0.00280374 | 0.00309598 | 0.00974026 | 0.00677048 |
| *bmp5* | Secreted Factors | Matrisome-associated | 0.01214953 | 0.00154799 | 0.00324675 | 0.00406229 |
| *s100b* | Secreted Factors | Matrisome-associated | 0.00373832 | 0.00232198 | 0.00974026 | 0.00406229 |
| *wnt16* | Secreted Factors | Matrisome-associated | 0.00560748 | 0.00464396 | 0 | 0.00947867 |
| *hbegfb* | Secreted Factors | Matrisome-associated | 0.00654206 | 0.00619195 | 0.00324675 | 0.00338524 |
| *mstnb* | Secreted Factors | Matrisome-associated | 0.00373832 | 0.00541796 | 0.00649351 | 0.00270819 |
| *tnfsf12* | Secreted Factors | Matrisome-associated | 0.00093458 | 0.00154799 | 0.00974026 | 0.00609343 |
| *hgfa* | Secreted Factors | Matrisome-associated | 0.0046729 | 0.00619195 | 0.00324675 | 0.00406229 |
| *fgf10a* | Secreted Factors | Matrisome-associated | 0.0046729 | 0.00309598 | 0 | 0.00812458 |
| *fgf16* | Secreted Factors | Matrisome-associated | 0.00654206 | 0.00309598 | 0 | 0.00609343 |
| *crlf1b* | Secreted Factors | Matrisome-associated | 0.00280374 | 0.00077399 | 0.00974026 | 0.00203114 |
| *csf3a* | Secreted Factors | Matrisome-associated | 0.0046729 | 0.00928793 | 0 | 0.0013541 |
| *fgf14* | Secreted Factors | Matrisome-associated | 0.00186916 | 0.00464396 | 0.00324675 | 0.00406229 |
| *egf* | Secreted Factors | Matrisome-associated | 0 | 0.00309598 | 0.00649351 | 0.00338524 |
| *wnt11* | Secreted Factors | Matrisome-associated | 0.00280374 | 0.00309598 | 0 | 0.00609343 |
| *ihhb* | Secreted Factors | Matrisome-associated | 0.00280374 | 0.00309598 | 0.00324675 | 0.00270819 |
| *dhh* | Secreted Factors | Matrisome-associated | 0.00654206 | 0.00232198 | 0 | 0.00270819 |
| *fgf4* | Secreted Factors | Matrisome-associated | 0.00280374 | 0.00386997 | 0 | 0.00473934 |
| *wnt6a* | Secreted Factors | Matrisome-associated | 0.00093458 | 0.00232198 | 0.00324675 | 0.00473934 |
| *bmp2b* | Secreted Factors | Matrisome-associated | 0.00373832 | 0.00154799 | 0.00324675 | 0.00270819 |
| *mst1* | Secreted Factors | Matrisome-associated | 0 | 0 | 0.00974026 | 0.0013541 |
| *angptl3* | Secreted Factors | Matrisome-associated | 0.00654206 | 0.00309598 | 0 | 0.0013541 |
| *cxcl14* | Secreted Factors | Matrisome-associated | 0.00280374 | 0 | 0 | 0.00812458 |
| *wif1* | Secreted Factors | Matrisome-associated | 0.00560748 | 0 | 0.00324675 | 0.00203114 |
| *tnfsf10* | Secreted Factors | Matrisome-associated | 0.00186916 | 0.00541796 | 0 | 0.00338524 |
| *tnfsf13b* | Secreted Factors | Matrisome-associated | 0.00373832 | 0 | 0.00324675 | 0.00338524 |
| *scube2* | Secreted Factors | Matrisome-associated | 0.00186916 | 0 | 0.00324675 | 0.00473934 |
| *il17d* | Secreted Factors | Matrisome-associated | 0.00093458 | 0.00464396 | 0 | 0.00406229 |
| *fgf1b* | Secreted Factors | Matrisome-associated | 0.00280374 | 0.00464396 | 0 | 0.00203114 |
| *wnt10a* | Secreted Factors | Matrisome-associated | 0.00186916 | 0.00077399 | 0 | 0.00677048 |
| *s100a11* | Secreted Factors | Matrisome-associated | 0.00093458 | 0.00232198 | 0.00324675 | 0.00270819 |
| *il1b* | Secreted Factors | Matrisome-associated | 0 | 0.00309598 | 0.00324675 | 0.00270819 |
| *wnt2bb* | Secreted Factors | Matrisome-associated | 0 | 0.00077399 | 0 | 0.00812458 |
| *ihha* | Secreted Factors | Matrisome-associated | 0.00280374 | 0 | 0.00324675 | 0.00270819 |
| *fgf2* | Secreted Factors | Matrisome-associated | 0.00093458 | 0.00154799 | 0 | 0.00609343 |
| *inha* | Secreted Factors | Matrisome-associated | 0 | 0.00696594 | 0 | 0.00067705 |
| *brinp3a.2* | Secreted Factors | Matrisome-associated | 0 | 0.00154799 | 0.00324675 | 0.00270819 |
| *lepb* | Secreted Factors | Matrisome-associated | 0 | 0 | 0 | 0.00744753 |

| | | | | | | |
|---|---|---|---|---|---|---|
| *vwc2l* | Secreted Factors | Matrisome-associated | 0.00373832 | 0.00232198 | 0 | 0.0013541 |
| *bmp3* | Secreted Factors | Matrisome-associated | 0.00186916 | 0.00154799 | 0.00324675 | 0.00067705 |
| *fgf18b* | Secreted Factors | Matrisome-associated | 0.00093458 | 0.00154799 | 0.00324675 | 0.0013541 |
| *s100a10a* | Secreted Factors | Matrisome-associated | 0.00186916 | 0.00309598 | 0 | 0.00203114 |
| *mdkb* | Secreted Factors | Matrisome-associated | 0.00280374 | 0.00077399 | 0 | 0.00338524 |
| *ism2b* | Secreted Factors | Matrisome-associated | 0 | 0.00154799 | 0.00324675 | 0.00203114 |
| *tgfa* | Secreted Factors | Matrisome-associated | 0.00093458 | 0.00154799 | 0 | 0.00406229 |
| *cbln4* | Secreted Factors | Matrisome-associated | 0.00186916 | 0 | 0.00324675 | 0.0013541 |
| *s100z* | Secreted Factors | Matrisome-associated | 0.00280374 | 0.00154799 | 0 | 0.00203114 |
| *wnt2ba* | Secreted Factors | Matrisome-associated | 0 | 0.00309598 | 0.00324675 | 0 |
| *ccl27b* | Secreted Factors | Matrisome-associated | 0.00186916 | 0.00232198 | 0 | 0.00203114 |
| *il19l* | Secreted Factors | Matrisome-associated | 0.00186916 | 0 | 0 | 0.00406229 |
| *cxcl8b.1* | Secreted Factors | Matrisome-associated | 0 | 0.00154799 | 0 | 0.00406229 |
| *epoa* | Secreted Factors | Matrisome-associated | 0.00280374 | 0.00077399 | 0 | 0.00203114 |
| *fgf10b* | Secreted Factors | Matrisome-associated | 0.00280374 | 0.00077399 | 0 | 0.00203114 |
| *wnt4b* | Secreted Factors | Matrisome-associated | 0 | 0.00464396 | 0 | 0.00067705 |
| *chrdl2* | Secreted Factors | Matrisome-associated | 0 | 0.00232198 | 0 | 0.00270819 |
| *fgf5* | Secreted Factors | Matrisome-associated | 0 | 0.00232198 | 0 | 0.00270819 |
| *fgf6b* | Secreted Factors | Matrisome-associated | 0.00186916 | 0.00232198 | 0 | 0.00067705 |
| *ccl19b* | Secreted Factors | Matrisome-associated | 0.00093458 | 0 | 0.00324675 | 0.00067705 |
| *fgf1a* | Secreted Factors | Matrisome-associated | 0 | 0.00077399 | 0.00324675 | 0.00067705 |
| *cbln1* | Secreted Factors | Matrisome-associated | 0.00093458 | 0.00232198 | 0 | 0.0013541 |
| *pdgfaa* | Secreted Factors | Matrisome-associated | 0.00186916 | 0 | 0 | 0.00270819 |
| *inhbb* | Secreted Factors | Matrisome-associated | 0.00093458 | 0.00154799 | 0 | 0.00203114 |
| *wnt7aa* | Secreted Factors | Matrisome-associated | 0.00093458 | 0.00077399 | 0 | 0.00270819 |
| *gdf5* | Secreted Factors | Matrisome-associated | 0.00093458 | 0.00077399 | 0 | 0.00270819 |
| *kitlgb* | Secreted Factors | Matrisome-associated | 0.00093458 | 0 | 0.00324675 | 0 |
| *fgf17* | Secreted Factors | Matrisome-associated | 0 | 0.00077399 | 0 | 0.00338524 |
| *bmp8a* | Secreted Factors | Matrisome-associated | 0.00186916 | 0.00077399 | 0 | 0.0013541 |
| *bmp7a* | Secreted Factors | Matrisome-associated | 0 | 0 | 0.00324675 | 0.00067705 |
| *ebi3* | Secreted Factors | Matrisome-associated | 0 | 0 | 0.00324675 | 0.00067705 |
| *sfrp2* | Secreted Factors | Matrisome-associated | 0 | 0 | 0.00324675 | 0.00067705 |
| *vwc2* | Secreted Factors | Matrisome-associated | 0.00093458 | 0.00077399 | 0 | 0.00203114 |
| *wnt4a* | Secreted Factors | Matrisome-associated | 0 | 0.00154799 | 0 | 0.00203114 |
| *tnfb* | Secreted Factors | Matrisome-associated | 0 | 0.00077399 | 0 | 0.00270819 |
| *fgfbp2a* | Secreted Factors | Matrisome-associated | 0 | 0 | 0.00324675 | 0 |
| *wnt3a* | Secreted Factors | Matrisome-associated | 0.00186916 | 0 | 0 | 0.0013541 |
| *shha* | Secreted Factors | Matrisome-associated | 0 | 0.00309598 | 0 | 0 |
| *tnfsf18* | Secreted Factors | Matrisome-associated | 0.00093458 | 0.00077399 | 0 | 0.0013541 |
| *gdf10b* | Secreted Factors | Matrisome-associated | 0 | 0.00232198 | 0 | 0.00067705 |
| *amh* | Secreted Factors | Matrisome-associated | 0 | 0.00154799 | 0 | 0.0013541 |
| *fgfbp2b* | Secreted Factors | Matrisome-associated | 0 | 0.00154799 | 0 | 0.0013541 |
| *angpt2a* | Secreted Factors | Matrisome-associated | 0.00186916 | 0.00077399 | 0 | 0 |
| *gh1* | Secreted Factors | Matrisome-associated | 0.00093458 | 0.00154799 | 0 | 0 |
| *gdf3* | Secreted Factors | Matrisome-associated | 0.00093458 | 0.00077399 | 0 | 0.00067705 |

| | | | | | | |
|---|---|---|---|---|---|---|
| *tnfsf14* | *Secreted Factors* | *Matrisome-associated* | *0.00093458* | *0.00077399* | *0* | *0.00067705* |
| *fgf13b* | *Secreted Factors* | *Matrisome-associated* | *0* | *0.00232198* | *0* | *0* |
| *ccl27a* | *Secreted Factors* | *Matrisome-associated* | *0.00093458* | *0* | *0* | *0.0013541* |
| *cxcl8b.3* | *Secreted Factors* | *Matrisome-associated* | *0.00093458* | *0* | *0* | *0.0013541* |
| *wnt8a* | *Secreted Factors* | *Matrisome-associated* | *0.00093458* | *0* | *0* | *0.0013541* |
| *ccl19a.2* | *Secreted Factors* | *Matrisome-associated* | *0* | *0.00154799* | *0* | *0.00067705* |
| *fgf21* | *Secreted Factors* | *Matrisome-associated* | *0* | *0.00154799* | *0* | *0.00067705* |
| *angptl1b* | *Secreted Factors* | *Matrisome-associated* | *0* | *0.00077399* | *0* | *0.0013541* |
| *hcfc2* | *Secreted Factors* | *Matrisome-associated* | *0* | *0.00077399* | *0* | *0.0013541* |
| *fgf22* | *Secreted Factors* | *Matrisome-associated* | *0* | *0* | *0* | *0.00203114* |
| *wnt7ab* | *Secreted Factors* | *Matrisome-associated* | *0.00093458* | *0.00077399* | *0* | *0* |
| *inhbab* | *Secreted Factors* | *Matrisome-associated* | *0.00093458* | *0* | *0* | *0.00067705* |
| *gdf10a* | *Secreted Factors* | *Matrisome-associated* | *0.00093458* | *0* | *0* | *0.00067705* |
| *s100a1* | *Secreted Factors* | *Matrisome-associated* | *0* | *0.00154799* | *0* | *0* |
| *fgf23* | *Secreted Factors* | *Matrisome-associated* | *0* | *0.00077399* | *0* | *0.00067705* |
| *bmp10* | *Secreted Factors* | *Matrisome-associated* | *0* | *0* | *0* | *0.0013541* |
| *il4* | *Secreted Factors* | *Matrisome-associated* | *0* | *0* | *0* | *0.0013541* |
| *gdf6a* | *Secreted Factors* | *Matrisome-associated* | *0.00093458* | *0* | *0* | *0* |
| *bmp15* | *Secreted Factors* | *Matrisome-associated* | *0* | *0.00077399* | *0* | *0* |
| *fgf8b* | *Secreted Factors* | *Matrisome-associated* | *0* | *0.00077399* | *0* | *0* |
| *il12a* | *Secreted Factors* | *Matrisome-associated* | *0* | *0.00077399* | *0* | *0* |
| *il12ba* | *Secreted Factors* | *Matrisome-associated* | *0* | *0.00077399* | *0* | *0* |
| *ins* | *Secreted Factors* | *Matrisome-associated* | *0* | *0.00077399* | *0* | *0* |
| *wnt5a* | *Secreted Factors* | *Matrisome-associated* | *0* | *0.00077399* | *0* | *0* |
| *wnt8b* | *Secreted Factors* | *Matrisome-associated* | *0* | *0.00077399* | *0* | *0* |
| *cbln2b* | *Secreted Factors* | *Matrisome-associated* | *0* | *0* | *0* | *0.00067705* |
| *gdf2* | *Secreted Factors* | *Matrisome-associated* | *0* | *0* | *0* | *0.00067705* |
| *insl5a* | *Secreted Factors* | *Matrisome-associated* | *0* | *0* | *0* | *0.00067705* |
| *tdgf1* | *Secreted Factors* | *Matrisome-associated* | *0* | *0* | *0* | *0.00067705* |
| *thpo* | *Secreted Factors* | *Matrisome-associated* | *0* | *0* | *0* | *0.00067705* |
| *wnt1* | *Secreted Factors* | *Matrisome-associated* | *0* | *0* | *0* | *0.00067705* |
| *wnt9b* | *Secreted Factors* | *Matrisome-associated* | *0* | *0* | *0* | *0.00067705* |
| *gdf7* | *Secreted Factors* | *Matrisome-associated* | *0* | *0* | *0* | *0.00067705* |

**Table S6. Input gene lists for STRING protein–protein interaction network analysis.** The complete input gene lists used for STRING (v12.0) protein-protein interaction network analysis across the four pairwise comparisons performed: (1) WT_3mt vs. BM_3mt, (2) WT_1yr vs. BM_1yr, (3) WT_3mt vs. WT_1yr, and (4) BM_3mt vs. BM_1yr. For each comparison, the table lists all differentially expressed genes (adjusted $p < 0.05$) submitted to STRING

| **Normal aging** | **Early disease** | **Late disease** | **Disease-associated aging** |
|---|---|---|---|
| *adam8a* | *thbs2b* | *gpc6a* | *ngfb* |
| *angptl4* | *col5a3a* | *lamc1* | *gpc5b* |
| *cilp* | *fgf12b* | *fn1b* | *fstl1b* |
| *coch* | *mfap5* | *ltbp3* | *angpt1* |
| *col11a1b* | *gpc6a* | *col5a2a* | *col5a3a* |
| *col17a1b* | *dcn* | *plxna3* | *dcn* |
| *col27a1b* | *gpc1b* | *ltbp4* | *paplna* |
| *col28a1b* | *igfbp3* | *dcn* | *col1a2* |
| *col5a2a* | *pcsk5b* | *anxa6* | *crim1* |
| *col6a1* | *mfge8b* | *col28a1b* | *thbs2b* |
| *crim1* | *egln1b* | *col5a3a* | *adam10a* |
| *egln1b* | *angptl4* | *col5a2b* | *col5a2a* |
| *fgl2a* | *col5a2b* | *col17a1b* | *cspg5a* |
| *fras1* | *sema4ba* | *bgna* | *sema4ba* |
| *gpc1b* | *bgna* | *muc5.2* | *ltbp4* |
| *gpc5b* | *col1a1b* | *fstl1b* | *postnb* |
| *gpc5c* | *plod2* | *vcanb* | *thbs3a* |
| *gpc6a* | *ntf3* | *lamb1b* | *lamc1* |
| *lamc1* | *crim1* | *paplnb* | *htra1b* |
| *loxl2a* | *lamc3* | *col16a1* | *col6a1* |
| *loxl2b* | *timp2b* | *crim1* | *serpinf1* |
| *mmp14a* | *igfbp5b* | *emilin1b* | *col28a1b* |
| *muc5.2* | *col11a2* | *ntn1a* | *tnfsf10l* |
| *ndnf* | *col1a1a* | *sulf1* | *fn1b* |
| *ngfa* | *pik3ip1* | *col6a1* | *emilin1b* |
| *paplnb* | *loxl2b* | *ndnf* | *fstl1a* |
| *pcsk5b* | *anxa6* | *htra1b* | *sulf1* |
| *plod3* | *col1a2* | *sema6bb* | *lamb1b* |
| *plxnb2b* | *lgi3* | *cilp* | *adam8b* |
| *sema6bb* | *thbs4b* | *fn1a* | *plxna3* |
| *srgn* | *angpt1* | *thbs2b* | *ltbp3* |
| *tgfb1b* | *serpinf1* | *postnb* | *slit3* |
| *thbs2b* | *bmp1a* | *angptl1a* | *adamtsl4* |
| *thbs3a* | *col19a1* | *coch* | *mfap5* |
| | *col28a1b* | *thbs3a* | *sema6bb* |
| | *bcan* | *mmp14a* | *il15* |

| | | | |
|---|---|---|---|
| | col5a2a | ccbe1 | thbs2a |
| | abi3bpb | abi3bpb | igfbp5b |
| | sema3d | mfap5 | f13a1b |
| | anxa4 | colec12 | ccbe1 |
| | sparc | fgf12b | adamts12 |
| | bmp1b | mmp2 | ntn1a |
| | anxa2a | tnfsf10l | col1a1b |
| | postnb | fgf18a | fn1a |
| | fam20ca | adam10a | prg4b |
| | anos1b | col22a1 | gdf10b |
| | frem3 | thbs4b | anxa11a |
| | adamts3 | sema4ba | agrn |
| | lamc1 | lamc2 | ntf3 |
| | pappab | chad | hmcn1 |
| | sdc2 | col10a1a | tll1 |
| | lamb1a | thbs2a | mmp2 |
| | angptl7 | ngfb | igfbp2b |
| | srpx | col27a1b | anxa4 |
| | vegfaa | loxa | vwde |
| | ptn | egln1b | nrg1 |
| | gdnfa | vwa1 | loxa |
| | plxna4 | gpc1b | plod2 |
| | lamb4 | il15 | angptl1a |
| | serpine2 | col11a1b | areg |
| | vwde | sema5a | crlf3 |
| | plod1a | loxl2a | mmp14a |
| | col2a1b | areg | ctsla |
| | col22a1 | tgfb2 | vcanb |
| | p4ha1b | | timp2b |
| | sema6bb | | igf2b |
| | smoc1 | | loxl3b |
| | kazald3 | | il34 |
| | ctsf | | sdc4 |
| | anxa11b | | anxa3b |
| | fstl1b | | anxa5b |
| | col8a2 | | pik3ip1 |
| | cspg5a | | rspo3 |
| | pdgfbb | | timp2a |
| | lgals2a | | sparc |
| | anxa3b | | p4ha1a |
| | hapln1a | | fras1 |
| | angptl2a | | pappab |
| | rspo1 | | bmp4 |
| | f13a1b | | tgfb1a |
| | cst3 | | serpine2 |

| | | | |
|---|---|---|---|
| | *cxcl12b* | | *sdc2* |
| | *gdf10b* | | *mmp14b* |
| | *bglapl* | | *ctsk* |
| | *emid1* | | *serpina10a* |
| | *gas6* | | *csf1a* |
| | *gldn* | | *vwa1* |
| | *nrg1* | | *vcana* |
| | *bmper* | | *lamc3* |
| | *col10a1a* | | *adamts15a* |
| | *igfbp2b* | | *tgfb1b* |
| | *ngfb* | | *fgl2b* |
| | | | *fgf18a* |
| | | | *adamts3* |
| | | | *srpx* |
| | | | *wnt9a* |
| | | | *col8a2* |
| | | | *smoc1* |
| | | | *rspo1* |
| | | | *anxa13* |
| | | | *frem3* |
| | | | *loxl1* |
| | | | *sema3d* |
| | | | *fbln1* |
| | | | *gpc4* |
| | | | *cxcl12a* |
| | | | *col16a1* |
| | | | *hapln1a* |
| | | | *serpinh1b* |
| | | | *angptl2a* |
| | | | *elnb* |
| | | | *gpc6a* |
| | | | *sema3aa* |
| | | | *lgals3b* |
| | | | *vegfaa* |
| | | | *ctsba* |
| | | | *anxa11b* |
| | | | *serpinh1a* |
| | | | *ecm1b* |
| | | | *pdgfbb* |
| | | | *plxna4* |
| | | | *adam8a* |
| | | | *colec12* |
| | | | *bmp7b* |
| | | | *ctsa* |
| | | | *plxnb1a* |

| | | | |
|---|---|---|---|
| | | | *lama5* |
| | | | *angptl4* |
| | | | *sema5a* |
| | | | *adamts5* |
| | | | *pappaa* |
| | | | *kitlga* |
| | | | *sulf2b* |
| | | | *egln1b* |
| | | | *plxnb2b* |
| | | | *gdnfa* |
| | | | *ctsf* |
| | | | *mgp* |
| | | | *lgals2a* |
| | | | *col1a1a* |
| | | | *sparcl1* |
| | | | *col2a1b* |
| | | | *s100a10b* |
| | | | *hcfc1b* |
| | | | *plod3* |
| | | | *clec19a* |
| | | | *gpc1b* |
| | | | *matn3a* |
| | | | *fgl2a* |
| | | | *tgfb2* |
| | | | *cxcl12b* |
| | | | *creld2* |
| | | | *ism1* |
| | | | *srgn* |
| | | | *sema4d* |
| | | | *fgf13a* |
| | | | *gas6* |
| | | | *frzb* |
| | | | *pcsk5b* |
| | | | *ntn1b* |
| | | | *bmp1a* |
| | | | *mmrn2a* |
| | | | *ism2a* |
| | | | *cst3* |
| | | | *lman1* |
| | | | *anos1a* |
| | | | *mfap1* |
| | | | *nrg3b* |
| | | | *hbegfa* |
| | | | *adamtsl5* |
| | | | *anxa13l* |

| | | | |
|---|---|---|---|
| | | | *thbs4a* |
| | | | *tnfsf11* |
| | | | *tgm1l1* |
| | | | *tgfb3* |
| | | | *anxa2a* |
| | | | *nrg2a* |
| | | | *lgals3a* |
| | | | *wnt5b* |
| | | | *lama4* |
| | | | *sema6d* |
| | | | *lgi3* |
| | | | *col4a6* |
| | | | *tinagl1* |
| | | | *cilp* |
| | | | *fgf12b* |
| | | | *eda* |
| | | | *adam17a* |
| | | | *hhip* |
| | | | *f13a1a.1* |
| | | | *sfrp1a* |
| | | | *ptn* |
| | | | *fam20b* |

**Table S7. Differentially expressed genes used for STRING network analysis.** Annotated gene lists used as input for STRING protein-protein interaction network analysis, with upregulated genes highlighted in red and downregulated genes highlighted in blue, for each of the two pairwise comparisons: Normal aging WT_3mt vs. WT_1yr, and Disease associated aging BM_3mt vs. BM_1yr. For each gene, the table reports the gene symbol, log2 fold change, adjusted p-value, and direction of regulation; UP (in salmon) /DOWN (in blue). This table corresponds to the gene sets visualized in the STRING network figures.

| **NORMAL AGING** | | | | | | | |
|---|---|---|---|---|---|---|---|
| **Gene_Symbol** | **Avg_Log2fc** | **P_Val** | **P_Val_Adj** | **Pct.1** | **Pct.2** | **Matrisome_Category** | **Matrisome_Division** |
| *muc5.2* | 3.44046473 | 4.1615E-08 | 0.00103089 | 0.052 | 0.007 | ECM-affiliated | Matrisome-associated |
| *paplnb* | 2.82635588 | 7.4397E-18 | 1.843E-13 | 0.12 | 0.014 | ECM Glycoproteins | Core matrisome |
| *ndnf* | 2.79660854 | 3.3157E-08 | 0.00082137 | 0.055 | 0.007 | ECM Glycoproteins | Core matrisome |
| *plxnb2b* | 2.44868524 | 8.746E-09 | 0.00021666 | 0.068 | 0.011 | ECM-affiliated | Matrisome-associated |
| *serpinh1b* | 2.3174407 | 3.5415E-07 | 0.0087731 | 0.078 | 0.02 | ECM Regulators | Matrisome-associated |
| *chad* | 2.2941082 | 5.4469E-07 | 0.01349309 | 0.065 | 0.014 | Proteoglycans | Core matrisome |
| *col17a1b* | 2.27053973 | 1.0849E-08 | 0.00026875 | 0.075 | 0.014 | Collagens | Core matrisome |
| *loxl2b* | 1.78588213 | 1.005E-11 | 2.4896E-07 | 0.25 | 0.105 | ECM Regulators | Matrisome-associated |
| *slit3* | 1.33902535 | 2.4979E-08 | 0.00061878 | 0.227 | 0.107 | ECM Glycoproteins | Core matrisome |
| *angptl4* | -2.8408214 | 4.125E-12 | 1.0218E-07 | 0.023 | 0.181 | Secreted Factors | Matrisome-associated |
| *sema6bb* | -1.4742454 | 7.5675E-07 | 0.01874633 | 0.045 | 0.154 | ECM-affiliated | Matrisome-associated |
| *col28a1b* | -1.1751332 | 1.0484E-08 | 0.00025972 | 0.13 | 0.293 | Collagens | Core matrisome |
| *gpc6a* | -1.1346319 | 1.4817E-11 | 3.6704E-07 | 0.25 | 0.449 | ECM-affiliated | Matrisome-associated |
| *crim1* | -1.1072733 | 6.6532E-14 | 1.6481E-09 | 0.282 | 0.514 | ECM Glycoproteins | Core matrisome |
| *col6a1* | -1.0675776 | 2.1879E-07 | 0.0054199 | 0.156 | 0.302 | Collagens | Core matrisome |
| **DISEASE-ASSOCIATED AGING** | | | | | | | |
| **Gene_Symbol** | **Avg_Log2fc** | **P_Val** | **P_Val_Adj** | **Pct.1** | **Pct.2** | **Matrisome_Category** | **Matrisome_Division** |
| *fgf7* | 5.12886434 | 1.6296E-08 | 0.00040368 | 0.024 | 0 | Secreted Factors | Matrisome-associated |
| *c1qtnf5* | 4.73767358 | 2.619E-12 | 6.4879E-08 | 0.039 | 0.001 | ECM-affiliated | Matrisome-associated |
| *areg* | 3.34091482 | 1.7976E-12 | 4.4531E-08 | 0.058 | 0.009 | Secreted Factors | Matrisome-associated |
| *fbln1* | 3.29236307 | 1.2278E-07 | 0.00304162 | 0.029 | 0.003 | ECM Glycoproteins | Core matrisome |
| *prg4b* | 3.10949901 | 2.0829E-13 | 5.1597E-09 | 0.061 | 0.009 | Proteoglycans | Core matrisome |
| *il34* | 2.80693624 | 1.7964E-09 | 4.4501E-05 | 0.041 | 0.005 | Secreted Factors | Matrisome-associated |
| *vcana* | 2.57247099 | 1.1111E-06 | 0.02752493 | 0.038 | 0.009 | Proteoglycans | Core matrisome |
| *timp2b* | 2.55900873 | 1.0818E-07 | 0.00267972 | 0.047 | 0.012 | ECM Regulators | Matrisome-associated |
| *rspo3* | 2.4117983 | 1.0626E-08 | 0.00026322 | 0.038 | 0.005 | ECM Glycoproteins | Core matrisome |
| *crlf3* | 2.32150942 | 7.147E-10 | 1.7704E-05 | 0.051 | 0.01 | Secreted Factors | Matrisome-associated |
| *serpina10a* | 2.25172109 | 1.4222E-06 | 0.03522986 | 0.032 | 0.006 | ECM Regulators | Matrisome-associated |
| *emilin1b* | 2.22028737 | 1.3079E-42 | 3.24E-38 | 0.264 | 0.068 | ECM Glycoproteins | Core matrisome |
| *ccbe1* | 2.12308199 | 1.4745E-17 | 3.6526E-13 | 0.112 | 0.028 | Secreted Factors | Matrisome-associated |
| *sulf1* | 2.06053453 | 2.2848E-61 | 5.66E-57 | 0.413 | 0.135 | ECM Regulators | Matrisome-associated |
| *fstl1a* | 2.00857011 | 1.4526E-09 | 3.5983E-05 | 0.058 | 0.014 | Secreted Factors | Matrisome-associated |

| | | | | | | | |
|---|---|---|---|---|---|---|---|
| *p4ha1a* | 1.95893934 | 1.6039E-08 | 0.00039731 | 0.052 | 0.013 | ECM Regulators | Matrisome-associated |
| *tll1* | 1.94443977 | 5.8864E-14 | 1.4582E-09 | 0.095 | 0.026 | ECM Regulators | Matrisome-associated |
| *col5a3a* | 1.9154607 | 3.5968E-50 | 8.9101E-46 | 0.369 | 0.127 | Collagens | Core matrisome |
| *paplna* | 1.88710659 | 1.4815E-12 | 3.67E-08 | 0.085 | 0.023 | ECM Glycoproteins | Core matrisome |
| *adamts12* | 1.87405044 | 2.3213E-08 | 0.00057504 | 0.056 | 0.015 | ECM Regulators | Matrisome-associated |
| *angpt1* | 1.87106658 | 5.4735E-16 | 1.3559E-11 | 0.124 | 0.039 | Secreted Factors | Matrisome-associated |
| *mmp2* | 1.84646461 | 1.4488E-12 | 3.589E-08 | 0.088 | 0.025 | ECM Regulators | Matrisome-associated |
| *htra1b* | 1.8375599 | 6.8862E-75 | 1.7058E-70 | 0.525 | 0.195 | ECM Regulators | Matrisome-associated |
| *bmp7b* | 1.80693624 | 8.9426E-07 | 0.02215259 | 0.053 | 0.018 | Secreted Factors | Matrisome-associated |
| *mfap5* | 1.79715522 | 4.6447E-14 | 1.1506E-09 | 0.093 | 0.024 | ECM Glycoproteins | Core matrisome |
| *adam8b* | 1.73098739 | 2.2149E-12 | 5.4866E-08 | 0.09 | 0.026 | ECM Regulators | Matrisome-associated |
| *sdc4* | 1.72310465 | 4.1414E-10 | 1.0259E-05 | 0.073 | 0.022 | ECM-affiliated | Matrisome-associated |
| *postnb* | 1.70803766 | 1.8052E-30 | 4.4719E-26 | 0.334 | 0.153 | ECM Glycoproteins | Core matrisome |
| *ntn1a* | 1.65346176 | 8.9965E-16 | 2.2286E-11 | 0.156 | 0.06 | ECM Glycoproteins | Core matrisome |
| *il15* | 1.63818179 | 3.6441E-15 | 9.0272E-11 | 0.137 | 0.05 | Secreted Factors | Matrisome-associated |
| *slit3* | 1.63267777 | 1.7915E-21 | 4.4378E-17 | 0.301 | 0.158 | ECM Glycoproteins | Core matrisome |
| *ntf3* | 1.62190435 | 1.0324E-07 | 0.00255757 | 0.056 | 0.017 | Secreted Factors | Matrisome-associated |
| *lamb1b* | 1.57977887 | 1.1061E-26 | 2.7401E-22 | 0.243 | 0.091 | ECM Glycoproteins | Core matrisome |
| *col8a2* | 1.56989705 | 1.2863E-07 | 0.00318641 | 0.086 | 0.037 | Collagens | Core matrisome |
| *cspg5a* | 1.51547343 | 2.3582E-07 | 0.00584175 | 0.073 | 0.029 | ECM-affiliated | Matrisome-associated |
| *hmcn1* | 1.50737596 | 2.1265E-14 | 5.2677E-10 | 0.138 | 0.052 | ECM Glycoproteins | Core matrisome |
| *anxa11a* | 1.50325985 | 1.0835E-09 | 2.6841E-05 | 0.097 | 0.038 | ECM-affiliated | Matrisome-associated |
| *loxl1* | 1.49881395 | 4.5389E-08 | 0.00112437 | 0.077 | 0.029 | ECM Regulators | Matrisome-associated |
| *loxa* | 1.49216287 | 4.1423E-12 | 1.0261E-07 | 0.129 | 0.053 | ECM Regulators | Matrisome-associated |
| *serpinf1* | 1.48765773 | 9.1587E-08 | 0.0022688 | 0.067 | 0.024 | ECM Regulators | Matrisome-associated |
| *sema6bb* | 1.44252482 | 2.9024E-24 | 7.1897E-20 | 0.28 | 0.128 | ECM-affiliated | Matrisome-associated |
| *thbs2a* | 1.41027727 | 7.6659E-17 | 1.899E-12 | 0.165 | 0.063 | ECM Glycoproteins | Core matrisome |
| *igfbp5b* | 1.38943049 | 1.3377E-11 | 3.3138E-07 | 0.199 | 0.108 | ECM Glycoproteins | Core matrisome |
| *adam10a* | 1.38853189 | 5.7462E-21 | 1.4234E-16 | 0.238 | 0.104 | ECM Regulators | Matrisome-associated |
| *thbs2b* | 1.38650549 | 1.0534E-23 | 2.6094E-19 | 0.311 | 0.155 | ECM Glycoproteins | Core matrisome |
| *fn1b* | 1.35800287 | 1.3502E-26 | 3.3447E-22 | 0.379 | 0.201 | ECM Glycoproteins | Core matrisome |
| *gpc4* | 1.34171299 | 5.1497E-07 | 0.01275679 | 0.072 | 0.029 | ECM-affiliated | Matrisome-associated |
| *fgf18a* | 1.33300506 | 1.7128E-08 | 0.00042429 | 0.08 | 0.03 | Secreted Factors | Matrisome-associated |
| *mmp14b* | 1.30650725 | 4.7505E-09 | 0.00011768 | 0.1 | 0.042 | ECM Regulators | Matrisome-associated |
| *ltbp3* | 1.30145853 | 5.5055E-23 | 1.3638E-18 | 0.272 | 0.123 | ECM Glycoproteins | Core matrisome |
| *adamtsl4* | 1.29841212 | 1.2464E-13 | 3.0876E-09 | 0.146 | 0.059 | ECM Regulators | Matrisome-associated |
| *mmp14a* | 1.26838421 | 2.1013E-10 | 5.2054E-06 | 0.108 | 0.043 | ECM Regulators | Matrisome-associated |
| *ltbp4* | 1.18965602 | 5.8161E-20 | 1.4408E-15 | 0.287 | 0.146 | ECM Glycoproteins | Core matrisome |
| *thbs3a* | 1.18308473 | 8.8732E-17 | 2.1981E-12 | 0.211 | 0.096 | ECM Glycoproteins | Core matrisome |
| *col5a2a* | 1.1627392 | 3.1806E-37 | 7.879E-33 | 0.481 | 0.257 | Collagens | Core matrisome |
| *dcn* | 1.15176474 | 1.3112E-19 | 3.2481E-15 | 0.27 | 0.132 | Proteoglycans | Core matrisome |
| *colec12* | 1.14321963 | 1.7109E-07 | 0.00423832 | 0.097 | 0.046 | ECM-affiliated | Matrisome-associated |
| *col28a1b* | 1.13462361 | 4.6258E-21 | 1.1459E-16 | 0.328 | 0.176 | Collagens | Core matrisome |
| *ctsla* | 1.11460365 | 1.8651E-07 | 0.00462022 | 0.096 | 0.045 | ECM Regulators | Matrisome-associated |
| *plod2* | 1.10208501 | 4.5214E-10 | 1.12E-05 | 0.196 | 0.112 | ECM Regulators | Matrisome-associated |

| | | | | | | | |
|---|---|---|---|---|---|---|---|
| *adamts15a* | 1.10198104 | 1.3109E-08 | 0.00032473 | 0.149 | 0.08 | ECM Regulators | Matrisome-associated |
| *tnfsf10l* | 1.09269073 | 1.1055E-10 | 2.7384E-06 | 0.152 | 0.074 | Secreted Factors | Matrisome-associated |
| *adamts3* | 1.08957521 | 3.4928E-07 | 0.00865225 | 0.097 | 0.046 | ECM Regulators | Matrisome-associated |
| *wnt9a* | 1.064734 | 4.0798E-07 | 0.01010644 | 0.093 | 0.044 | Secreted Factors | Matrisome-associated |
| *fn1a* | 1.03474216 | 1.1961E-12 | 2.963E-08 | 0.221 | 0.119 | ECM Glycoproteins | Core matrisome |
| *fstl1b* | 0.94642558 | 5.1627E-24 | 1.2789E-19 | 0.356 | 0.185 | Secreted Factors | Matrisome-associated |
| *timp2a* | 0.94443977 | 3.0992E-07 | 0.00767732 | 0.103 | 0.05 | ECM Regulators | Matrisome-associated |
| *loxl3b* | 0.88444561 | 1.3404E-08 | 0.00033205 | 0.175 | 0.101 | ECM Regulators | Matrisome-associated |
| *angptl1a* | 0.85584584 | 1.6529E-09 | 4.0946E-05 | 0.247 | 0.158 | Secreted Factors | Matrisome-associated |
| *cxcl12a* | 0.84690265 | 1.488E-06 | 0.03686195 | 0.102 | 0.053 | Secreted Factors | Matrisome-associated |
| *plxna3* | 0.83535532 | 1.0113E-15 | 2.5053E-11 | 0.352 | 0.217 | ECM-affiliated | Matrisome-associated |
| *sema4ba* | 0.81713201 | 3.0475E-11 | 7.5492E-07 | 0.441 | 0.334 | ECM-affiliated | Matrisome-associated |
| *crim1* | 0.72500439 | 2.1648E-18 | 5.3626E-14 | 0.434 | 0.28 | ECM Glycoproteins | Core matrisome |
| *sparc* | 0.71551322 | 4.0877E-07 | 0.01012601 | 0.187 | 0.118 | ECM Glycoproteins | Core matrisome |
| *col6a1* | 0.65395538 | 2.3878E-12 | 5.9151E-08 | 0.443 | 0.32 | Collagens | Core matrisome |
| *vcanb* | 0.52063206 | 5.0775E-08 | 0.00125781 | 0.242 | 0.158 | Proteoglycans | Core matrisome |
| *col1a2* | 0.44661932 | 7.2797E-09 | 0.00018033 | 0.35 | 0.249 | Collagens | Core matrisome |
| *sparcl1* | -2.9744235 | 1.0813E-10 | 2.6787E-06 | 0.005 | 0.04 | ECM Glycoproteins | Core matrisome |
| *gpc5b* | -2.9448047 | 1.1799E-35 | 2.9228E-31 | 0.022 | 0.153 | ECM-affiliated | Matrisome-associated |
| *fgf12a* | -2.8935035 | 2.9856E-08 | 0.0007396 | 0.005 | 0.034 | Secreted Factors | Matrisome-associated |
| *frem3* | -2.0397673 | 9.1012E-22 | 2.2545E-17 | 0.039 | 0.142 | ECM-affiliated | Matrisome-associated |
| *fras1* | -1.9074427 | 1.2497E-24 | 3.0957E-20 | 0.064 | 0.19 | ECM Glycoproteins | Core matrisome |
| *egln1b* | -1.5726524 | 5.5073E-20 | 1.3643E-15 | 0.063 | 0.173 | ECM Regulators | Matrisome-associated |
| *sema5a* | -1.4885196 | 3.5008E-14 | 8.6723E-10 | 0.042 | 0.119 | ECM-affiliated Proteins | Matrisome-associated |
| *sema3aa* | -1.3209812 | 8.6452E-18 | 2.1416E-13 | 0.091 | 0.204 | ECM-affiliated | Matrisome-associated |
| *abi3bpb* | -0.8436598 | 2.8628E-07 | 0.00709172 | 0.186 | 0.269 | ECM Glycoproteins | Core matrisome |
| *gpc5c* | -0.8124598 | 7.6806E-07 | 0.0190265 | 0.08 | 0.138 | ECM-affiliated | Matrisome-associated |
| *loxl2b* | -0.7649701 | 1.483E-09 | 3.6738E-05 | 0.146 | 0.235 | ECM Regulators | Matrisome-associated |
| *slit2* | -0.6889004 | 3.5735E-08 | 0.00088523 | 0.114 | 0.19 | ECM Glycoproteins | Core matrisome |
